# Multilayer diffusion networks as a tool to assess the structure and functioning of fine grain sub-specific plant-pollinator networks

**DOI:** 10.1101/2021.04.23.441120

**Authors:** Alfonso Allen-Perkins, María Hurtado, David García-Callejas, Oscar Godoy, Ignasi Bartomeus

**Author notes:** Repo: https://github.com/RadicalCommEcol/Multi_motifs.

## Abstract

Interaction networks are a widely used tool to understand the dynamics of plant-pollinator ecological communities. However, while most mutualistic networks have been defined at the species level, ecological processes such as pollination take place at different scales, including the individual or patch levels. Yet, current approaches studying fine-grain sub-specific plant-pollinator networks only account for interactions among nodes belonging to a single plant species due to the conceptual and mathematical limitations of modeling simultaneously several plant species each composed of several nodes. Here, we introduce a multilayer diffusion network framework that allows modeling simple diffusion processes between nodes pertaining to the same or different layers (i.e. species). It is designed to depict from the network structure the potential conspecific and heterospecific pollen flows among plant individuals or patches. This potential pollen flow is modeled as a transport-like system, in which pollen grain movements are represented as random-walkers that diffuse on an ensemble of bipartite layers of conspecific plants and their shared pollinators. We exemplify this physical conceptualization using a dataset of nine fine-grain sub-specific plant-pollinator networks from a Mediterranean grassland of annual plants, where plant nodes represent groups of conspecifics within patches of 1m^2^. The diffusion networks show pollinators effectively connecting sets of patches of the same and different plant species, forming a modular structure. Interestingly, different properties of the network structure, such as the conspecific pollen arrival probability and the number of conspecific subgraphs in which plants are embedded, are critical for the seed production of different plant species. We provide a simple but robust set of metrics to calculate potential pollen flow and scale down network ecology to functioning properties at the individual or patch level, where most ecological processes take place, hence moving forward the description and interpretation of species-rich communities across scales.

## Introduction

Network approaches have been increasingly used in ecology to interpret the complex structure of interactions among species (Bascompte *et al*., 2003; Martinez, 1991; Thebault & Fontaine, 2010). By understanding ecological communities as networks in which species are linked through pairwise interactions, prior theoretical and empirical work has shown that network architecture strongly influences species dynamics (Bartomeus *et al*., 2021; Bascompte & Jordano, 2013; Brose *et al*., 2006), with quantifiable effects on community properties such as local stability (Allesina & Tang, 2012), robustness to extinctions (Memmott *et al*., 2004), or the impact of invasive species or climate change on local assemblages (Memmott *et al*., 2007). However, despite the recent knowledge gained on the architecture of ecological networks, we are still lacking a solid foundation linking network architecture with higher-level ecological functions, such as primary productivity, decomposition, or pollination (Bergamo *et al*., 2020; Magrach *et al*., 2021; Poisot *et al*., 2013; Thompson *et al*., 2012).

To quantify the structure of the ecological network and its links to ecological functioning, most approaches are based on a species-level perspective. However, this view ignores the fact that several ecological processes occur at smaller scales and depend on individual variation (Bolnick *et al*., 2011; Dall *et al*., 2012; Herrera, 2017; Sih *et al*., 2012; Wolf & Weissing, 2012). For example, plant pollination is a key function mediated by insects that transport pollen from one plant individual to another. Collapsing all plant individuals from a given species into a single aggregate (i.e. node) assumes that there is no intraspecific variation in traits such as floral display, plant height, or flower morphology, and ultimately on pollinator visitation rates. This simplification clashes with several empirical examples showing that within-species variation can result in different assemblages of pollinators (Arroyo-Correa *et al*., 2021; Gómez *et al*., 2020). Pollinators can discern and respond to intraspecific flower variation at the patch level (Herrera, 2017), influencing the vegetative (e.g., growth rate, carbon assimilation) and reproductive (e.g., fecundity) performance of individuals (Herrera, 2017). Not surprisingly, in the last decade the number of studies that analyze individual-based networks has grown from 6 studies in 2010 to 82 in 2020 (reviewed by Guimarães (2020)). Nevertheless, the study of individual-based networks is still in its infancy (Guimarães, 2020).

For instance, there are only a dozen studies exploring individual-based plant-insect pollination networks. This is due to both theoretical and methodological reasons. Empirically, the sampling effort required to resolve all species interactions into individual ones is labor-intensive, which partly explains the scarcity of empirical studies of individual-based networks, either for one species (Dáttilo *et al*., 2015; Gómez & Perfectti, 2012; Gómez *et al*., 2011; Kuppler *et al*., 2017, 2016; Soares *et al*., 2020; Tur *et al*., 2013; Valverde *et al*., 2015) or more species (Dupont *et al*., 2014, 2010; Pornon *et al*., 2017). A valid way to overcome the labor-intensive work of documenting interactions of small individuals of annual plants is to at least capture fine-grain sub-specific variability at the patch level (e.g. *∼* 1m^2^ for annual plants, Ghazoul, 2006; Maia *et al*., 2019). More concerning is the fact that we lack robust methods to resolve the complexity of integrating individual or patch-level nodes belonging to multiple species. As a consequence, most studies on individual-based networks focus on the individuals of one species (usually a plant species linked by its pollinators (Gómez & Perfectti, 2012; Gómez *et al*., 2011; Soares *et al*., 2020; Valverde *et al*., 2015), or a single pollinator (Dupont *et al*., 2014)), ignoring individuals of other species of the plant community.

We present a set of tools derived from multilayer networks and diffusion dynamics (hereafter referred to as “multilayer diffusion networks”) to understand the general patterns of interactions between conspecific and heterospecific groups of plant individuals belonging to multiple species. Multilayer networks are networks with nodes and links distributed across *layers*. Each layer represents aspects or features of the nodes or the links that belong to the layer, and the links that connect nodes in different layers (i.e. interlinks) provide information on the processes or interdependencies operating between such layers (see the reviews of Boccaletti *et al*. (2014) and Kivela *et al*. (2014) for further details about the multilayer paradigm and its formal definitions). In ecology, multilayer networks typically depict species interactions through time and space, or across types of interactions or behaviors (Crestani *et al*., 2019; Garćıa-Callejas *et al*., 2018; Hutchinson *et al*., 2018; Pilosof *et al*., 2017), but its application using one layer per species within a community has not yet been explored. Here we take advantage of the flexibility of the multilayer formalism to conceptualize fine-grain sub-specific pollination networks as diffusion systems (Aleta *et al*., 2017; Domenico *et al*., 2014). Diffusion systems on top of networks model how information flows across the nodes based on the pattern of interactions of the network (i.e., its topology). From a plant-centric point of view, the advantage of using diffusion networks is that the information flow can be analogous to the pollen flow dynamics, and hence, have a clear ecological interpretation. Note that rather than mechanistically modeling pollen flow, we aim to use generalizable and well-studied information flow metrics as proxies of pollen diffusion.

In a nutshell, when combining multilayer and diffusion networks, we propose to analyze the potential pollen flow mediated by pollinators in an ensemble of bipartite layers of conspecific plant individual nodes coupled through pollinator species nodes. Specifically, conspecific plant individuals and their pollinator species are the nodes of each layer, whereas interlayer links account for shared pollinators across layers. Interestingly, these interlayer links can account for plants’ and pollinators’ phenologies, or pollinator efficiencies (see Figs. 1(a) and 1(e)). Note that in our framework, individual nodes can refer both to individuals that belong to a given species for large-size plants, but they can also represent groups of small plants belonging to the same patch (i.e. within a 1 m^2^). We often will be forced to collapse pollinators to the species level, given that they are mobile species, for which we often lack individual-based information (Arroyo-Correa *et al*., 2021; Gómez & Perfectti, 2012), but the framework works with individual pollinator nodes as well when this information is available. This framework allows us to study in detail the structure and effects of interactions from the node level (Masuda *et al*., 2017), to the entire network level (Farage *et al*., 2021; Rosvall *et al*., 2009), including meso-scale descriptors such as subgraphs (Milo *et al*., 2002; Porter *et al*., 2009; Simmons *et al*., 2018).

**Fig. 1:**
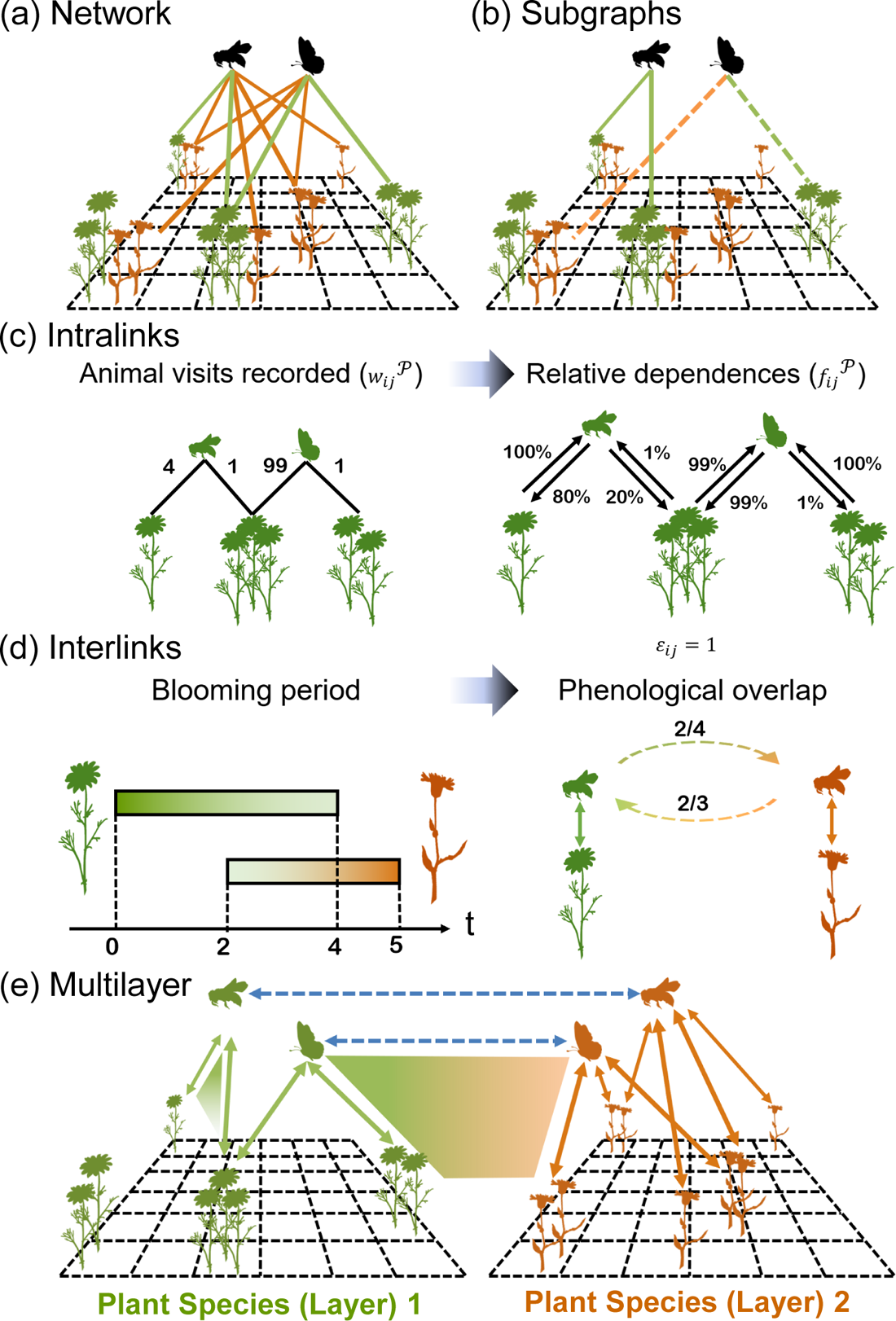
(a) An example of a visitation network for a 6 *×* 6 plot with two plant species of annual plants (modeled as patch individual nodes) and two pollinator species, respectively. For the sake of readability, weights *w^P^* have been omitted. (b) In this panel, we highlight examples of homospecific (solid lines) and heterospecific (dotted lines) subgraphs, respectively, in Fig. 1(a). (c) An example of a visitation network depicting the raw number of visits observed, and the resulting flow arrangement taking values from 0 to 1 for plant species 1 (green) in Fig. 1(a). (d) An example of the estimation of interlinks for plant species 1 and 2 (green and orange, respectively) in Fig. 1(a). (e) Resulting multilayer network for the arrangement in Fig. 1(a). Continuous double-arrows represent directed intralayer links, whereas dashed ones show directed interlinks. Shaded areas highlight the multilayer analogs of the subgraphs in Fig. 1(b). As can be seen, layers consist of conspecific focal individual patches of plants and their interactions with their pollinators, and shared insects connect layers depending on their interspecific floral visits and on the phenological overlap among plant species.

Here, we first present the analytical details of the proposed framework by testing the behavior of the proposed metrics in *in silico* networks, and then we apply it to a particular case study using nine fine-scale sampled field plots in a Mediterranean grassland ecosystem, which further allows linking these metrics to individual plant reproduction. We hypothesize that nodes having a position that maximizes the potential arrival of conspecific information flow (i.e., pollen) will show a comparatively higher seed production per flower. Conversely, nodes whose information flow mainly comes from other layers may increase heterospecific pollen arrival and decrease their seed set, as has been previously shown experimentally (Morales & Traveset, 2009; Moreira-Herńandez & Muchhala, 2019).

## Analytical framework

### Multilayer networks

The multilayer networks we present here are ensembles of bipartite layers of conspecific plant nodes and their pollinators, that are coupled through shared pollinator nodes, where connections encode the information flow between pairs of nodes mediated by pollinators (see Figs. 1(a), 1(e) and 3 for an example). Here, each layer contains conspecific fine-grain sub-specific plant nodes (such as patch- or individual-based nodes), the pollinator nodes (species-based nodes) of that plant species, and their interactions (*intralinks*). To model the information flow between two layers mediated by the pollinator movements and species phenologies, if a pollinator node is present in both layers, we connect the shared pollinator nodes through *interlinks* (see below the definition of intra- and interlinks, respectively, in subsection “Multilayer diffusion networks”). By doing so, we assume that pollinators can easily reach all plant nodes within the system, that is, their foraging range largely exceeds the system size. These multilayer networks fit in the ‘diagonally coupled’ category described in (Pilosof *et al*., 2017), where networks with different interaction types (defined here by the plant species involved) are connected through shared species. In addition, they can also be described as node-colored multilayers, where each layer or plant species represents a color (see (Kivela *et al*., 2014)). Note that the same pollinator species node can appear in different layers, but are effectively treated as a single species thanks to the interlink connections that always connect all instances of each pollinator species node.

### Diffusion dynamics

To unveil potential pollen flows, we model the information spreading across the multilayer nodes using random walk dynamics, a process that has been applied thoroughly on top of networks (de Arruda *et al*., 2018) with widespread applications such as the analysis of disease spreading (Bestehorn *et al*., 2021), among other diffusion processes on networks (Masuda *et al*., 2017).

The usual (discrete-time) random walk is a random sequence of nodes that describes how a *walker* (a proxy for information exchange *sensu* Rosvall & Bergstrom (2008), from e.g. a rumor in human social networks to a pollen grain in plant-pollinator ones) moves from node to node. The sequence is generated as follows: given a starting node *i* (in our case, a plant node), denoted as ‘origin of the walk’ (or origin of the information, our proxy for pollen grains), at each discrete time step *t*, the random walker moves to one of its nearest out-neighbors, that is, to one of the nodes that have a connection from that where the walker is (Masuda *et al*., 2017). If the network is weighted, then the underlying assumption is that the larger the weight of a connection, the more likely it will be used to transmit information (see Eq. (24) in Masuda *et al*. (2017)). Note that weights can be directional and asymmetric, and hence the probability of going from one node *i* to *j* can be different from the probability of going from *j* to *i* (see weight calculation in subsection “Multilayer assembly”). In the case of multilayer networks, because of their peculiar interconnected structure, random walkers can also move from one layer to another, provided interlinks are connecting those layers. For instance, in the case of transportation systems where layers may represent underground, bus, and railway networks, random walk dynamics can be used to mimic trips among locations making use of more than one transport mode (Domenico *et al*., 2014).

In this work, for the sake of simplicity, we suppose that transition probabilities among nodes (i.e., the weights of the links) are asymmetric and do not vary with time and, consequently, do not depend on the previous positions of the walker. This also means that the structure of our multilayers is static. Our main hypothesis is that the relevant abiotic and biotic factors that drive the asymptotic potential pollen flow among plant and pollinator nodes during a given period of time are encoded in that fixed multilayer structure. Nevertheless, more complex scenarios where the transitions of the walker (from node to node) depend on its past positions and some sequences of positions are favored and others are discouraged (or even forbidden) could be also considered (see the section devoted to memory networks in Masuda *et al*. (2017) for an example).

### Multilayer assembly

To model the diffusion of random walkers on top of our multilayers, the weight of the link from node *i* to node *j* represents the probability that pollen (i.e. information) in *i* flows to *j* through that link, regardless of whether it is an intra- or interlink. Thus, those weights should take values between 0 and 1. How to extract potential information flow probabilities from observed field data depends on the data collected.

Here we propose a way to quantify those probabilities (i.e., the network structure) from visitation frequency data. Flower visitation data is a common and sensible proxy for pollination and ultimately gene flow (Page *et al*., 2021; Vázquez *et al*., 2005). Visitation rates can be measured at the individual or patch level, since visitation decisions depend mainly on the patch context (Ghazoul, 2006; Seifan *et al*., 2014).

To turn visitation data into probabilities of exchanging pollen between a pair of nodes, we can calculate the weight of intralinks (i.e., the connections within a given layer) as follows. Firstly, for each plant-species *P*, we can build a weighted, undirected bipartite graph, denoted here as *visitation network* (VN-*P*), from the plant–pollinator interaction matrix (see left-hand side of Fig. 1(c) for an example), where link weight represents the total number of visits that the plant node *i* of species *P* received from a pollinator species *j*. Secondly, assuming that, in a given visitation network VN-*P*, the larger the weight of a connection *w^P^* between nodes *i* and *j*, the more likely that the interaction will transfer pollen between such nodes, we derived a directed *flow network* from the undirected visitation graph by calculating the fraction of plant *i* pollen that is passed to pollinator *j* (or alternatively the fraction of pollen carried by pollinator *i* that is passed to plant *j*) as follows:

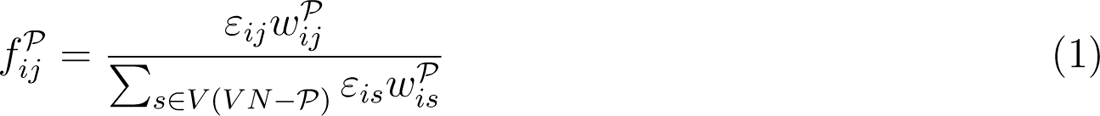

where *f^P^_ij_ ∈* [0, 1] represents the fraction of information (or pollen grains) allocated in *i* that flows to *j* in the layer *P*, *ε_ij_∈* [0, 1] is the efficiency of the exchanges from node *i* to *j*, V(VN-*P*) denotes the nodes in the visitation network of *P*, and *w^P^* = 0 is the link weight, the relative number of visits between species *i* and *j*. Equation 1, in practice, models a situation in which e.g. the information flow from a pollinator species to plant individuals is divided according to the relative number of visits to these plant individuals (see Fig. 1(c) for an example). These flow networks are directed, and we further assume that the information flow between two plant nodes does not decay with distance, that is, visitor species effectively link all plant nodes they interact with. On the one hand, this assumption is justified if we consider that all the pollinators can move within the system under analysis easily. On the other hand, it is a requirement to meet the condition that the transition probabilities of walkers are constant. Nevertheless, as mentioned above, this approach can be generalized to impose information flow decay with distance, for instance, by making *ε_ij_*dependent on the position of plants along the random walk (Hod, 2003).

Finally, to couple the different layers and, thus, assemble the multilayer network, we quantified the interconnections between insects of the same species that visit several plant species (layers). These are modeled as interlinks, representing how easily the potential pollen flows from one plant species to another due to the phenological overlap between them and the interspecific floral visits of pollinators. To do so, we proceeded as follows: if a given pollinator *x* was present in layers *P* and *L*, we created one interlink from the pollinator node *x* in layer *P* (i.e., *x^P^*) to *x* in layer *L* (i.e., *x^L^*) with weight *f ^P→L^*, and another interlayer connection from *L* to *P* with weight *f ^L→P^*, where *P* and *L* denote respectively two plant species, and *f ^P→L^* is equal to the phenological overlap of *P* and *L* divided by the total duration of *P*’s phenology (see Fig. 1(d) for an example of these calculations). As in the case of intralinks, this approach for estimating interlinks can be easily extended to account for alternative definitions (where -for example-the weight of interlinks is also weighted by the number of shared visits), or even additional restrictions to diffusion between layers. For example, by multiplying *f ^P→L^* by *π^P→L^*, the probability that pollinator species *x* deposits pollen of plant species *P* in *L*, we could consider that differences in flower morphology and stigma position can lead to specialization in pollen location along a pollinator’s body parts (Armbruster *et al*., 1994), which, in turn, may prevent high levels of heterospecific pollen deposition. To conclude, note that, according to previous definitions, interlayer links are directed (i.e., *f ^P→L^* and *f ^L→P^* can be different), and the most important thing: if a given layer is isolated (i.e., it has no interlinks), its plant individuals will only receive conspecific pollen mediated by pollinators, and no heterospecific pollen.

### Network metrics

Once we created the multilayer arrangements, we assessed how the position of plant nodes within that structure shapes the potential pollen flow that reaches them and potentially affects plant reproduction. In particular, we estimated a series of diffusion network metrics related to the entire network- and node-level structural patterns, and we complemented it with subgraph-level descriptors. We considered subgraph-level processes because it has been recently suggested that they may be particularly important for analyzing the functional consequences of community structure: at that scale, the local patterns of realized direct interactions can reveal the different mechanisms through which individual- and species-based nodes can indirectly influence each other (Simmons *et al*., 2018) and it has been shown that some sub-graphs are over-represented in plant-pollinator networks (i.e. form distinctive motifs) (Lanuza *et al*., 2023). For the network-level structure, we calculated (i) the modular partitions that better describe network information flows (subsection “Modularity”). For the node-level structure, we obtained (ii) the plant node’s probability of receiving information from conspecific and heterospecific plant nodes, respectively (subsection “Potential arrival of information”). Finally, for each plot of our study system, we characterized subgraph-level structure through (iii) the 3-node subgraph characterizations, denoted as homospecific subgraphs and heterospecific subgraphs (subsection “Subgraph analysis”).

#### Modularity

Network modularity measures how closely connected nodes are divided into modules or compartments (Porter *et al*., 2009). Our hypothesis is that plant nodes belonging to modules where their species is very abundant will have greater reproductive success than plant nodes belonging to modules where that species is rare. Since our multilayers encode potential pollen flows, we defined modules as groups of nodes that best capture flows within and across layers, rather than clusters with a high internal density of links (Domenico *et al*., 2015). Among the different strategies to detect modules, the one that best matches the above definition is that of Infomap (R package `infomapecology` v0.1.2.3, Farage *et al*. (2021)) (Supplementary material Appendix 2). Infomap uses the map equation to measure the minimum number of bits (or code length, L) that are needed to describe the movement of a random walker in and between the modules of a given network partition *M* (Farage *et al*., 2021; Rosvall *et al*., 2009). Then, the algorithm finds the partition that requires the least amount of information to describe modular flows (see further details on the algorithm in (Bohlin *et al*., 2014; Farage *et al*., 2021)). Importantly, although the nodes in our multilayers belong to just one module, in the case of insect nodes, their “physical” counterparts (that is, the observed insect species) can be assigned to different modules in different layers (Farage *et al*., 2021). In addition, note that we approximated pollen flows between plants and pollinators from the number of visits observed in the field, so we applied the *directed* flow model in Infomap (Farage *et al*., 2021).

Once the optimal module partition for a multilayer is found, we can test if the total number of modules *m* and the code length *L* can be explained at random. To do so, we compared those values of *L* and *m* with the distributions obtained from two null models that constrain, on the one hand, the degree distribution of plants and pollinators, along with the empirical phenology, and, on the other hand, the degree distribution of pollinators (see details in Supplementary material Appendix 3).

#### Potential arrival of pollen

The random walk dynamics on top of multilayer networks allow estimating a proxy of plant nodes’ probabilities of receiving pollen from other plant nodes, both conspecific and heterospecific. Here, we hypothesize that the probability of receiving different pollen from conspecific and heterospecific plant nodes can explain the node’s functionality (e.g. pollen deposition in stigmas, and ultimately seed production per fruit). We followed the methodology presented by Masuda *et al*. (2017) for characterizing discrete-time random walk dynamics and calculated Π^Coupled^, the probability of finding the random walker (a pollen grain) in plant node *i* at discrete time *t*, when the origin of the walker (that is, the plant node that “released” the pollen grain at time *t* = 0) was the plant node *j*. That probability can be extracted from the so-called random-walk’s master equation, which expresses the probability of finding the random walker in *i* at time *t* in terms of the probabilities of being at any node at time *t −* 1:

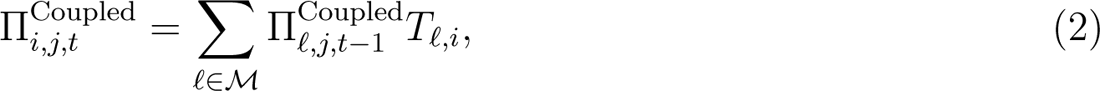

where *M* represents the set of all nodes in the multilayer network, and *T_ℓ,i_*is the probability that pollen in node *ℓ* flows to node *i*. Note that, according to definitions introduced above, *T_ℓ,i_* = *f ^P^* (see Eq. 1) when nodes *ℓ* and *i* are connected by an intralink, and *T_ℓ,i_* = *f ^L→P^* otherwise (that is, when nodes *ℓ* and *i* represent an observed pollinator species that visits plant species *L* and *P*, respectively).

We also estimated the probability of finding a pollen grain “released” by the plant node *j* in plant node *i* at time *t*, when the layers are uncoupled, Π^Uncoupled^. This can be obtained from Eq. 2 by setting the weights of interlinks to zero (i.e., by making *T_ℓ,i_* = 0 when nodes *ℓ* and *i* represent pollinator species) and, consequently, there is no information flow from heterospecific plant nodes. The estimation of Π^Uncoupled^ models a scenario where different factors prevent the heterospecific pollen deposition, such as the foraging behavior of pollinators (e.g. their floral constancy (Jakobsson *et al*., 2008)) or the specialization in pollen location along a pollinator’s body parts, induced by differences in flower morphology and stigma position (Armbruster *et al*., 1994).

From the above probabilities, we derived the following metrics for information flow:

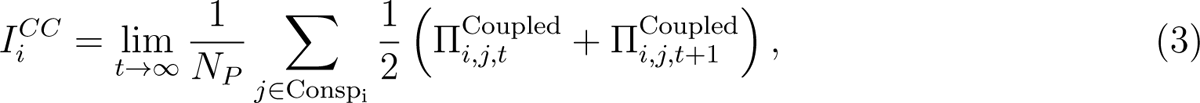

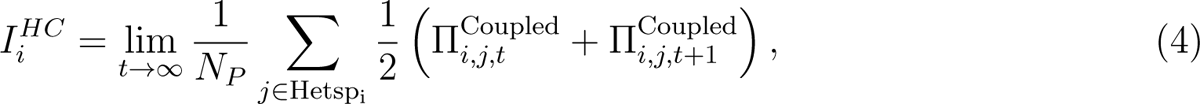

and

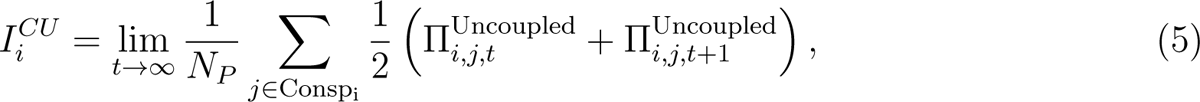

where *I^CC^_i_* and *I^HC^_i_* represent the average asymptotic probability of receiving pollen from con-specific and heterospecific plant nodes, respectively, when layers are coupled; *I^CU^* is the average asymptotic probability of receiving pollen from conspecific nodes when layers are uncoupled; *N_P_* refers to the total number of plant nodes; and Consp_i_ and Hetsp_i_ represent the set of conspecific and heterospecific plant nodes of node *i*, respectively. Lastly, the average asymptotic probability of receiving pollen from heterospecific plant nodes when layers are uncoupled, *I^HU^*, is zero by definition. All these probabilities represent the plant’s total probability of receiving conspecific/heterospecific pollen (from all potential sources), when the layers are coupled/uncoupled, and when time tends to infinity.

Note that we are adapting well-characterized dynamics (Eq. 2) from unipartite networks (Masuda *et al*., 2017) to bipartite multilayer networks. The methods are equally valid for uni- and bipartite networks, with the only difference being that here we need to estimate an average limiting distribution (Eq. 3-5) instead of a stationary probability distribution (Supplementary material Appendix 1).

#### Subgraph analysis

To complement the network- and node-level descriptions of the multilayer structures, we incorporate additional details on the local architecture of direct and indirect interactions among nodes by decomposing the networks into subgraphs, the basic building blocks of communities (Milo *et al*., 2002). Subgraph analyses are well developed for food webs (Cirtwill *et al*., 2018), mutualistic (Simmons *et al*., 2020, 2018), and competitive networks (Godoy *et al*., 2017), but have seldom been applied to multilayer networks. In our case, we only analyzed the number of undirected triplets present in our visitation networks, that is, the pattern of connections of undirected path graphs with length two (see examples in Fig. 1(b)). For the sake of simplicity, hereafter we only use the term “subgraph” to refer to triplets. However, even such simple three-node subgraphs can be differentiated according to the layers involved (see examples in Fig. 1(e)). Considering subgraphs containing one or two layers is interesting because it perfectly maps onto the effects of indirect conspecific and heterospecific information flows on focal individual’s per capita seed production.

In addition, it moves forward the bipartite subgraph descriptions from species-based networks, in which all the subgraph nodes are heterospecific (see Simmons *et al*. (2020, 2018)), to fine-grain sub-specific arrangements with conspecific and heterospecific nodes.

To conduct our subgraph analyses, we introduced a novel subgraph categorization according to the plant species involved: If all the plant nodes belong to the same species, the subgraph was referred to as *homospecific subgraph*; otherwise, the subgraph was classified as *heterospecific* (see Fig. 1(b)). According to previous studies on species-based mutualistic bipartite networks, such subgraphs can represent indirect competitive (Mitchell *et al*., 2009; Ye *et al*., 2013) or facilitative interactions (Carvalheiro *et al*., 2014; Ghazoul, 2006). For example, in Fig. 1(a) plant individuals may be involved in an exploitative competition for finite pollinator resources, or interference competition through interspecific pollen deposition (Flanagan *et al*., 2010; Mitchell *et al*., 2009; Ye *et al*., 2013). Conversely, facilitative effects may occur when the presence of one focal individual increases pollinator visits to a coflowering individual. Unfortunately, without further information on species performance, the observed patterns can not be attributed to a specific process.

To conduct our analysis, in each plot, we focused on the number of homospecific and heterospecific subgraphs of a target plant node, which inform about how many other conspecific and heterospecific plant nodes (i.e., *acting plant nodes*) contribute to the diet of its pollinators, respectively. Since the acting plant nodes of a given target plant may potentially not belong to the same module as the target plant, homospecific and heterospecific subgraphs complement the modularity analyses described above. Furthermore, to incorporate species’ phenological overlap in our subgraph analysis, we measured species’ overlap at a weekly resolution, which was the temporal resolution of our visitation data. We calculated homospecific and heterospecific subgraphs from the weekly bipartite networks observed (see Fig. 1(a) for an example of such networks). Heterospecific subgraphs only appear when different plant species that share pollinators are co-flowering (i.e. both present in a given weekly sampling). Then, we calculated the total number of homospecific and heterospecific subgraphs for a given focal individual by summing up the number of subgraphs containing that individual throughout all sampling weeks. Note that, if the temporal resolution of the sampling allows it, different temporal slices can be selected to calculate phenological overlap among nodes (e.g. daily, monthly). In addition, we tested if the number of subgraphs is different from those of a random model where phenology is preserved (see Supplementary material Appendix 4).

All the above calculations were performed in R v4.3.0 (R Core Team, 2021). A guide to ensemble multilayer diffusion networks and calculating all metrics is included in the associated code (section “Code availability”).

### Sensitivity analyses of network metrics

We used computer simulations to show the generality of our approach and understand how the proposed metrics change in response to changes in network structure. To do so, first, we built 400 synthetic but realistic multilayer diffusion networks from a model based on the following input variables: (i) the total number of plant nodes, (ii) the total number of pollinator nodes, (iii) the total number of intralinks, (iv) the total number of interlinks, and (v) the number of layers. To ensure that the intralayer degree distributions of those *in-silico* multilayers are realistic (Fig. A6.1), we constrained their average intralayer link density (i.e., the total number of intralinks divided by the total number of plant nodes) and set it equal to two, and also established the total number of pollinator nodes, interlinks and layers as 25, 20 and five, respectively, which are similar to the average numbers of those features in our experimental case study (22.89 *±* 12.09 pollinator species, 16.00 *±* 14.21 interlinks, and 4.40 *±* 2.13 layers, see “Community structure in Empirical Results” below). Note that fixing the intralayer link density along with the number of pollinator nodes is equivalent to constraining the intralayer connectance sensu (Dormann *et al*., 2009), which is given by the number of intralinks divided by the product of the number of plant nodes and pollinator nodes. Thus, the model parameterization presented here only depends on one variable, namely: the total number of intralinks. Then, to keep the intralink density constant (and equal to two links per plant node), we adjust the number of plant individuals to half that of intralinks. We set values between 100 and 200 intralinks, which again are similar to those found in our empirical multilayers (176.89 *±* 89.23 intralinks). Importantly, with no additional constraints, the modular features of the studied in-silico networks are realistic (Fig. A6.2). Finally, we analyzed the dependence of the subgraph- and node-level metrics on the number of intralinks in those 400 in-silico multilayers with constant link density. We also interpreted how the number of intralinks and plant nodes affect those metrics when they vary one at a time. All the simulations and analyses were conducted in R 4.3.0 (R Core Team, 2021) (for further details on the random multilayer generator see Supplementary material Appendix 5). Note that systems where those constraints are relaxed may exhibit different behaviors, but exploring how contrasting network topologies affect diffusion dynamics is beyond the scope of this paper.

In a nutshell, the results from these simulations show that subgraphs-level metrics are more sensitive than node-level ones to variations in link richness (Fig. 2). Conversely, those trends suggest that node-level metrics depend more on the position of intralinks than subgraph-metrics. Overall, the number of both homospecific and heterospecific subgraphs increases with an increase in the number of intralinks, when keeping intralink density constant. However, the number of homospecific subgraphs grows faster than that of heterospecific subgraphs, as expected, since the latter is more constrained by the presence of pollinator species without interlinks. These trends for multilayer networks with constant intralink density suggest that subgraph metrics are mainly driven by the number of intralinks (Suppl. subsection 6.1, top panels in Fig. A6.5) instead of by the number of plant nodes (Suppl. subsection 6.2, top panels in Fig. A6.8). In addition, they indicate that our generator of synthetic multilayers works as expected, and can be used to test and prove the generality of our approach.

**Fig. 2:**
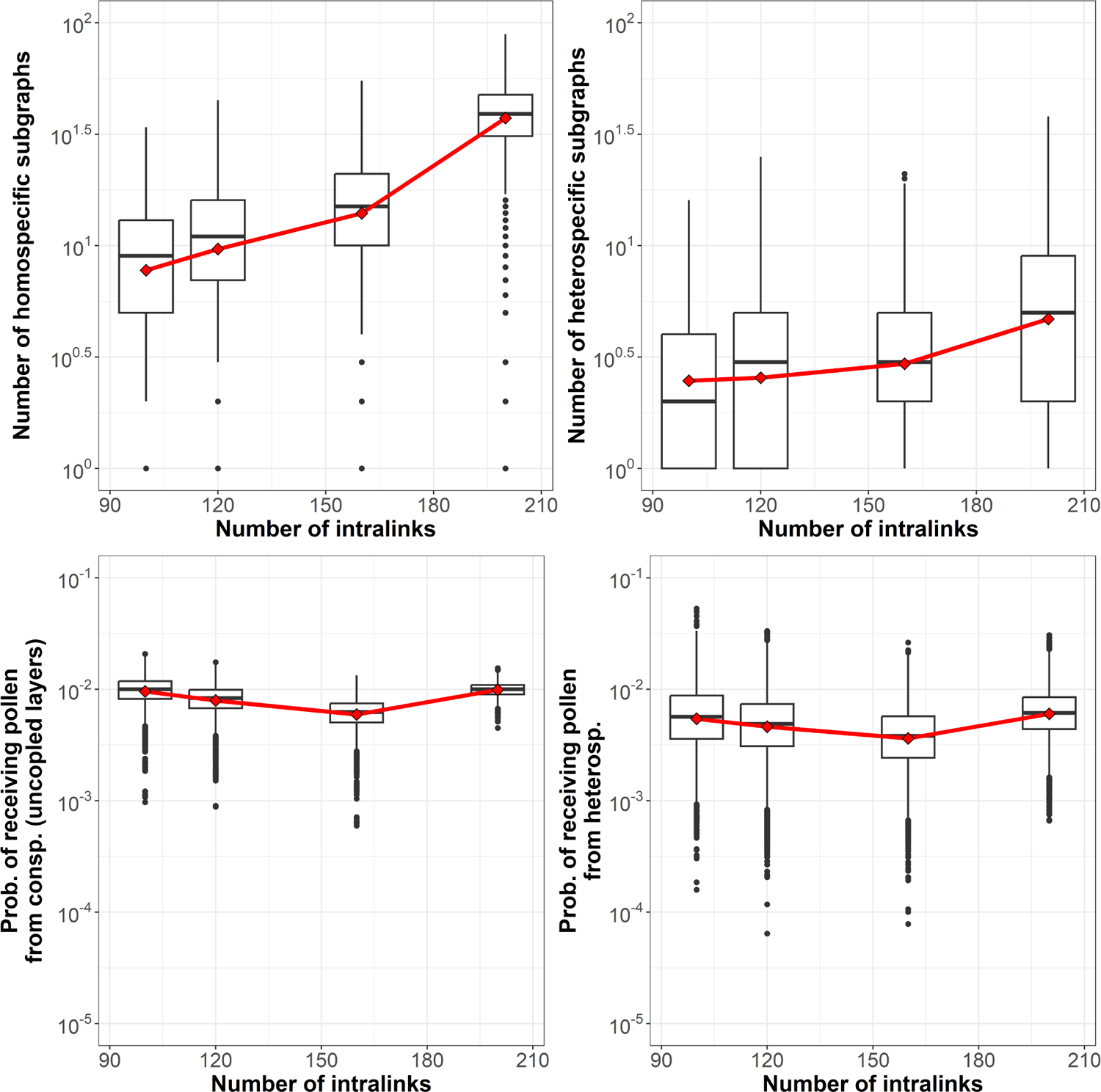
Top panels: Dependence of the total number of homospecific subgraphs (left panel), and total number of heterospecific subgraphs (right panel) on the number of intralinks. Bottom panels: Dependence of the probability of receiving pollen from conspecific plant nodes when the layers are uncoupled (left panel), and of the probability of receiving pollen from heterospecific plant nodes (right panel) on the total number of intralinks. Mean values are shown with red diamonds. Red lines are a guide to the eye to highlight the mean value trends.

Regarding the plant node’s total probability of receiving conspecific and heterospecific pollen (from all potential sources), when layers are both coupled and uncoupled, our results show that these metrics have a small but significant variation with the number of intralink richness (Figs. 2, A6.3 and A6.4) in our highly constrained *in-silico* multilayers. Given that increasing intralink density does not impact the average probability of receiving conspecific pollen (Suppl. sub section 5.1, left bottom panel in Fig. A6.5 and A6.6), these results suggest that the number of plant nodes (Suppl. subsection 6.2, bottom panels in Fig. A6.8) and the specific pattern of connections of each node are driving the nodes’ probability of receiving pollen, and not the number of connections in the community. Interestingly, the total probability of heterospecific arrival is of the same magnitude as that of conspecific pollen, even when layers are uncoupled. That highlights the great impact that interlinks exert in multilayer networks, even when not all pollinator species visit all plant species (layers). In addition, given that link density is fixed along with the number of pollinators and interlinks, scenarios where pollinator species visit on average very few or a lot of plant nodes (that is, scenarios with a very low or very high number of intralinks) slightly improve conspecific and heterospecific pollen diffusion. That confirms an expected result for our constrained *in-silico* multilayers: those structural features that increase the probability of conspecific pollen transfer also increase the probability of heterospecific pollen transfer, when plant species are coupled by pollinators. Thus, these simulations confirm that the selected metrics are useful for understanding the topology of fine-grain sub-specific plant-pollinator networks.

### Case study: A Mediterranean annual plant community

To test our framework, we use an observational study in Caracoles Ranch, a 2680 ha saline grassland located within Doñana NP, south-western Spain (37*^◦^*04’01.0” N, 6*^◦^*19’16.2” W). This grassland is dominated by annual herbs of small size creating patches of monospecific stands as well as more diverse spatial configurations (Hurtado *et al*., 2023). We located nine square plots of side 8.5 m distributed in three sub-areas with different flooding regimes to capture variability in the dynamics of annual plants (details in Lanuza *et al*. (2018)). The average distance between these three locations was 100 m and the average distance between plots within each location was 57 m. Each plot was divided into 36 square subplots of side 1 m with aisles of 0.5 m in between to allow access to subplots where measurements were taken (324 subplots in total). Despite some pollinators can potentially fly between plots, we treat them as independent for practical reasons, and because when resources are abundant and pollinators small, like in our case, most pollinators forage in less than 100 m radius (Kendall *et al*., 2022).

During spring 2020, we sampled groups of plant individuals and their pollinators across the nine established plots (see Hurtado *et al*. (2023) for details). Specifically, we observed 23 co-occurring annual plant species (Supplementary material Appendix 7 table A7.1). We surveyed weekly (weather permitting) the identities of pollinators and their number of visits to each plant species within each subplot from February to June, covering the phenology of all plant species (see Supplementary material Appendix 8). Visits were only considered when the floral visitor (hereafter pollinator) touched the reproductive organs of the plant. All subplots within a plot were simultaneously sampled for 30 min each week by walking slowly between the subplots, but pollinator observations were recorded at the subplot level (1 m^2^). The plot survey order was randomized between weeks to avoid sampling effects. All pollinators were either identified during the survey or were net-collected for their posterior identification at the lab. Overall, this procedure rendered approximately 54 hours of pollinators sampled over 19 weeks, and an estimated sampling coverage of subplot plant–pollinator interactions in each plot network of 90% (see Appendix 9 for further details in Supplementary material). Thus, our sampling procedure is appropriate to quantitatively characterize plant-pollinator interactions at fine-grain sub-specific scales in our system. Pollinators were identified to the species (42.34%) or morphospecies (57.56%) level (Supplementary material Appendix 10 table A11). In addition, to analyze the composition of the visitor spectrum of each plant species and characterize how they connect the module of multilayers (i.e., their specialization roles), we also assigned each insect to one of the following eleven taxonomic groups: small beetles, big beetles, butterflies, flies, flower beetles, house flies, hoverflies, humbleflies, small flies, solitary bees, and wasps (Supplementary material Appendix 10 table A10.1). Insects only visited the flowers of 10 of the 23 species present (Supplementary material Appendix 8 Fig. A8.1), and most of these visits were concentrated on three Asteraceae species, namely: *Chamaemelum fuscatum*, *Leontodon maroccanus* and *Pulicaria paludosa*.

To obtain a measure of pollination function per subplot, seed production per floral unit was sampled in one individual plant of each species present per subplot and plot. We collected one fruit per individual per subplot and plot from the three most attractive species for insects (*C. fuscatum*, *L. maroccanus* and *P. paludosa*) which represent 95.65% of all insect visits and, thus, are the focus of the seed production analysis (see below; Supplementary material Appendix 7 table A7.1). We cleaned the collected fruits and counted the number of developed seeds. *C. fuscatum* and *P. paludosa* depend on pollinators to maximize seed set (selfing rates are low), while *L. maroccanus* is mostly self-compatible (sefling rates are high), but pollinators can be important for cross-pollination (Hurtado *et al*., 2023).

### Multilayer assembly

As mentioned above, we have reproductive-success-related data per subplot, and we measured the frequency of visitation of the sampled plant individuals at the patch level. To build our multilayer network, we aggregate the information on the number of visits for each species in each subplot in a single node for which we can associate their visitation patterns and their total seed production. This is a fair assumption, as the main driver of visitor attraction for small annual co-flowering plants is their context, that is, the flower density of the patch and its spatial distribution (Ghazoul, 2006; Seifan *et al*., 2014). From visitation frequency data, we followed the framework presented above to build nine multilayer diffusion networks (one per plot) to describe (i) the potential information flow between individual plant patches (i.e. our fine-grain sub-specific nodes) mediated by pollinators (i.e. we assume that pollinators connecting two plant nodes are transferring pollen), and (ii) the effect of flowering phenology and pollinator efficiency on such potential flow.

Finally, we assumed that the taxonomic group of pollinator nodes affects their information exchanges with plant nodes with beetles being less efficient than flies, and flies less efficient than bees. This information is depicted in the efficiencies *ε_ij_*that are needed to compute the intralinks (Supplementary material Appendix 11 table A11.1).

### Network metrics

Following our analytical framework, we extracted structural information from the multilayer diffusion networks at network-, node- and subgraph-levels. For the network-level structure, we used Infomap to estimate the optimal modular partitions and their specialization roles (sensu Olesen *et al*. (2007)). We also tested if the observed module partitions are different from those from a random expectation. For the node-level structure, we obtained the plant node’s average asymptotic probabilities of receiving information from conspecific and heterospecific plant nodes, both when layers are coupled and uncoupled. Finally, for the subgraph-level structure, we calculated the total number of homospecific and heterospecific subgraphs, considering weekly phenological overlap among species. We tested if the number of subgraphs is different from those of a random model where phenology is preserved.

#### Seed production analysis

One key aspect of this multilayer approach is that we can study the impact of direct and indirect interactions between plants and pollinators at different structural scales on flower seed production. We explore that relationship by fitting generalized linear mixed models (GLMMs) to the individuals of the three species that attracted more pollinators (*C. fuscatum*, *L. maroccanus*, and *P. paludosa*, an early, middle, and late phenology species, respectively). Since our response variable (seed production per fruit) represents count data and we detected overdispersion issues in our exploratory analysis, we used negative binomial models in our inferences with a log link function and a constant zero-inflated term (see Eq. A17.1 in Appendix 17). As explanatory variables, all models included the following descriptors: (i) for the node-level structure, the plant nodes’ probabilities of receiving conspecific (i.e., when layers are uncoupled) and heterospecific pollen from other plant nodes (when layers are coupled); and (ii) subgraph-level structural descriptors, i.e. the number of homospecific and heterospecific subgraphs. We did not consider the probabilities of receiving conspecific pollen (when layers are coupled) because it exhibits a high positive correlation with the values obtained when the layers are uncoupled (Spearman’s *ρ* = 0.66, p-value *<* 0.05). We also used “plot” as a random intercept to account for multiple individuals of the same plant species measured at each plot. In addition, we tested how the results of our models vary if we add the total number of visits that an individual patch of plants received to the previous explanatory variables. To keep the regression variables on similar scales, all explanatory variables were centered and scaled during the analysis.

Our analyses were conducted in R, with the glmmTMB v1.1.3 package (Brooks *et al*., 2017). We found no collinearity among explanatory variables when we checked their variance inflation factors with the R-package performance v0.8.0 (Lüdecke *et al*., 2020). We also checked model assumptions with the R-package DHARMa v0.4.5 (Hartig, 2020).

### Empirical Results

We recorded 1,794 insect visits in our system from February to June 2020. The distributions of visits among the taxonomic groups of pollinators, plant species, and plots, respectively, showed a high variation (Supplementary materials Appendix 12 Fig. A12.1). Most interactions involved *L. maroccanus* (1,337 observations) and, to a lesser extent, *C. fuscatum* (268) or *P. paludosa* (111). Small flies and flower beetles were the most abundant insects visiting *C. fuscatum*, while small beetles and flower beetles were the most abundant for *L. maroccanus*, and solitary bees for *P. paludosa*. The number of visits per plot ranges between 360 and 44, with a mean value of 199.33 *±* 113.18 insect visits.

#### Community structure

Plot multilayers obtained from the above field observations contain on average 4.4 *±* 2.13 plant species (layers), 71.89 *±* 37.34 nodes (of which 49.00 *±* 25.29 are plant nodes, and 22.89 *±* 12.09 pollinator nodes), 176.89 *±* 89.23 intralinks, and 16.00 *±* 14.21 interlinks (see Fig. 3 for an example of the multilayer networks analyzed and Supplementary material Appendix 13 for the graphs of the remaining multilayers).

**Fig. 3:**
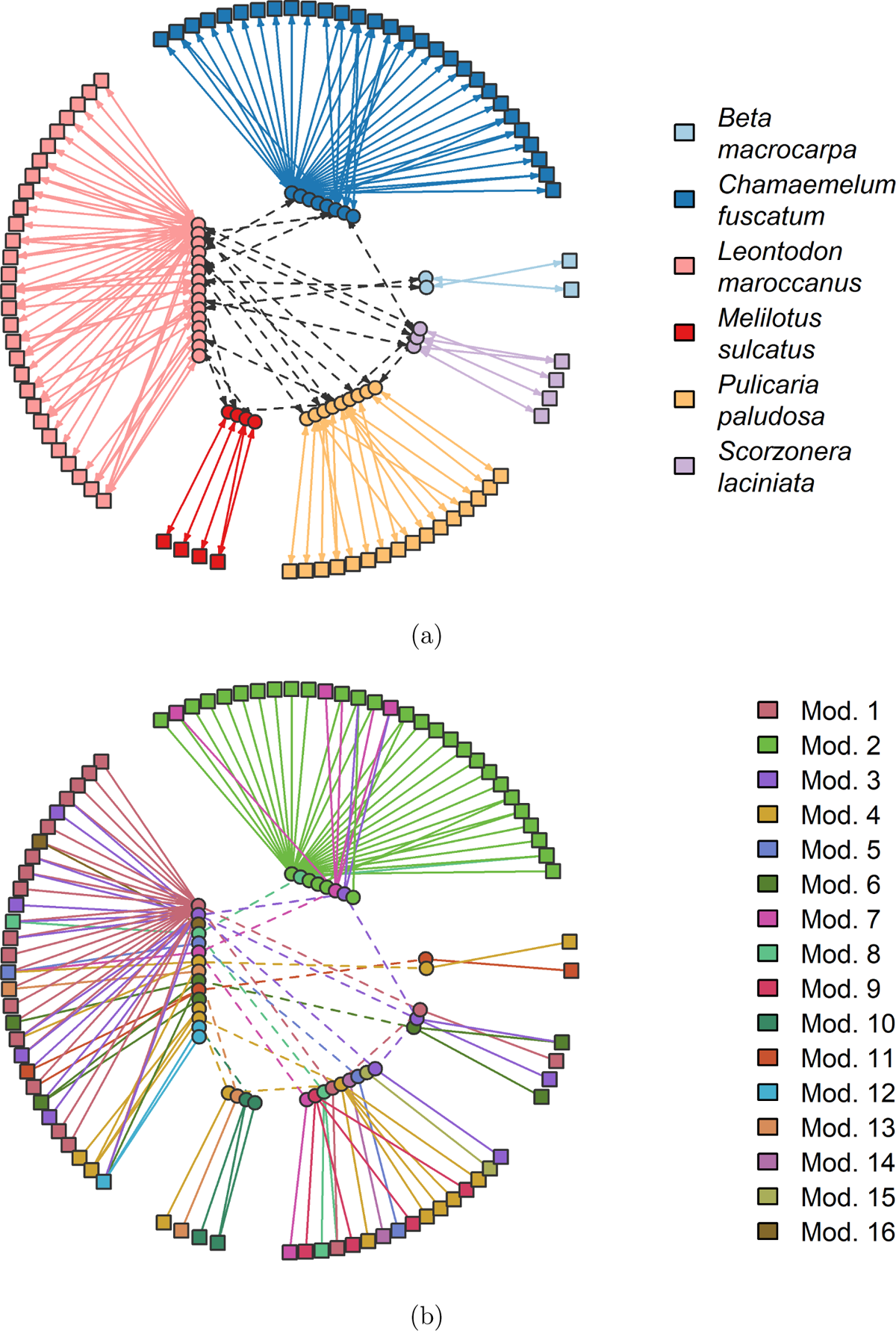
(a) Resulting multilayer for plot 8. Each color refers to a layer: *B. macrocarpa* (light blue), *C. fuscatum* (blue), *L. maroccanus* (pink), *M. sulcatus* (red), *P. paludosa* (yellow), and *S. laciniata* (malve). We represent plant individual patch-level nodes and insect species with square and circular nodes, and their interactions by continuous links. Note that a pollinator group can be visiting two species of plants, i.e. being present in two layers, and this is denoted by black dashed edges. (b) Modules identified with Infomap (R package infomapecology v0.1.2.3, Farage *et al*. (2021)) for the multilayer in Fig. 3(a). Here nodes with the same color belong to the same module. Note that as interlinks are constrained by phenology, this is also reflected in the modularity analysis, with module 2 separating *C. fuscatum*, a species with a distinctive early phenology.

We found on average 10.78 modules per plot using Infomap, being (3, 16) the corresponding 95% confidence interval (CI). One-third of the modules contained at least two plant species (Figs. 3(b) and 4), Supplementary materials Appendix 12 Fig. A12.3), which shows that insect visitors are effectively linking plant layers. Overall, the modules contained a small number of patches and are dominated by a single plant species (i.e. within the same layer), but many contain more than one plant species. The average size of modules is 6.67 nodes with CI (2, 36) and the average number of plant individuals per module is 4.55 with CI (1, 26). As expected, the larger the plot richness, the larger the number of modules (the coefficient of determination between both variables is 0.85, with p-value *<* 0.05). Our results also suggest that modules reflect phenological constraints. For instance, *C. fuscatum* plant individuals in plot two do not share modules with those of *P. paludosa* (Figs. 3(b) and 4) because their respective phenologies do not overlap and phenology is accounted for when depicting the network of interactions (Supplementary material Appendix 8 Fig. A8.1).

**Fig. 4:**
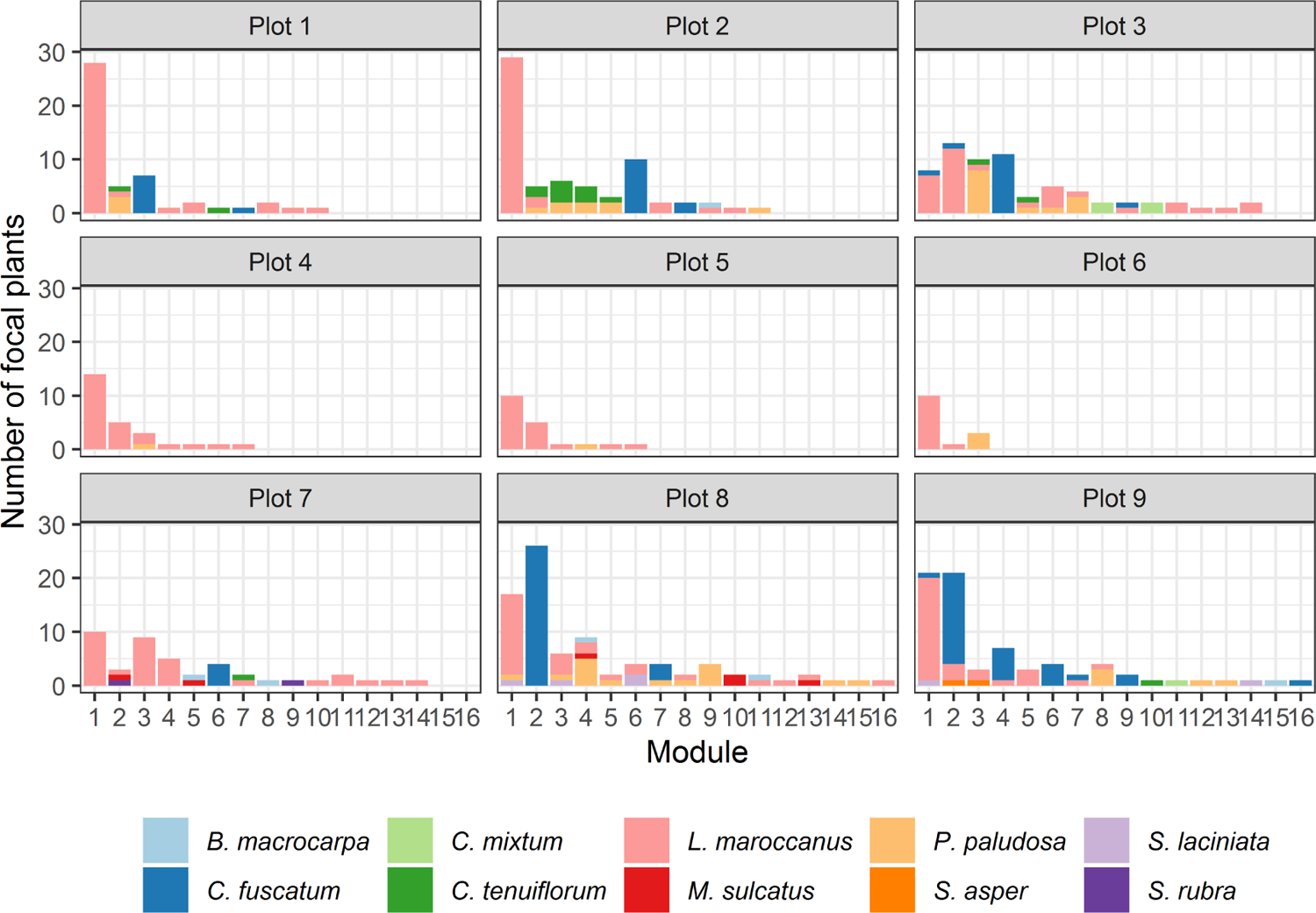
Number of plant nodes per module, plant species, and plot.

Results for insect species, which represent on average one-third of the nodes per module (2.12 pollinator nodes per module with CI (1, 6)), are very similar to those of plants (Supplementary materials Appendix 12 Fig. A12.3). Additional details on module features, such as specialization roles, can be found in Supplementary materials (Appendix 14).

Finally, we tested if the observed module partitions are different from a random expectation in which either the degree distribution of plants and pollinators is constrained, along with the empirical phenology, or the degree distribution of pollinators (Supplementary material Appendix 3). According to our results, the number of modules is not significantly different from that of null models in most plots (see Supplementary material Appendix 15). However, the optimal partitions of the observed multilayers provide a significantly better description of the potential pollen flows within the whole system (or exhibit more modular regularities, i.e., the observed values for the map equation, *L_Observed_*, are smaller) than those estimated for random graphs. In other words, while the number of modules obtained is consistent with a random expectation, their composition is significantly different from it.

### Potential arrival of pollen

Overall, we found that the total probability of receiving pollen from conspecifics and heterospecific plant nodes is highly and positively correlated with the total number of visits a plant node received (Spearman’s *ρ >* 0.50, p-value *<* 0.05). Contrary to the results for synthetic multilayers (in which the intralinks were randomly added, see subsection “Sensitivity analyses of network metrics”), the most visited plant species (*C. fuscatum*, *L. maroccanus* and *P. Paludosa*) have a higher probability of receiving pollen from conspecific nodes than from heterospecific nodes, especially when the probability of conspecific pollen is calculated with the layers uncoupled (paired Wilcoxon-test: p-value *<* 0.05) (Fig. 5). Uncoupled layers assume pollinator behavior avoids loss of pollen to heterospecific plants, but as reported above, the probability of coupled and uncoupled conspecific pollen arrival is highly correlated (Spearman’s *ρ >* 0.66, p-value *<* 0.05).

**Fig. 5:**
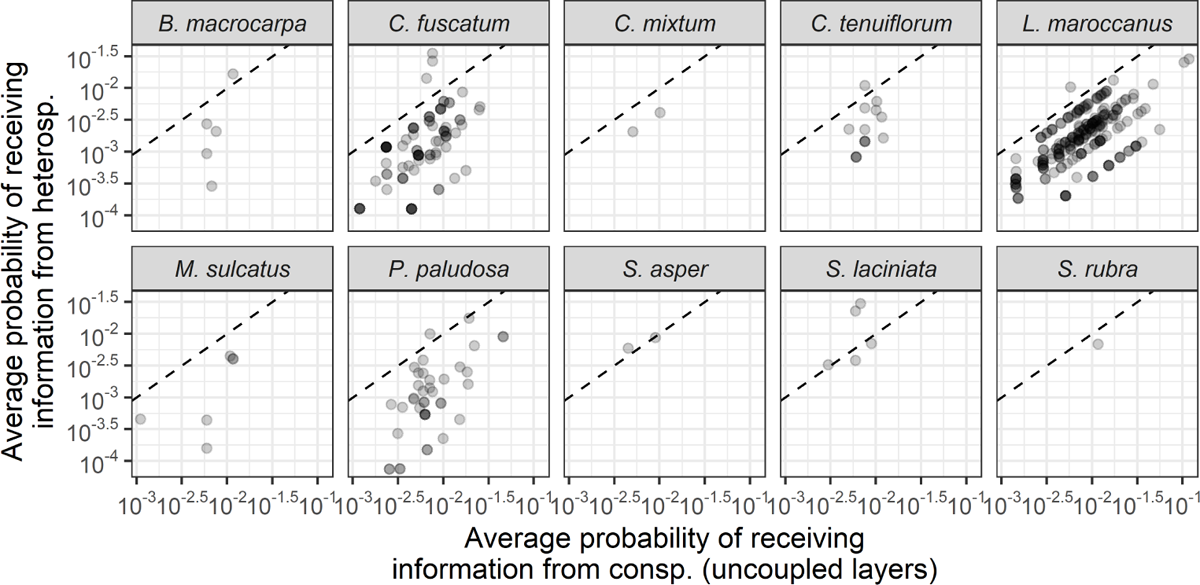
Probability of receiving pollen from conspecific and heterospecific plant nodes per focal individual patch and plant species, pooling the data for all nine plot networks. Points show results for individual patches of plants. Dashed lines locate those plant individual patches whose probabilities of receiving pollen from conspecific and heterospecific plant nodes are equal.

### Homospecific and heterospecific subgraphs

In our communities, the number of homospecific subgraphs is highly correlated with the total number of visits (Spearman’s *ρ* = 0.73, p-value *<* 0.05), whereas the number of heterospecific subgraphs is weak (Spearman’s *ρ* = 0.11, p-value *<* 0.05). Homospecific subgraphs are more abundant than heterospecific ones in *C. fuscatum* and *L. maroccanus* nodes (paired Wilcoxon-test: p-value *<* 0.05 for both plant species), whereas both metrics are comparable in most focal individuals of *P. paludosa* (paired Wilcoxon-test: p-value = 0.14), and, in the case of other plant species, heterospecific subgraphs predominate over homospecific subgraphs (paired Wilcoxon-test: p-value *<* 0.05 only for C. tenuiflorum) (Fig. 6). That means that nodes belonging to the plant species with the largest number of visits and the smallest number of pollinators (such as *C. fuscatum* and *L. maroccanus*) tend to share more weekly visits of the same insects within their conspecific nodes than with heterospecific nodes, whereas those nodes of rarer plant species share all their pollinators with heterospecific nodes (such as those of *L. maroccanus*), along the sampling weeks (Supplementary material Appendix 8 Fig. A8.2). Nevertheless, our null model analysis (based on random networks where the degree distribution of plants and pollinators is constrained, along with the empirical phenology (Supplementary material Appendix 3)) indicates that, overall, the observed number of homospecific subgraphs of *C. fuscatum* and *L. maroccanus* nodes tend to be significantly smaller than those obtained from a random expectation (Supplementary materials Appendix 16 Fig. A16.1). Remarkably, in the case of *C. fuscatum*, we observe that most of its homospecific subgraphs are non-significant. This is due to the reduced phenological overlap among *C. fuscatum* and other plant species (see Fig. A8.1). Finally, results for heterospecific subgraphs show that, in general, their number is larger than expected (Supplementary materials Appendix 16 Fig. A16.2).

**Fig. 6:**
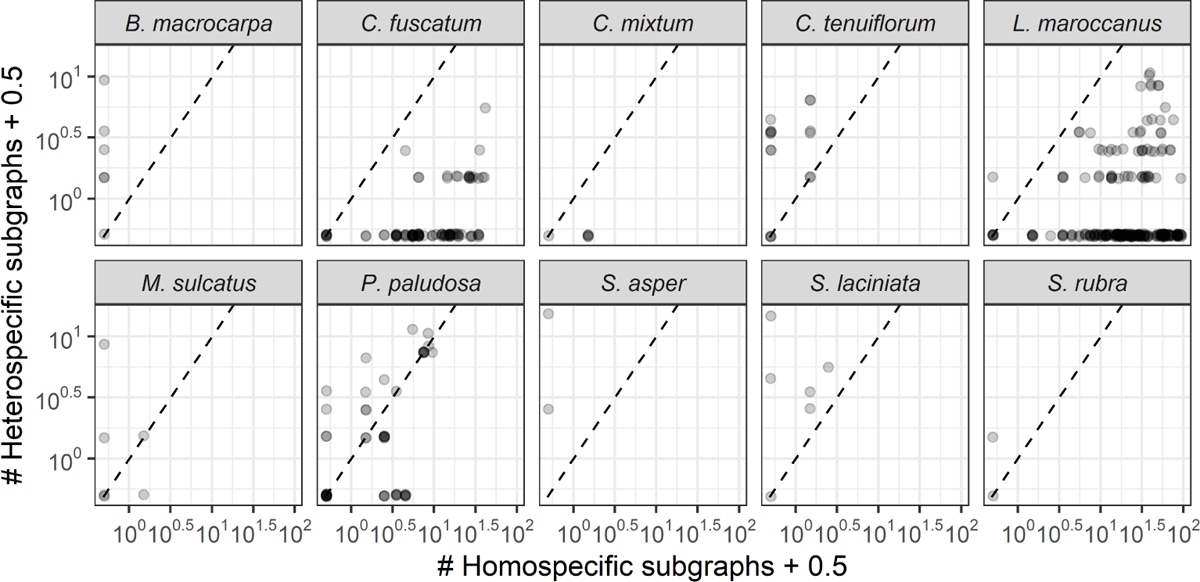
Number of different homospecific and heterospecific subgraphs per focal individual patch and plant species, pooling the data for all nine plot networks. Points show results for plant nodes, i.e., individual patches of plants. Dashed lines locate those plant individual patches whose numbers of homospecific and heterospecific subgraphs are equal.

### Seed production analysis

Results for the GLMMs of the three plant species that attracted the most pollinators (*C. fuscatum*, *L. maroccanus* and *P. paludosa*) show that, overall, different metrics are relevant for understanding the seed production patterns of the different species (see Fig. 7). At the node level, the probability of receiving pollen from conspecific patch individuals (when there is no pollen exchange with heterospecific nodes) is significant for *C. fuscatum* and *P. paludosa*, whereas the probability of receiving information from heterospecifics is not significant for any of the species studied. The trends for *C. fuscatum* and *P. paludosa* are those we hypothesized: the seed set of their plant nodes is larger when the probability of receiving pollen from conspecific plant nodes increases.

**Fig. 7:**
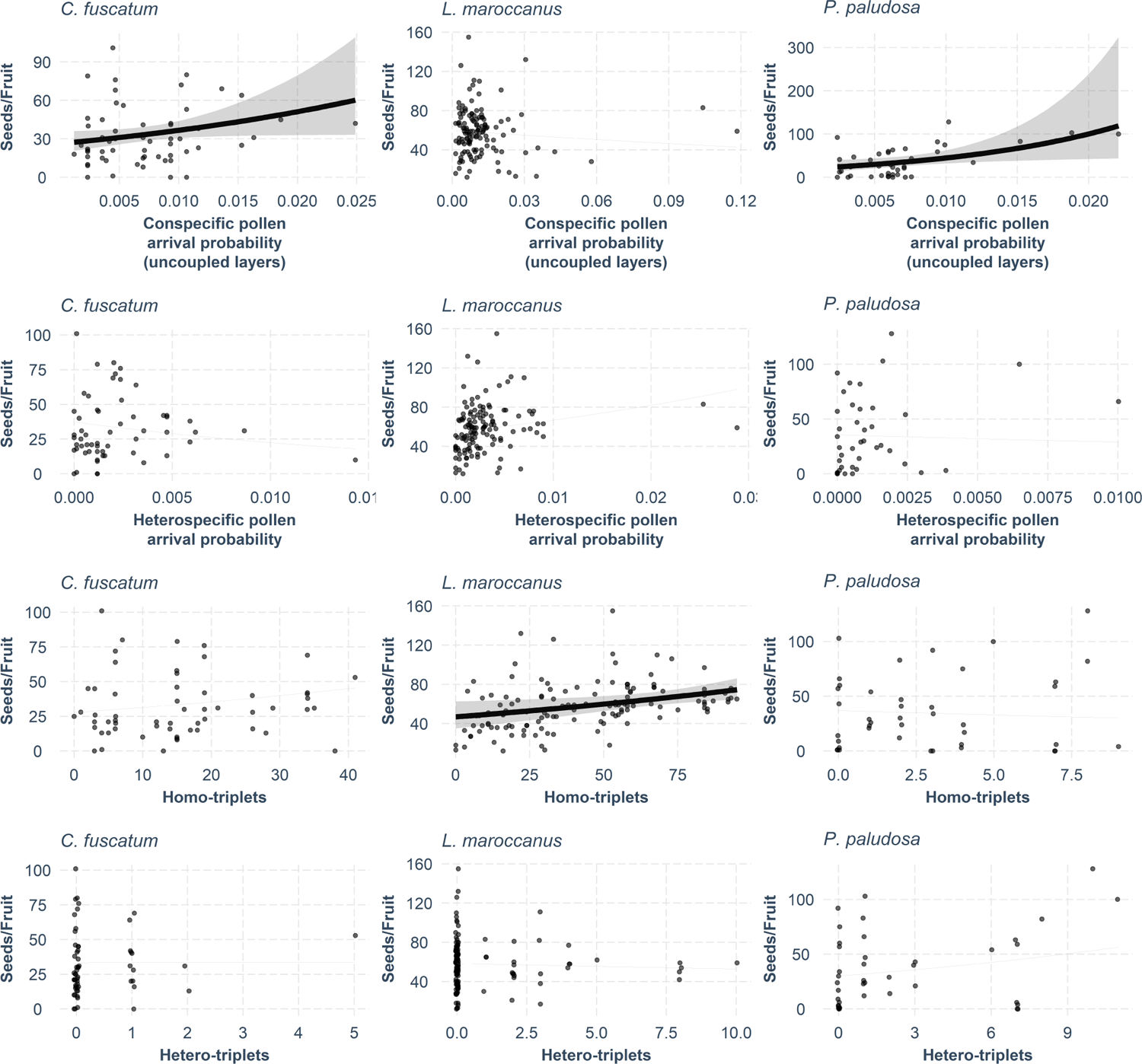
Results for plant’s explanatory variables in the seed production GLMMs (R-package jtools v2.1.4 (Long, 2020)): Conspecific pollen arrival probability (first row), heterospecific pollen arrival probability (second row), homospecific subgraphs (third row), and heterospecific subgraphs (fourth row). In panels with significant explanatory variables (p-value *<* 0.05), we represent (i) the expected values of explanatory variables (black line), (ii) the confidence interval for the expected values (gray band), and (iii) dependence of seed production on the explanatory variable (dark gray dots); otherwise, only the latter is shown.

Regarding the subgraph-level, the total number of homospecific subgraphs only significantly influences the seed production of *L. maroccanus*, and the number of heterospecific subgraphs, in contrast, is not significant for any of the species considered (Supplementary material Appendix 17 Fig. 7 and table A17.1). Plant nodes of *L. maroccanus* follow our expectations: the larger the number of homospecific subgraphs, the larger the increase in seed production. That is, there is an increase in the number of seeds per fruit when plants are immersed in more short-path conspecific connections through shared pollinators.

Finally, since the number of seeds per fruit exhibits a low but significant correlation with visitation rates of *L. maroccanus* and *P. paludosa* (Spearman’s *ρ*s are 0.28 and 0.38, respectively, with p-value *<* 0.05), we tested the results of our models when the covariate “visits” is added to each of them (see Supplementary material Appendix 18). Overall, in the supplementary model for *L. maroccanus* the structural metric that was previously significant continues to be significant, along with the heterospecific subgraphs and the variable “visits”. In contrast, the supplementary models for *C. fuscatum* and *P. paludosa* shows no significant variables other than the zero inflation term, although the effect-sizes of the node- and subgraph-level variables are similar to that of the structural variables presented in our framework and are comparable to the effect-size of “visits”.

## Discussion

Our approach shows that the analysis of plant-pollinator networks at different scales allows a deep understanding of the information flow in such networks, which represents the potential flow of pollen among plant species and, therefore, can be used to better understand the determinants of individual plant fitness. To achieve this, we propose to model the connections between plant individuals mediated by pollinators in each community as a diffusion system, in which there is an ensemble of bipartite layers of conspecific plants that are coupled through insect species that transfer pollen. This multilayer conceptualization allows analyzing structural properties from subgraphs and the entire networks, as well as their relationship with pollen flow. By applying this conceptualization to a highly resolved Mediterranean grassland community we identify a few abundant species that display a mostly conspecific flow of information between their individuals, as opposed to the majority of species, in which information flow involves paths with individuals of other species. We show that these patterns are reflected in individual seed production, with different species being affected by their connections at different scales.

Visualizing species-rich plant communities as individual nodes of each species in their own layer and layers linked via shared pollinators can rapidly help to visualize the within-species and among-species dynamics (Fig. 3). For example, more clustered structures, where interlinks are abundant, may describe potential insect competition among plant species (Pauw, 2013). In addition, as shown in *in-silico* and real multilayer networks, encoding potential pollen flow probabilities in the weight of links highlights that the whole multilayer structure is key for characterizing information spreading at different scales (Masuda *et al*., 2017). For instance, at the network-level, links shape the modular organization of the different plant nodes in modules, whereas at the node-level, connections condition the plant node’s total probabilities of receiving pollen from conspecifics and heterospecifics. While plant-pollinator species modularity has helped depict evolutionary units (Olesen *et al*., 2007), ecological consequences of a modular structure remained elusive (e.g. de Manincor *et al*., 2020). Pioneer studies have shown that individual-level networks show a more modular structure than species-level counterparts (Tur *et al*., 2015). However, a proper assessment of the information flow patterns requires integrating individuals within species as well as using diffusion-like modularity detection algorithms, as proposed here. Using this framework, the next step is to identify which characteristics (e.g. size or location) make individual plants of patches more central (Arroyo-Correa *et al*., 2021).

The proposed framework is flexible enough to incorporate most study designs (e.g. different types of fine-grained sub-specific systems, from individuals of a large-size plant species to groups of small plants belonging to the same patch as in the case study) and study systems (e.g. food-webs, plant-frugivore webs). Moreover, the flexibility of this modeling framework allows refining the quantification of the flow probabilities by considering, for instance, additional phenological constraints or by making the pollinator efficiencies dependent on the spatial distances among plant nodes. Our model, hence, balances out the compromise between traceability and specificity. While other options such as agent-based models can allow a deeper mechanistic characterization of pollen flow, they would need to be tailored specifically to each study system and lose generality. Linking network structure, information flow and seed production can reveal elusive relationships between network structure and functioning (Thompson *et al*., 2012). While the nine analyzed individual plant-pollinator networks show information flow patterns that are not different from a random expectation in the number of modules observed, these modules are not random in their composition. That is, there are ecological processes structuring how plants are grouped together in modules in our system. Overall, modularity partitions based on information flow and diffusion dynamics, as implemented here, allow a phenomenological estimate of indirect effects between individuals of the same and different species, mediated by individuals of other guilds. This can improve the characterization of indirect effects such as apparent competition (Holt, 1977) or pollinator-mediated facilitation or competition effects (Carvalheiro *et al*., 2014) in fine-scale sub-specific networks. For example, the relatively few pollinators linking plant species in our empirical networks (interlinks in our multilayer networks) are key in terms of redistributing pollen flow across the network. Indeed, those insect species that channel interlinks turn out to be network and module hubs of our arrangements (Supplementary material Appendix 14). However, understanding the directionality of indirect effects requires data on plant reproductive success in contrasting situations as similar network patterns can emerge from different processes.

Whether these indirect estimates are reflected in ecological vital rates, such as seed production, is the last question that we tackled in our study system. Previous studies at the species level show weak relationships between network structure and function (i.e. seed production; Ĺazaro *et al*. (2020); Magrach *et al*. (2021)). This is not surprising given the large variability expected in any vital rate across individuals of the same species (Bolnick *et al*., 2003). We found here that conspecific potential pollen transport positively influences seedset in three Mediterranean species. Interestingly, while *C. fuscatum* and P. paludosa show that the probability of pollen arrival when considering the whole network of interactions is correlated with the seed production per fruit, *L. maroccanus* is largely affected at a smaller scale, that is, by the number of homospecific triplets. As seen in the simulations, the number of subgraphs is partially correlated with the number of intralinks considered, while the overall network-level pollen arrival probability varies very little with it. Hence, this might explain why the seedset of the most abundant species, which also receives the most number of visits, *L. maroccanus*, is more sensitive to the subgraphs short-path connections than to the larger structure of the network of interactions. However, differences in their main pollinator guilds Hurtado *et al*. (2023) can also explain the observed results, and more case studies are needed to understand general patterns. In any case, contrary to the general view of a negative effect of heterospecific pollen flow on plant fitness (Ashman & Schoen, 1996; Lopezaraiza–Mikel *et al*., 2007; Morales & Traveset, 2008), our results show that indirect interactions with heterospecifics do not necessarily harm reproductive success. More studies on how pollinator behaviour may preclude heterospecific pollen deposition are needed. All in all, plant seed production is influenced by several factors not accounted for in these analyses a direct test of this framework would benefit from using pollen deposition data. Unfortunately, this one is rare and hard to collect. Yet, despite the uncertainty induced by the noise of the observations, as well as the existence of important variables left out of the models, we observe species-specific patterns in data that suggest that different levels of network structure can explain variations in seed production.

Collecting data to model individual-based networks at the community level is a daunting task, yet as more examples emerge (Arroyo-Correa *et al*., 2023), we will be able with our framework to document generalities and validate untested assumptions. For instance, while we show here that for small annual plants, using fine-scale patch-level observations may be a good proxy of individual-based pollinator visits, the most critical assumption is that individuals linked by the same pollinator species have larger probabilities of mating (Arroyo-Correa *et al*., 2021; Gómez *et al*., 2020). While reasonable at the scales investigated, pollinator behavior (Devaux *et al*., 2014) and morphology are expected to further modulate this relationship. Following individual pollinators to test for floral constancy (e.g. Jakobsson *et al*., 2008), diet breadth overlap between individuals (Brosi, 2016), or analyzing pollen loads on pollinators bodies (Bosch *et al*., 2009) deserves future attention to test this assumption. The advantage of our framework is that all these sources of information can be readily included in it because assessments about different pollinator efficiencies or flow decay with the spatial distance between nodes can be easily incorporated. In addition, since the connections among layers and plant individuals rely on the taxonomic identification of pollinator nodes and resolving those nodes to species level may not always be possible (due to taxonomic or logistical constraints), future research assessing an adequate taxonomic level (or a mixture of levels) for modeling the mating events encoded in our multilayers can be relevant. Here we used morphology-based keys that exploit the morphological differences between the sampled organisms, along with species and family designations. Thus, we expect that resolving those higher taxonomic identifications to species level may reduce the weight and number of interlinks in our networks while altering the number and weights of intralinks. For that reason, a deeper exploration of how topology shapes the probabilities of receiving pollen is, indeed, an interesting venue for a follow-up study. Further, our framework is exemplified by a plant-pollinator system, but it is widely applicable to other types of interaction networks.

Overall, we provide a solid set of flexible tools, and accompanying R code, to thoroughly analyze the structure of individual-based networks. Downscaling interaction network ecology to the individual level may be important to map functions and processes occurring at such scale, and to better capture intraspecific variability. Further research on fine-grain sub-specific, community-wide networks would enable us to link network theory to processes determining the emergent properties of ecological communities at multiple scales.

### Speculations

In our study system, individuals of an abundant plant species like *L. maroccanus* enhance their reproductive success by having a large number of homospecific subgraphs, whereas other less abundant species like *P. paludosa* benefit from their indirect interactions with co-flowering conspecific partners at the network level. How general this pattern is remains to be seen, but the outcome of these metrics may depend on the context, including species abundance, pollinator behavior, or spatial distribution. For example, the spatial clustering and the degree of spatial intermixing among plant species will influence the position of plants within the multilayer structure (i.e., their information arrival probabilities and subgraph composition) over seed production (Thomson *et al*., 2018). This is particularly relevant when upscaling our mechanistic framework with individual-based networks to other interaction types such as individual plant–frugivore interaction networks, where the birds linking different layers cover wider distances and exert more complex behaviours (Crestani *et al*., 2019).

## Supplementary material

### 1 Random-walks on top of bipartite networks

The information/pollen exchanges between conspecific and heterospecific plant nodes in our framework rely on random-walks on top of bipartite networks. Since a random walk on a bipartite graph is periodic, the existence of a unique limiting distribution is not guaranteed when a discrete time approach is used. For instance, if our network consisted only of one plant node linked to one pollinator node, the information (pollen) exchanged between both nodes would move from one node to another at each time step. Thus, the probability of finding the grain of pollen in the plant node will be equal to one at time steps *t* = 0, 2, 4, *· · ·*; and zero for *t* = 1, 3, 5 *· · ·*. Hence, no limiting probability distribution exists when time tends to infinity.

To cope with that issue, we estimated an average limiting distribution (that is, an average probability of finding a grain of pollen in a given plant node when time tends to infinity) under different conditions (Eqs. 3-5 in the main text). In the example above, the average asymptotic probability of finding the grain of pollen in the plant node is 0.5 *×* (1 + 0) = 0.5. Therefore, the only difference with this method when applied to (aperiodic) unipartite networks is this estimation of an average limiting distribution. In our tests, the asymptotic average behavior when time tends to infinity is robust, and thus these proposed methods are well-behaved in bipartite networks such as the ones from our study system.

### 2 Why do we use Infomap here?

Infomap’s approach contrasts with the usual strategy to detect modules in studies of ecological networks, namely: to identify modular partitions that maximize the internal density of links within the modules, by maximizing the objective function *Q*, called modularity, or its generalizations (reviewed in Thébault (2013)). Infomap, which is denoted as the “dynamical perspective of community detection” in (Rosvall *et al*., 2019), unlike other methods, relies on the static structure of the network, but also on random-walk dynamics to determine the optimal modular partition. Although network flows depend on the density of links, the optimization of the map equation and the one of *Q* usually produce different compartments. Since there is no single “true” network partition (Peel *et al*., 2017), here we applied Infomap because that is the method that best matches our interest in flows’ characterization. Furthermore, Infomap has been thoroughly described mathematically and computationally, is widely used in non-ecological disciplines, and its use in ecological applications is increasing (reviewed in Farage *et al*. (2021)).

### 3 Null-models for testing the significance of modularity

After estimating the map equation (in bits, *L_Observed_*) and the number of modules (*m_Observed_*) for the multilayer of a given system, we want to know whether these values are significant compared with a random expectation. To contrast the significance of such measures, we proposed two null models. In our first null model (Null model 1), the total number of observed interactions per plant species and per pollinator is preserved in each layer. That is, given a network, if pollinator *A* is connected to *x* nodes of plant species *B* (through *x* weighted intralinks), then the algorithm generates a new network by replacing the plant nodes in those *x* intralinks that *A* has by another *x* plant nodes of *B* randomly selected with equal probability. These constraints allow us to keep the phenological overlap and the number of interlinks as in the empirical networks.

The second null model (Null model 2) is similar to the first one, but conceptually, the phenological overlap between plant species (mediated by pollinators) is not maintained. In particular, this model preserves the total number of observed interactions per insect visitor and assumes that the weight of interlinks is those of the observed multilayers, but the links of a given pollinator node can be assigned to any plant species. That is, given a network, if pollinator *A* is connected to *x* nodes of plant species *B* and *y* nodes of plant species *C* (through *x* weighted intralinks in layer *B* and *y* intralinks in layer *C*), then the algorithm generates a new network by replacing the plant nodes in those *x* + *y* intralinks that *A* has by another *x* + *y* plant nodes randomly selected (with equal probability) from all the plant nodes that are present in the network, regardless of their species. Considering these two complementary null models allows to test the importance of phenological overlap in explaining the observed modularity patterns.

### 4 Testing significance of modularity

After calculating the total number of homospecific and heterospecific subgraphs for those plant individuals that received insect visits, we contrasted the significance of such values by comparing them with a random expectation. To do so, we proposed a null model in which the total number of observed visits per plant species and per pollinator are preserved in a given temporal unit (e.g., a week, if that is our selected temporal resolution). In that model, we randomly reshuffled the insect visits received by one plant species among those plant individuals that were present in a certain plot, during a given week. Thus, the experimental phenology observed in each plot was preserved. To derive a null distribution of the total number of homospecific and heterospecific subgraphs per focal individual, we generated an ensemble of 500 null networks for each plot and week. For each randomized realization, we calculated the number of homospecific and heterospecific subgraphs per plant individual, insect visitor, and week; and we pooled the results for plant individuals. Then, since null distributions are not usually Gaussian (see Fig. A4.1 for an example), we estimated their confidence intervals for our metrics and tested whether the observed values belong to such intervals (non-significant result) or not (significant result).

**Fig. A4.1:**
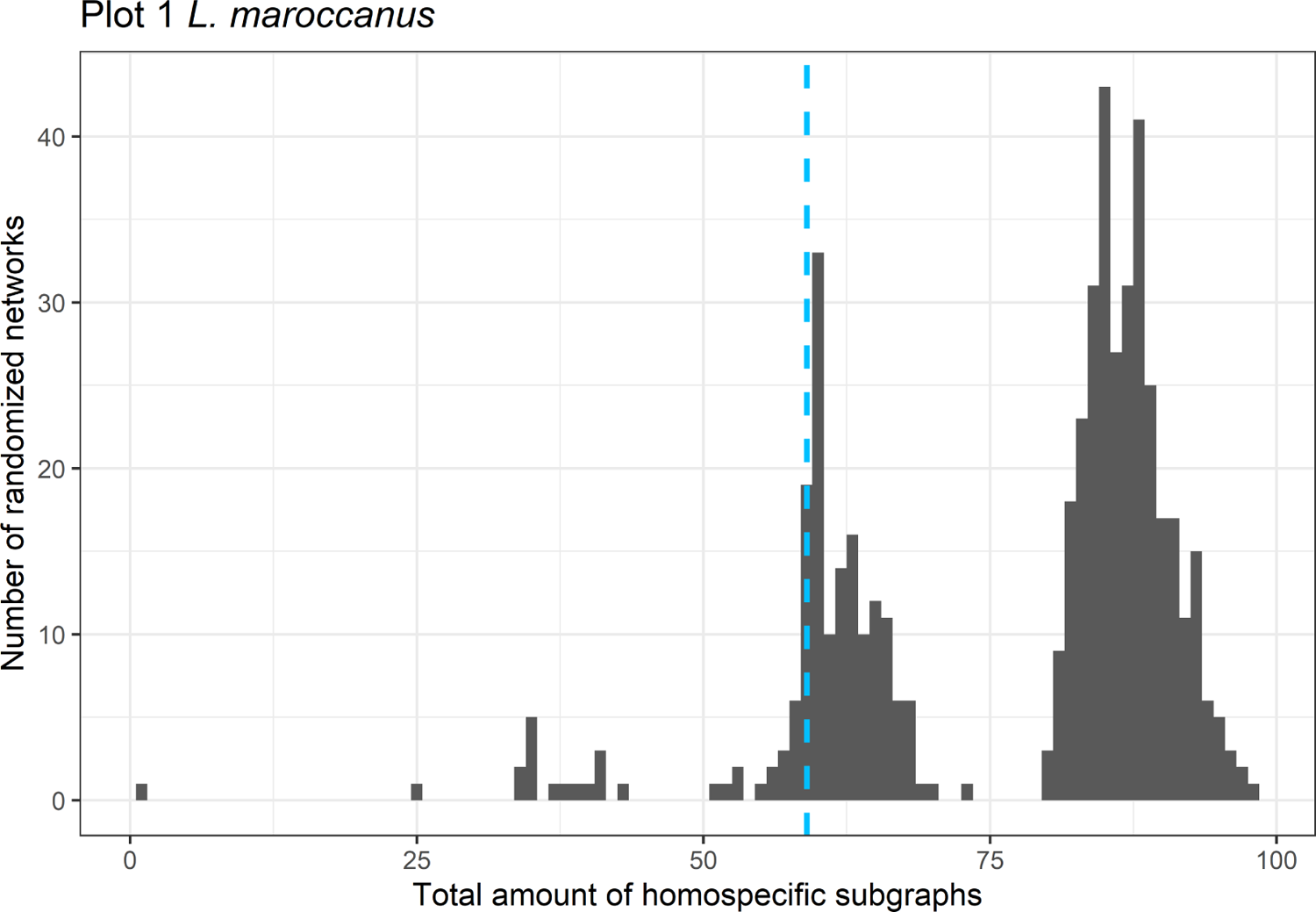
Null distribution of homospecific subgraphs for the focal individual of *L. maroccanus* located in the plot 1 and subplot A3, obtained from 500 simulations. The observed value is marked with a vertical blue dashed line, and it is significantly smaller.

## 5 Description of the random model used in the analysis of network metrics

We built a function in R that creates a data frame with all the information we need to create a multilayer network, following the procedure described in “Multilayer assembly” (Analytical Framework). In table A5.1 we show an example of the resulting data frames.

The function has the following input parameters:

- **number intralinks**: total number of intralinks in the resulting multilayer.
- **number interlinks**: total number of interlinks in the resulting multilayer.
- **number plant individuals**: total number of plant nodes in the resulting multilayer.
- **number pollinator sp**: total number of pollinator nodes in the resulting multilayer.
- **number plant sp**: number of plant species. In our simulations, we set the value of this parameter to 5, the mean number of plant species observed in our plot multilayers.
- **average number visits**: average number of visits received by a plant individual from a pollinator species. In our simulations, we set the value of this parameter to 5.
- **mean phenology plant sp i**: the ISO week number in which we expect the maximum number of visits of pollinators to plant species *i*. We assumed that the number of visits received by the plant species *i* in a given week follows a normal distribution. In each simulation, we randomly generate the parameter mean phenology plant sp i, by randomly sampling a normal distribution with mean equal to 16 and standard deviation equal to 5.
- **sd phenology plant sp i**: the standard deviation of the normal distribution that we used to model the number of visits received by the plant species *i* in a given week. In our simulations, we set the value of this parameter to 2.

Our function randomly distributes the number of interlinks among the pollinator species and, then, it randomly generates interaction matrices between individual plants and pollinator species. Finally, when a random interaction matrix is compatible with the input parameters selected by the user, the random generator stops and that interaction matrix is formatted as the data frame in table A5.1. The process to build a random multilayer is summarized in text Box 5.1.

### Box 5.1

#### Algorithm for generating random multilayers

The process to build a random multilayer can be summarized in the following steps:

1. Define the interlinks of the multilayer. To do so, the algorithm creates an empty “total number of plant species” *×* “total number of pollinator species” interaction matrix with the values provided by the user for those parameters. In the synthetic multilayers we analyzed here, those values are 5 and 25 species, respectively. Then, it randomly generates links between plant and pollinator species that are compatible with the total number of interlinks defined by the user. This matrix is binary, that is, its elements are 0 or 1.
2. Define the total number of plant individuals and intralinks for each plant species. To do this, we randomly sample two integer sets from a normal distribution. Each sample contains as many integers as there are plant species (i.e., 5 integers per sample). In the first sample, the sum of the sampled integers is equal to the total number of plant individuals defined by the user, and each number represents the number of plant individuals of the respective species. In the second set, their sum is equal to the total number of intralinks defined by the user, and each integer represents the intralinks that each plant species (layer) has. Since we are drawing very few integers from a normal distribution (5 integers in total per sample), the total number of plant individuals and intralink per layer tend to be similar.
3. Define the intralinks within each layer. To do so, the algorithm creates an empty “total number of plant individuals” *×* “total number of pollinator species” interaction matrix for each layer. Then, it generates intralinks that meet the following restrictions: i) plant individuals of a given species only interact with the pollinator species identified in step 1; ii) the total amount of intralinks for that layer should be that defined in step 2; and iii) every plant individual should have at least one intralink. These matrices are binary, that is, their elements are 0 or 1.
4. Estimate the total number of pollinator visits received by plant individuals. To do so, we suppose that plant individuals receive on average 5 visits from pollinators (as in our real systems) and estimate the total number of visits received by all plant individuals as 5 *×* “total number of plant individuals”. Next, we distribute that total number of visits among each plant individual, following the same process described in step 2. Thus, we obtain the total number of visits received by each individual plant from the different pollinator species, along the flowering season. Finally, we distribute the visits received by each plant individual among their intralinks as in step 2.
5. Define a synthetic record of pollinator visits to plant individuals. Intra and inter-link weights depend on the phenological overlap of plant species. For that reason, first, we randomly determine the week in which the flowering peak of each plant species takes place by sampling as many numbers as plant species are from a normal distribution (mean = 16, standard deviation = 5). Then, for each visit received by each individual plant we randomly generate the week in which it was registered. To do so, we sample a value from a normal distribution with the mean determined above and a standard deviation equal to 2.
6. Compute the intra- and inter-link weights, as well as the metrics in our toolbox, following the pipeline described in the main text.

Due to the process described above, the number of individuals and intralinks is not evenly distributed among plant species in our synthetic multilayers. However, as mentioned earlier, since we are sampling integers from a normal distribution and we only consider five different plant species, the total number of plant individuals and intralink per plant species (layer) tend to be similar in these synthetic multilayers.

**Table A5.1:** Example of the output of the random model used in the analysis of network metrics.

| Subplot | Plant | Pollinator | Visits | Week |
| --- | --- | --- | --- | --- |
| ind_1 | plant_sp_1 | pol_sp_1 | 3 | 10 |
| ind_1 | plant_sp_1 | pol_sp_2 | 2 | 16 |
| ind_2 | plant_sp_1 | pol_sp_1 | 1 | 12 |
| ... | ... | ... | ... | ... |

## 6 Additional materials for the network sensitivity analyses

**Fig. A6.1:**
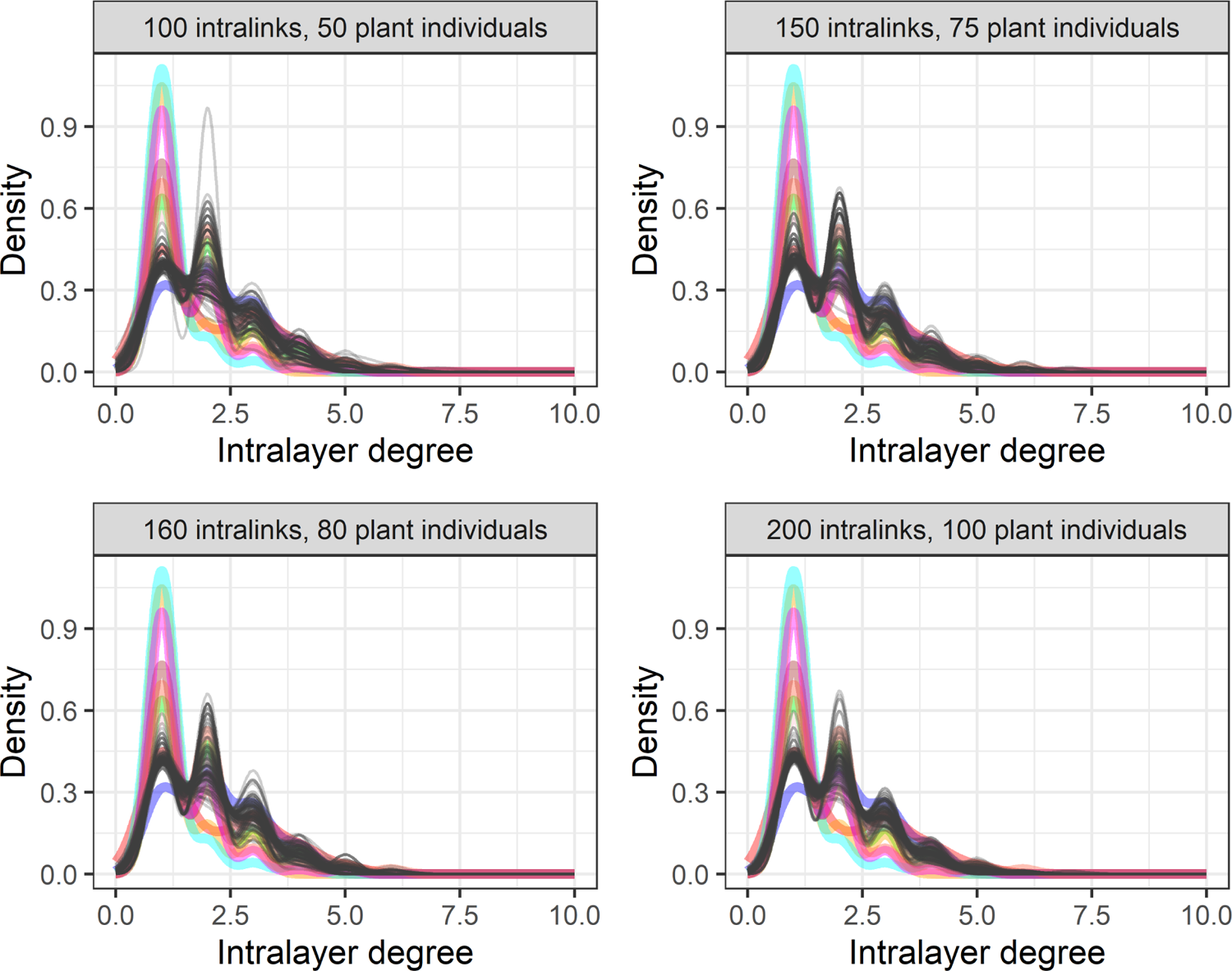
Intralayer degree distribution for synthetic (gray lines) and empirical networks (colored lines). The number of intralinks and plant nodes of the synthetic networks are displayed in each panel, and their number of interlinks and pollinator species are 20 and 25, respectively.

**Fig. A6.2:**
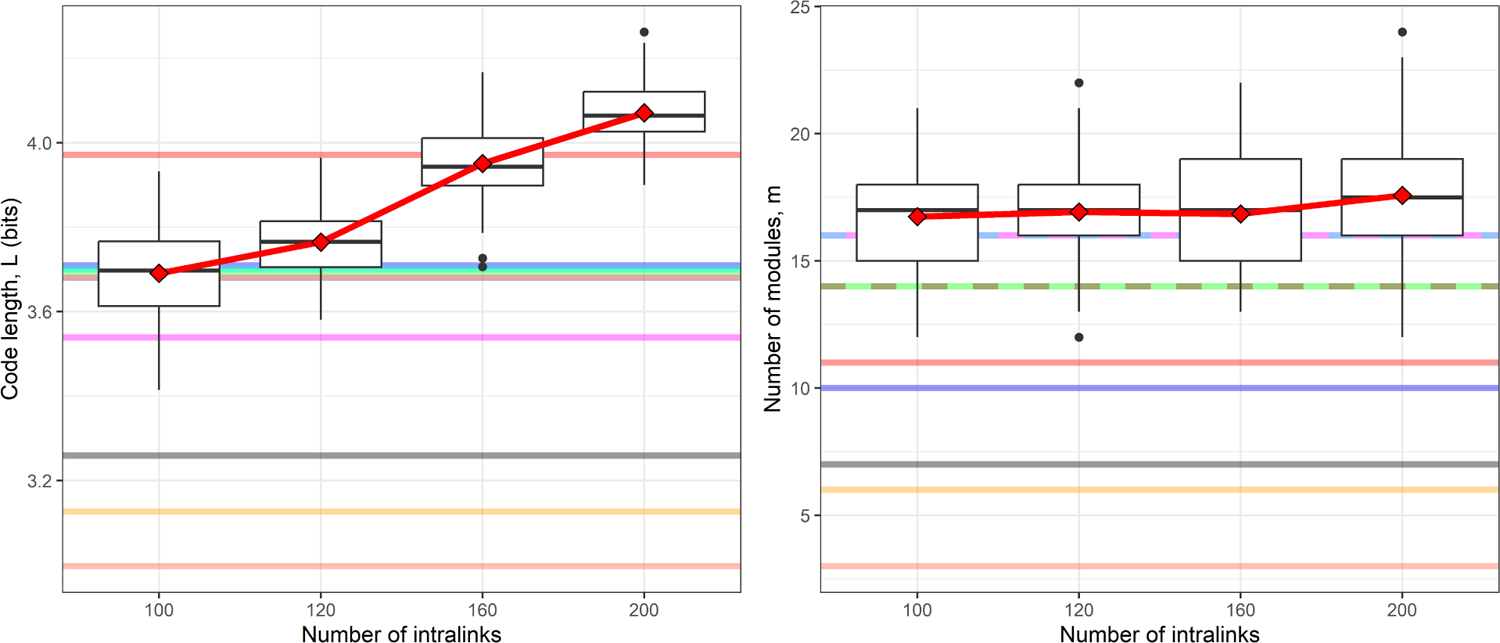
Box-plots for the code length (*L*) and number of modules (*m*) of synthetic, under the constraints in Fig. A6.1. The horizontal colored lines show the values we found in our empirical networks. Some horizontal lines are dashed to allow the representation of empirical values of *m* that overlap. To obtain the results for synthetic networks we assumed that all pollinator species have an efficiency equal to one.

**Fig. A6.3:**
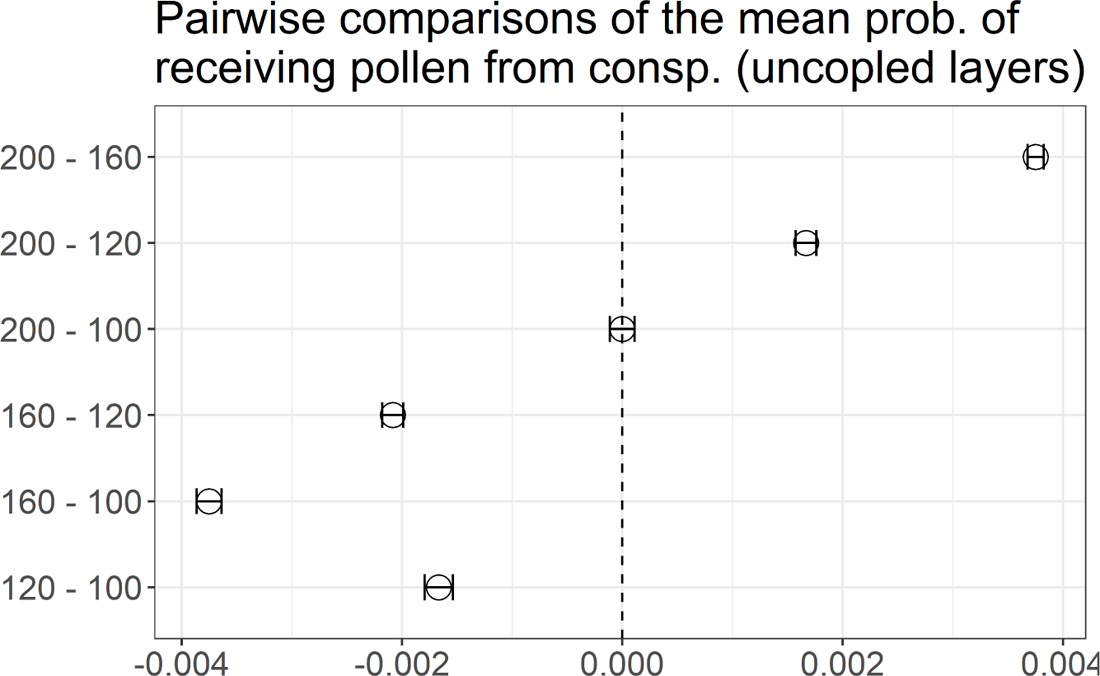
Pairwise comparisons of the mean values of the probability of receiving pollen from conspecifics in Fig. 2 by number of plant nodes. The vertical axis shows the groups (by number of intralinks) for which the mean probability of receiving pollen from conspecifics is compared, and the horizontal axis displays the mean value of the corresponding differences (circle) and 95% confidence intervals (error bars). Vertical dashed line is a guide to the eye that locates mean differences equal to zero. Confidence intervals that do not contain zero indicate a mean difference between groups that is statistically significant (with 95% confidence level). The comparisons of multiple means were performed with the package multcomp v1.4-19 (Hothorn *et al*., 2008) with the procedure proposed by (Herberich *et al*., 2010).

**Fig. A6.4:**
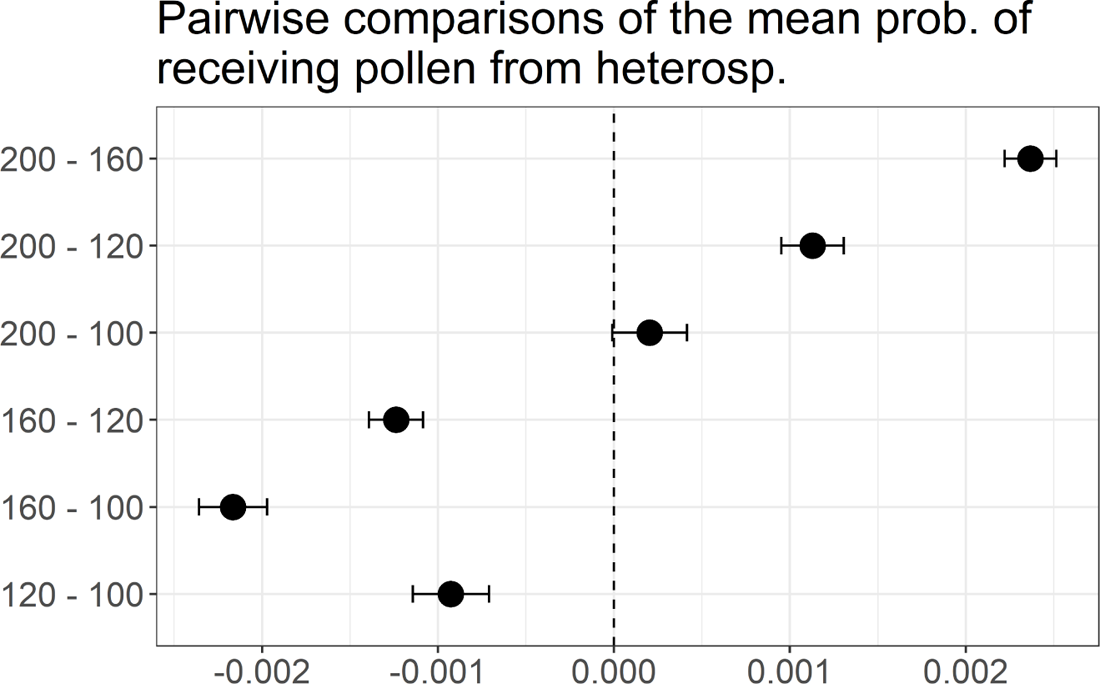
Pairwise comparisons of the mean values of the probability of receiving pollen from heterospecifics in Fig. 2 by number of plant nodes. The vertical axis shows the groups (by number of intralinks) for which the mean probability of receiving pollen from heterospecifics is compared, and the horizontal axis displays the mean value of the corresponding differences (circle) and 95% confidence intervals (error bars). Vertical dashed line is a guide to the eye that locates mean differences equal to zero. Confidence intervals that do not contain zero indicate a mean difference between groups that is statistically significant (with 95% confidence level). The comparisons of multiple means were performed with the package multcomp v1.4-19 (Hothorn *et al*., 2008) with the procedure proposed by (Herberich *et al*., 2010).

**Fig. A6.5:**
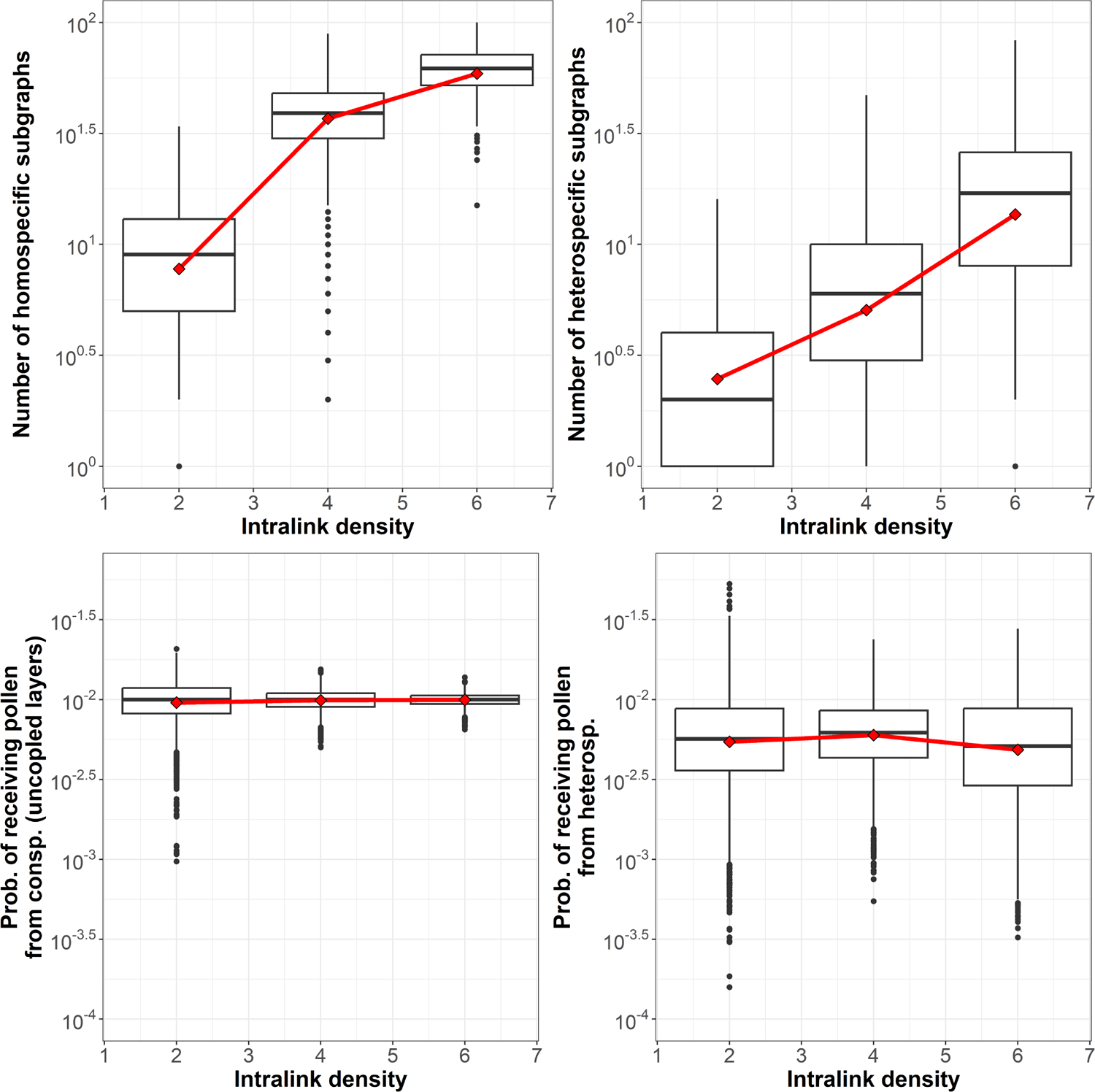
Effect of the intralink density on the subgraph descriptors and probabilities of receiving pollen. Top panels: Dependence of the total number of homospecific subgraphs (left panel), and total number of heterospecific subgraphs (right panel) on the number of intralinks. Bottom panels: Dependence of the probability of receiving pollen from conspecific plant nodes when the layers are uncoupled (left panel), and of the probability of receiving pollen from heterospecific plant nodes (right panel) on the total number of intralinks. Mean values are shown with red diamonds. Red lines are a guide to the eye to highlight the mean value trends.

**Fig. A6.6:**
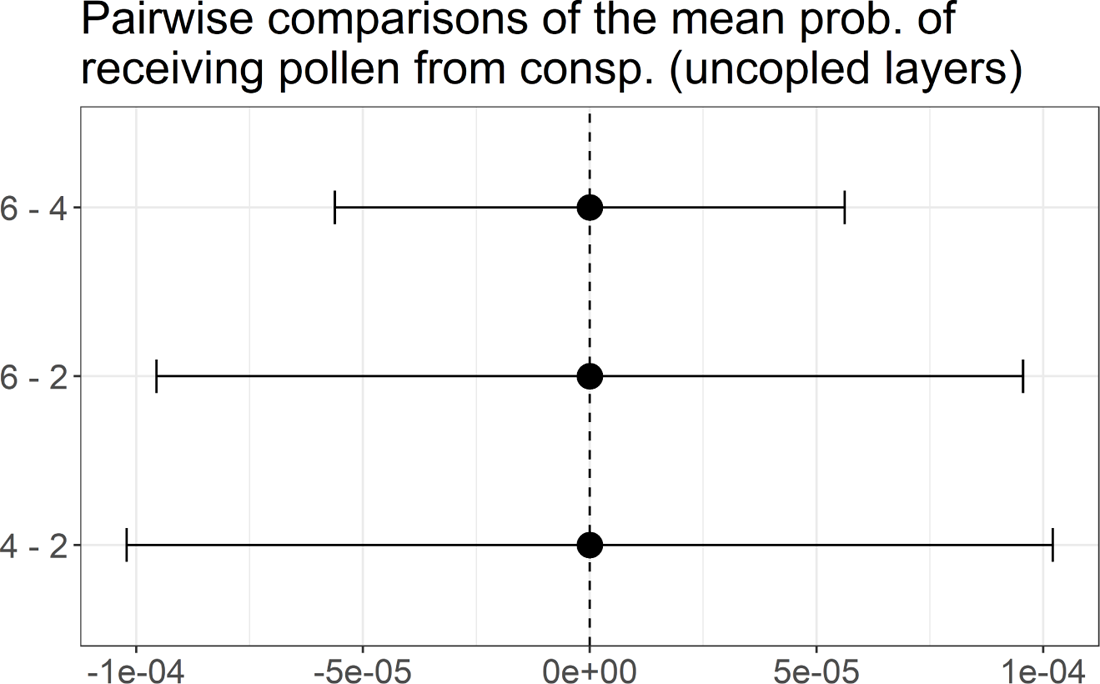
Pairwise comparisons of the mean values of the probability of receiving pollen from conspecifics by link density for the multilayers in Fig. A6.5 by number of plant nodes. The vertical axis shows the groups (by intralink density) for which the mean probability of receiving pollen from conspecifics is compared, and the horizontal axis displays the mean value of the corresponding differences (circle) and 95% confidence intervals (error bars). Vertical dashed line is a guide to the eye that locates mean differences equal to zero. Confidence intervals that do not contain zero indicate a mean difference between groups that is statistically significant (with 95% confidence level). The comparisons of multiple means were performed with the package multcomp v1.4-19 (Hothorn *et al*., 2008) with the procedure proposed by (Herberich *et al*., 2010).

**Fig. A6.7:**
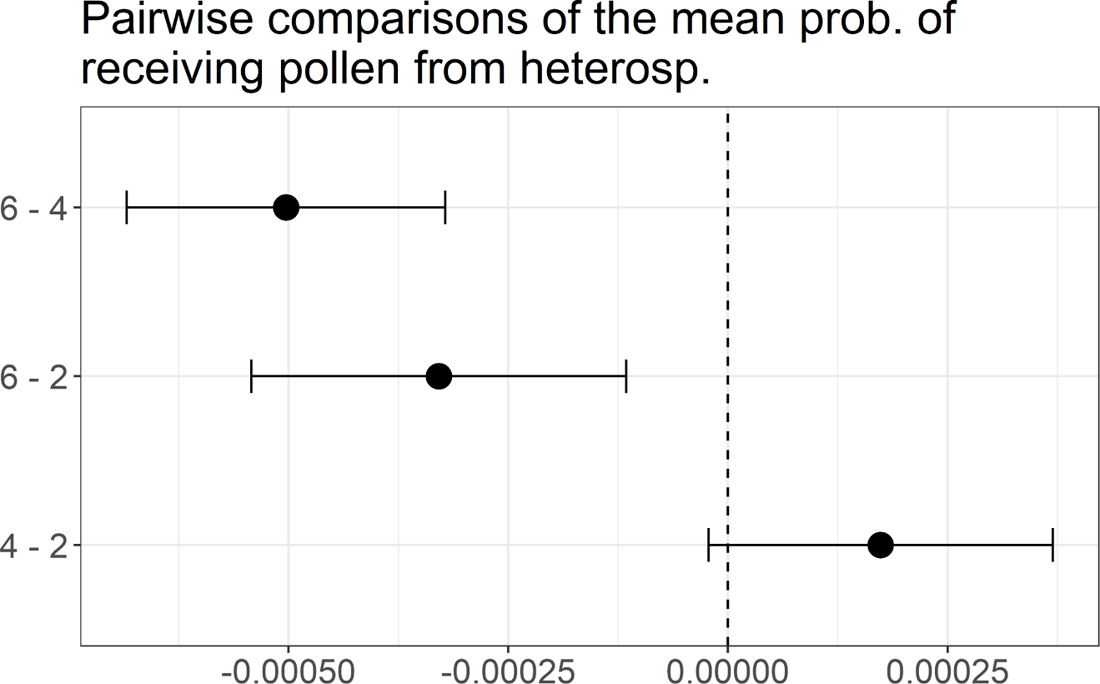
Pairwise comparisons of the mean values of the probability of receiving pollen from heterospecifics by link density for the multilayers in Fig. A6.5 by number of plant nodes. The vertical axis shows the groups (by intralink density) for which the mean probability of receiving pollen from heterospecifics is compared, and the horizontal axis displays the mean value of the corresponding differences (circle) and 95% confidence intervals (error bars). Vertical dashed line is a guide to the eye that locates mean differences equal to zero. Confidence intervals that do not contain zero indicate a mean difference between groups that is statistically significant (with 95% confidence level). The comparisons of multiple means were performed with the package multcomp v1.4-19 (Hothorn *et al*., 2008) with the procedure proposed by (Herberich *et al*., 2010).

**Fig. A6.8:**
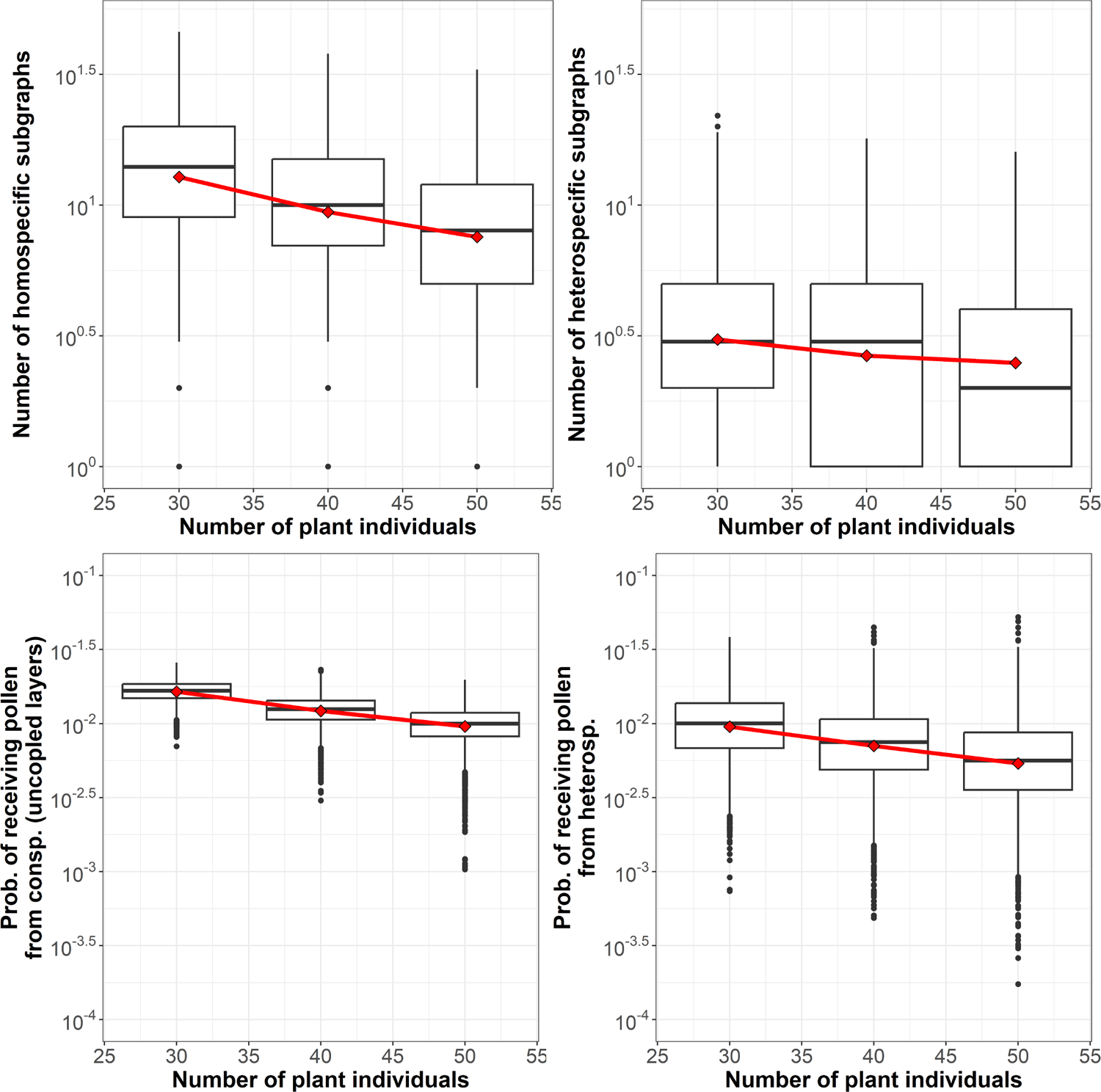
Effect of the number of plant individuals on the subgraph descriptors and probabilities of receiving pollen. Top panels: Dependence of the total number of homospecific subgraphs (left panel), and total number of heterospecific subgraphs (right panel) on the number of intralinks. Bottom panels: Dependence of the probability of receiving pollen from conspecific plant nodes when the layers are uncoupled (left panel), and of the probability of receiving pollen from heterospecific plant nodes (right panel) on the total number of intralinks. Mean values are shown with red diamonds. Red lines are a guide to the eye to highlight the mean value trends.

**Fig. A6.9:**
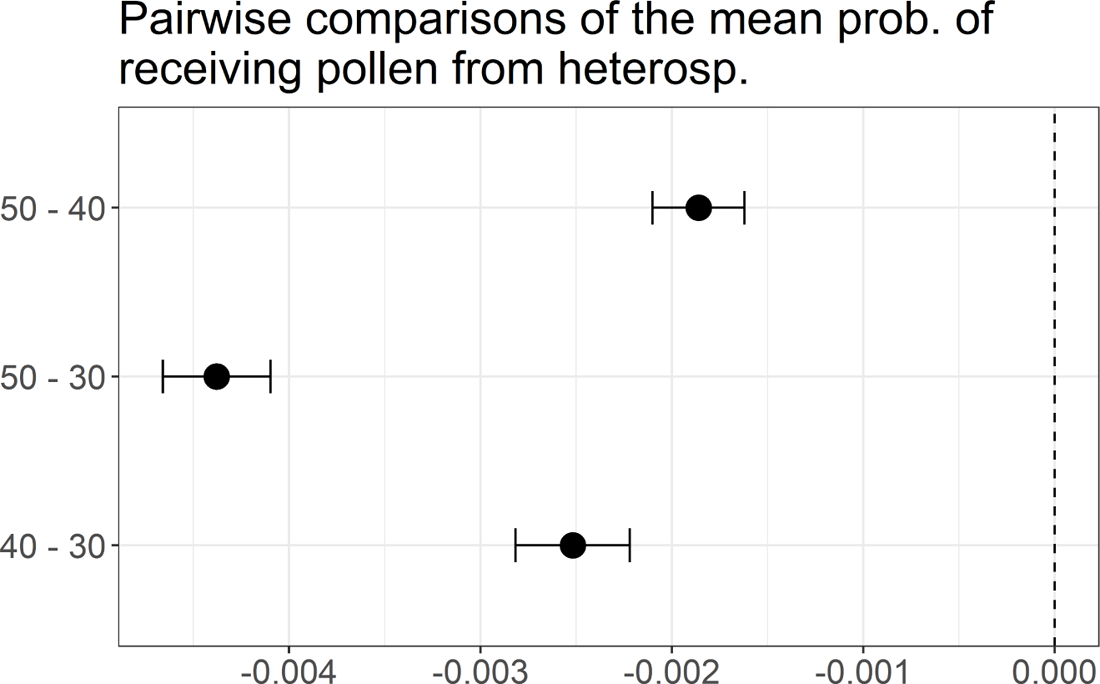
Pairwise comparisons of the mean values of the number of heterospecific motifs by number of plant individuals for the multilayers in Fig. A6.8 by number of plant nodes. The vertical axis shows the groups (by total number of plant nodes) for which the mean number of heterospecific subgraphs is compared, and the horizontal axis displays the mean value of the corresponding differences (circle) and 95% confidence intervals (error bars). Vertical dashed line is a guide to the eye that locates mean differences equal to zero. Confidence intervals that do not contain zero indicate a mean difference between groups that is statistically significant (with 95% confidence level). The comparisons of multiple means were performed with the package multcomp v1.4-19 (Hothorn *et al*., 2008) with the procedure proposed by (Herberich *et al*., 2010).

**Fig. A6.10:**
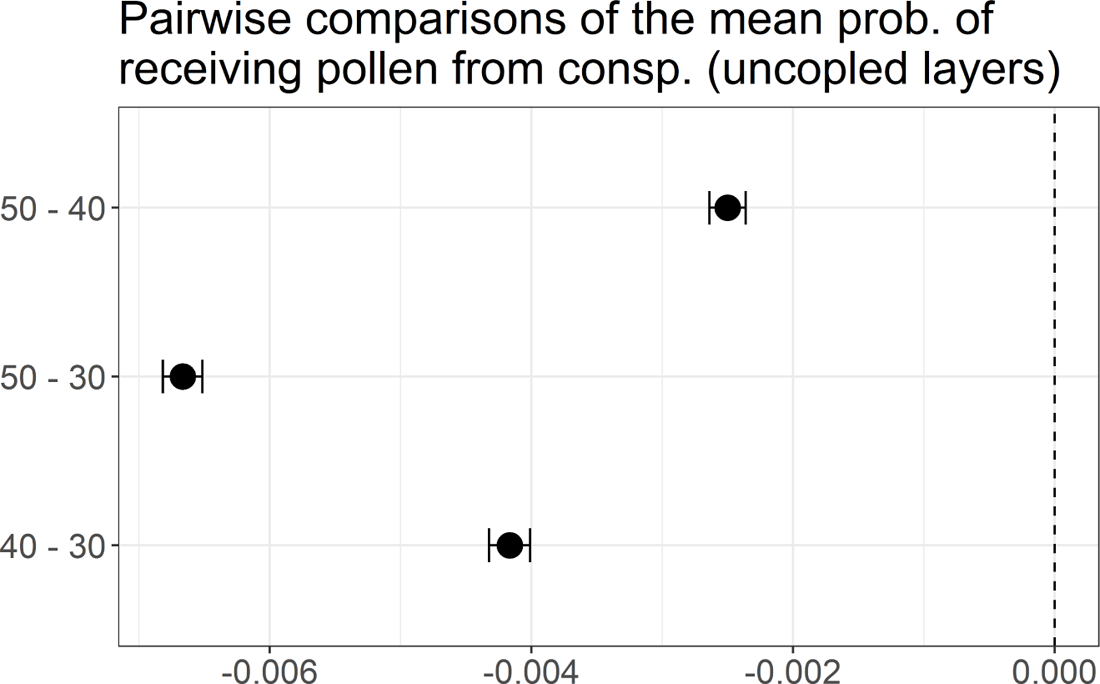
Pairwise comparisons of the mean values of the probability of receiving pollen from conspecifics by number of plant individuals for the multilayers in Fig. A6.8 by number of plant nodes. The vertical axis shows the groups (by total number of plant nodes) for which the mean probability of receiving pollen from conspecifics is compared, and the horizontal axis displays the mean value of the corresponding differences (circle) and 95% confidence intervals (error bars). Vertical dashed line is a guide to the eye that locates mean differences equal to zero. Confidence intervals that do not contain zero indicate a mean difference between groups that is statistically significant (with 95% confidence level). The comparisons of multiple means were performed with the package multcomp v1.4-19 (Hothorn *et al*., 2008) with the procedure proposed by (Herberich *et al*., 2010).

**Fig. A6.11:**
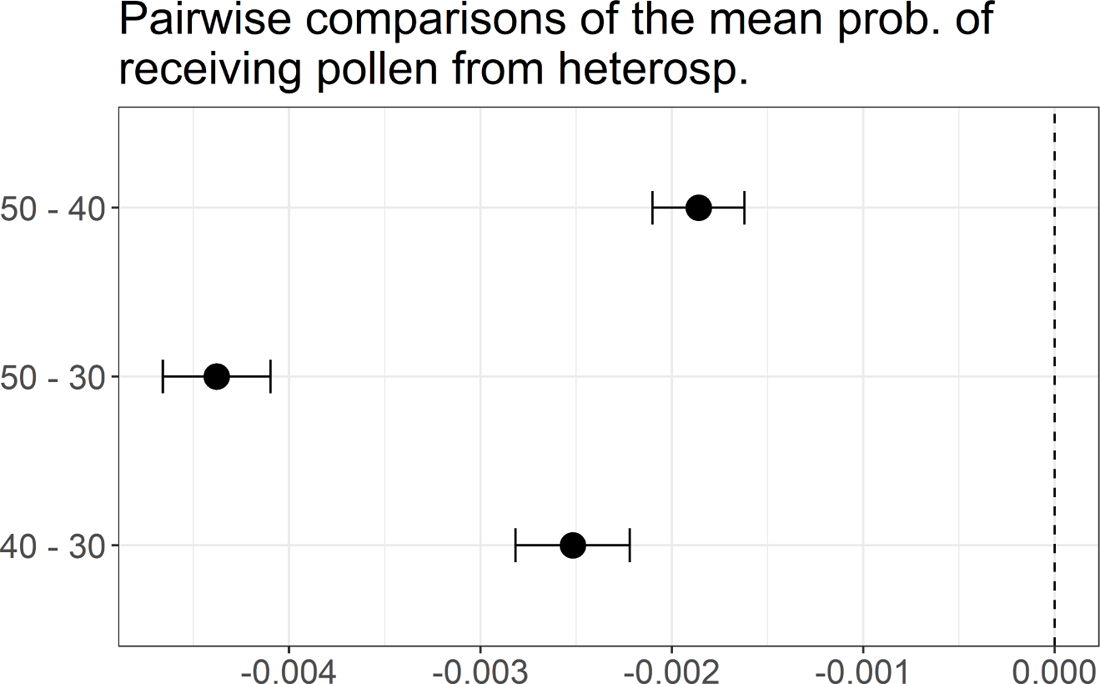
Pairwise comparisons of the mean values of the probability of receiving pollen from heterospecifics by number of plant individuals for the multilayers in Fig. A6.8 by number of plant nodes. The vertical axis shows the groups (by total number of plant nodes) for which the mean probability of receiving pollen from heterospecifics is compared, and the horizontal axis displays the mean value of the corresponding differences (circle) and 95% confidence intervals (error bars). Vertical dashed line is a guide to the eye that locates mean differences equal to zero. Confidence intervals that do not contain zero indicate a mean difference between groups that is statistically significant (with 95% confidence level). The comparisons of multiple means were performed with the package multcomp v1.4-19 (Hothorn *et al*., 2008) with the procedure proposed by (Herberich *et al*., 2010).

### 6.1 The effect of the number of intralinks on subgraph and probability metrics

The effect of the number of intralinks on our subgraph and probability metrics in multilayers where we kept the number of plant individuals, plant species, pollinator species, and inter-links constant (50 plant individuals, 5 plant species, 25 pollinator species, and 20 interlinks). As explained in the main text, since the number of plant individuals is fixed, our analyses are equivalent to studying the effect of intralink density. In addition, note that increasing intralink density while keeping the number of plant individuals and interlinks constant implies increasing the number of intralinks connected to pollinator species with interlinks. Otherwise, the number of interlinks would increase. For that reason, we expect the following direct outcomes: 1) an increase in the number of homospecific and heterospecific subgraphs, but faster for the latter; and 2) a decrease in the asymptotic probability that a heterospecific grain of pollen is transferred to a given plant individual, since pollinator nodes that distribute heterospecific pollen are linked to a larger number of plant individuals.

Our results for our first additional analysis (Fig. A6.5) confirm our expectations for the number of subgraphs (top panels in Fig. A6.5): increasing the intralink density (while keeping constant the number of plant nodes) increases the number of homospecific and heterospecific subgraphs, but the latter grows faster. Regarding the plant node’s total probability of receiving pollen (from all potential sources), our findings show that an increase in link density does not alter significantly the probability of receiving conspecific pollen when layers are uncoupled (left bottom panel in Fig. A6.5, and Fig. A6.6), whereas it significantly reduced the probability of receiving heterospecific pollen (right bottom panel in Fig. A6.5, and Fig. A6.7), confirming our hypothesis for this metric. Nevertheless, our estimation of both asymptotic probabilities indicates that they are mainly driven by the total number of plant individuals (Np) and are inversely proportional to Np (as expected from Eqs. 4 and 5 in the main text). Specifically, the results for the conspecific pollen arrival (when plant species (layers) are isolated) confirm that increasing the intralayer link density only increases the proportion of plant individuals whose probability of receiving a grain of conspecific pollen is equal to the limit value (1/5) *×* (1/2) *×* (1/number of consp. plant ind.). Here, 1/5 represents the fraction of conspecific pollen grains produced within the multilayer (1/ number of plant sp. 10 consp. plant ind./ 50 plant ind.), 1/2 accounts for the probability of finding a grain of pollen in a given pollinator node at time step *t*, and (1 / consp. plant ind. 1/10) is the asymptotic probability that the pollinator node transfers the grain of pollen to a given plant individual at time step *t* + 1 (see Eq. 5 in the main text).

### 6.2 The effect of the number of plant nodes on subgraph and probability metrics

The effect of the number of plant individuals on the previous metrics when keeping the number of intralinks, plant species, pollinator species, and inter-links constant (100 intralinks, 5 plant species, 25 pollinator species, and 20 interlinks). Since intralinks are randomly generated in our model and their number is fixed, we expect that increasing the number of individual plants will increase the number of pollinator nodes linked to just one plant individual. For that reason, we expect a decrease in the number of homospecific and heterospecific subgraphs. Regarding the asymptotic probabilities, we anticipate that increasing the number of plant individuals will reduce the asymptotic probability of receiving pollen from both conspecifics and heterospecifics because both probabilities are inversely proportional to the total number of plant individuals by definition (Eqs. 4 and 5 in the main text).

The results of the above study (Fig. A6.8) show the expected decreasing trends in all four metrics when the total number of plant individuals increases. Those trends were significant (see Figs. A6.9-A6.11).

## 7 List plant species observed in Caracoles (2020)

**Table A7.1:** List of plant species observed in Caracoles Ranch with floral visitors from a total of 23 species surveyed in the community. Taxonomic family of each species is provided. Sample size represents the total number of individuals sampled for reproductive success for each focal species, and it is correlated with their natural abundance observed at the study site.

| Species | Family | Abundance | Sample size |
| --- | --- | --- | --- |
| *Beta macrocarpa* | Amaranthaceae | 1,992 | 250 |
| *Centaurium tenuiflorum* | Gentianaceae | 1,946 | 190 |
| *Chamaemelum fuscatum* | Asteraceae | 1,347 | 151 |
| *Chamaemelum mixtum* | Asteraceae | 143 | 14 |
| *Leontodon maroccanus* | Asteraceae | 9,090 | 269 |
| *Melilotus sulcatus* | Fabaceae | 1,100 | 53 |
| *Pulicaria paludosa* | Asteraceae | 1,418 | 137 |
| *Scorzonera laciniata* | Asteraceae | 791 | 117 |
| *Sonchus asper* | Asteraceae | 1,012 | 106 |
| *Spergularia rubra* | Caryophyllaceae | 2,318 | 58 |

## 8 Plant phenologies

Plant phenologies were extracted from our records of insect visits. Detailed weekly surveys of floral visitors during the flowering season showed that insects only visited the flowers of 10 of the species, and most of those visits were concentrated on only three of them (*Chamaemelum fuscatum*, *Leontodon maroccanus* and *Pulicaria paludosa*). As can be seen in Fig. A8.1, the phenologies of the plant species that received insect visits are varied, being able to differentiate early, medium and late phenologies. The earlier species is *C. fuscatum*; the medium species are *L. maroccanus*, *Spergularia rubra*, *Scorzonera laciniata*, *Sonchus asper*, *Chamaemelum mixtum*, *Melilotus sulcatus* and *Beta macrocarpa*; and the late species are *P. paludosa*, *Centaurium tenuiflorum*, and in 2020 a second flowering peak of *L. maroccanus*.

To estimate the duration of plant phenologies (in weeks), we assumed that, if a plant species received insect visits in weeks *i* and *j* (for *i ̸*= *j*), then such plant species were also visited during the period between *i* and *j*. For instance, *B. macrocarpa*’s phenology lasts 6 weeks (from the 17th week to the 22nd one).

Finally, in Fig. A8.2 we disaggregate the phenological overlap among plant species by using the taxonomic groups of flower visitors.

**Fig. A8.1:**
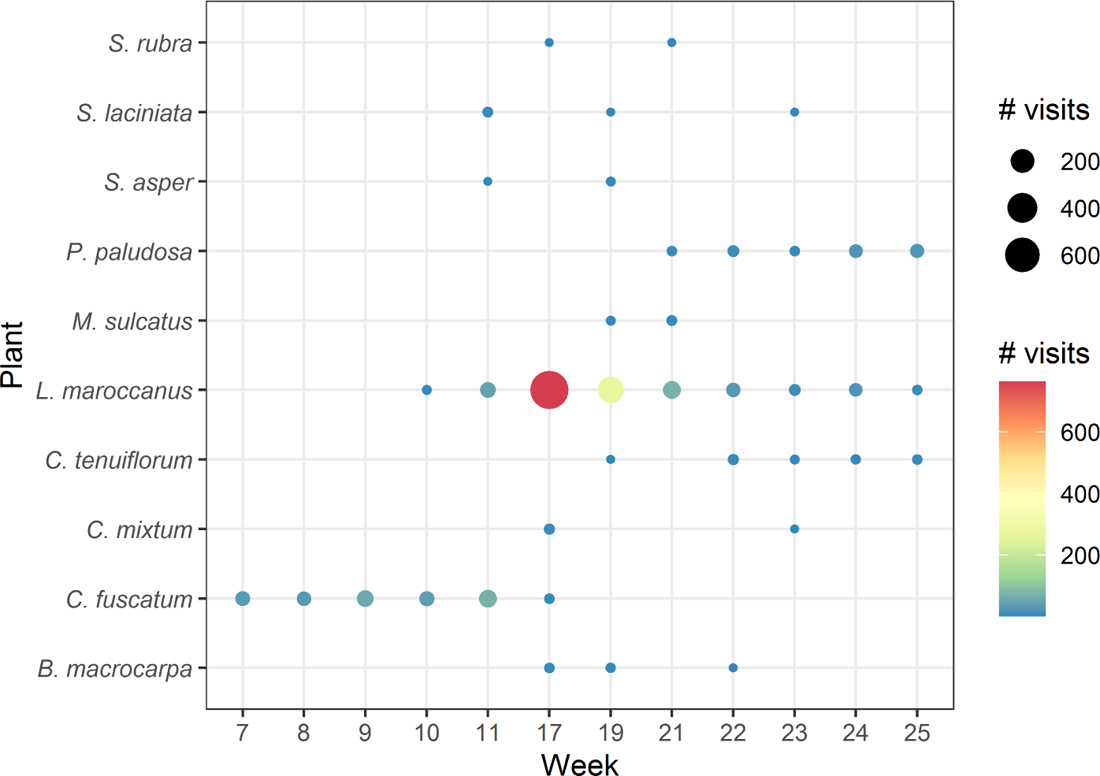
Total number of insect visits received by plant species per week in 2020.

**Fig. A8.2:**
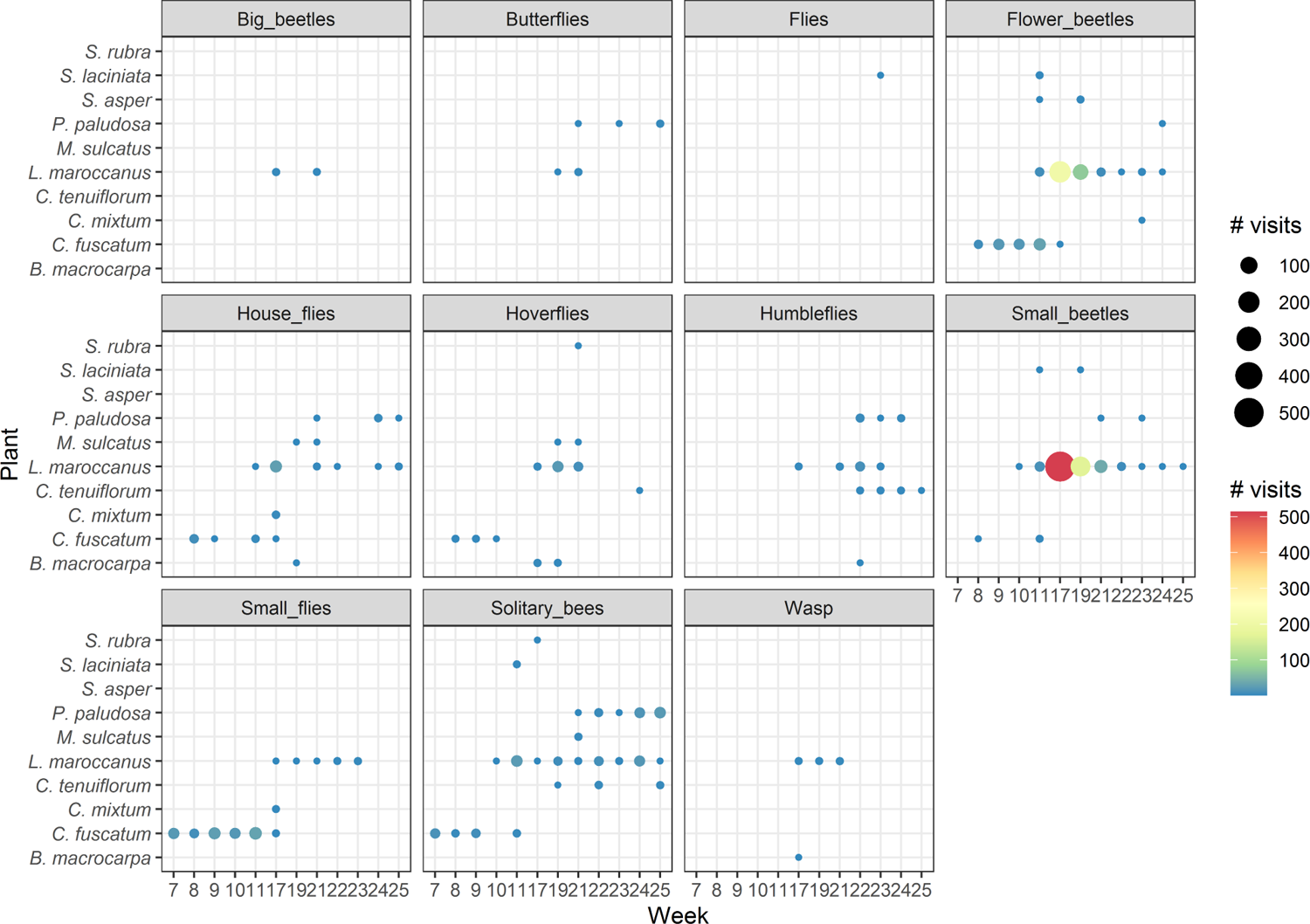
Total number of insect visits received by plant species and taxonomic group of visitors per week in 2020.

## 9 Sampling coverage of Caracoles (2020)

We have estimated the sampling coverage of (individual) plant–pollinator interactions in each plot network, pooling the data for all plant species, and using the coverage framework introduced in (Chao *et al*., 2014). The sampling coverage (or coverage) of the observed interactions is simply their total relative abundances, or equivalently, the proportion of the total number of interactions in an assemblage that belong to interactions represented in the sample. To calculate the coverage in each plot, we used the R-package iNEXT v3.0.0 (Hsieh *et al*., 2016). As can be seen in Fig. A9.1, the sampling coverage in all the plots is approximately equal to 90%. This means that less than 10% of the total interactions in the assemblage belong to undetected interactions. For this reason, extremely rare, undetected interactions do not make a significant contribution to that proportion, even if there are many such interactions. In addition, note that the above estimations do not account for the existence of forbidden interactions, that is, non-occurrences of pairwise interactions that can be accounted for by biological constraints, such as spatio-temporal uncoupling, size or reward mismatching, foraging constraints and physiological–biochemical constraints (Jordano, 2016). Hence, the real sampling coverage of our pollination networks could be higher than 90%.

Finally, it is important to note that all empirical networks may be biased by incomplete sampling, for instance, failing to detect the interactions of rare species (Blüthgen, 2010; Blüthgen *et al*., 2008). This uncertainty affects the community detection presented in this work, as well as the network metrics we discussed so far. For instance, the map equation assumes complete data and, when networks are sparse and may contain missing links, the optimal solution is distorted (Smiljaníc *et al*., 2020). To overcome these obstacles, further investigation should consider the use of improved flow-based community detection algorithms in development (Smiljaníc *et al*., 2020; Young *et al*., 2020, 2021).

**Fig. A9.1:**
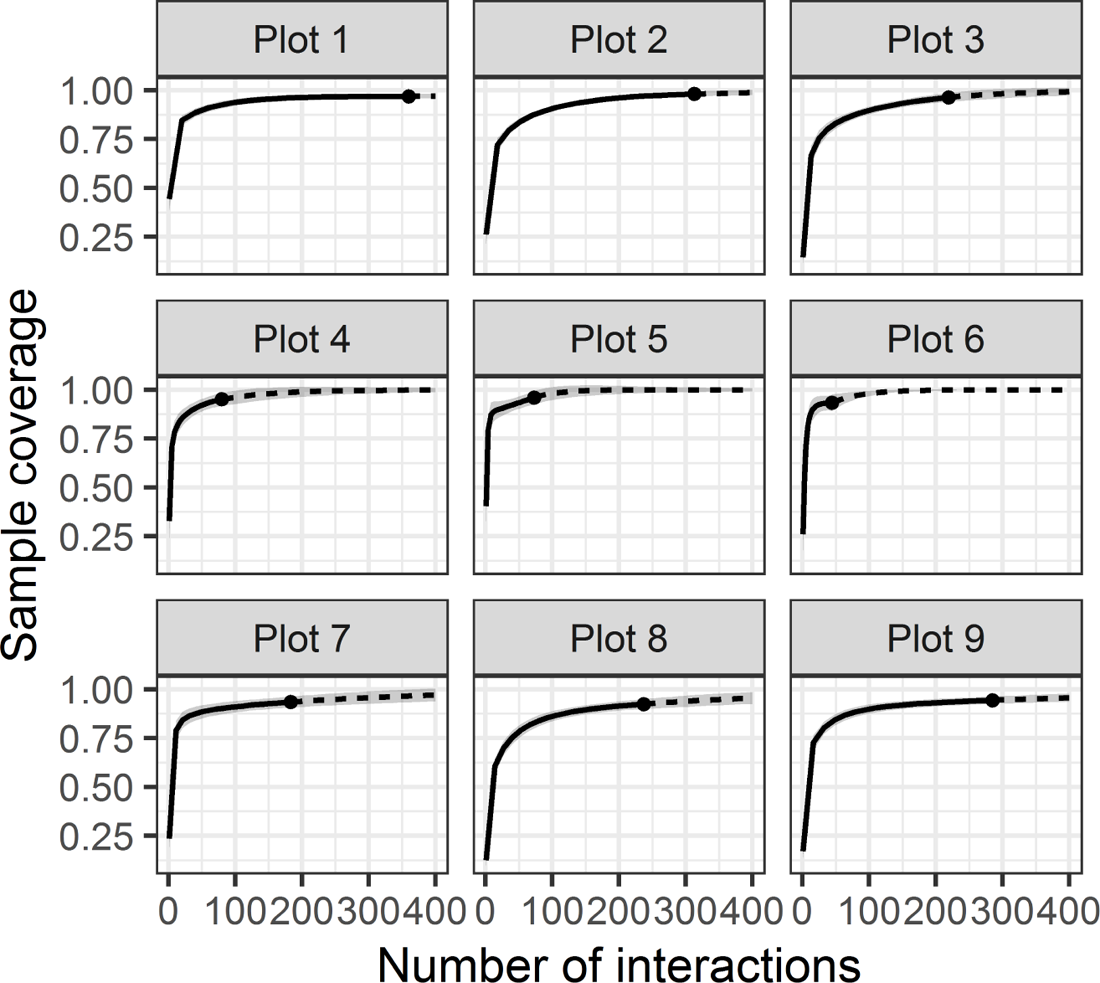
Sample coverage for rarefied samples (solid line) and extrapolated samples (dashed line) as a function of sample size for interaction samples from the plot networks studied. The 95% confidence intervals were obtained by a bootstrap method based on 200 replications. Reference samples are denoted by solid dots.

## 10 List of taxonomic groups, families and species of Caracoles (2020)

**Table A10.1:** Number of visits observed in Caracoles Ranch by taxonomic group, family and identifier.

| Taxonomic groups, Family, ID | Visits |
| --- | --- |
| **Big beetles** | **5** |
| **Cerambycidae** | **2** |
| Cerambycidae | 2 |
| **Meloidae** | **3** |
| *Lagorina sericea* | 3 |
| **Butterflies** | **13** |
| **Geometridae** | **2** |
| Geometridae | 2 |
| **Lasiocampidae** | **5** |
| *Lasiocampa trifolii* | 5 |
| **Nymphalidae** | **2** |
| *Vanessa cardui* | 2 |
| **Pieridae** | **4** |
| *Euchloe crameri* | 1 |
| *Pieris brassicae* | 3 |
| **Flies** | **1** |
| **Diptera** | **1** |
| Other diptera | 1 |
| **Flower beetles** | **395** |
| **Chrysomelidae** | **6** |
| *Cryptocephalus sp.* | 6 |
| **Elateridae** | **13** |
| Elateridae | 13 |
| **Melyridae** | **345** |
| *Malachius bipustulatus* | 9 |
| Melyridae | 1 |
| *Psilothrix viridicoerulea* | 335 |
| **Mordellidae** | **16** |
| Mordellidae | 16 |
| **Oedemeridae** | **15** |
| Oedemeridae | 15 |
| **House flies** | **82** |
| **Calliphoridae** | **6** |
| Calliphoridae | 6 |
| **Muscidae** | **44** |
| *Musca sp.* | 44 |
| **Sarcophagidae** | **23** |
| *Sarcophaga sp.* | 23 |
| **Tachinidae** | **9** |
| *Cylindromyia sp.* | 9 |
| **Hoverflies** | **62** |
| **Syrphidae** | **62** |
| *Episyrphus balteatus* | 14 |
| *Eristalis sp.* | 3 |
| *Eupeodes corollae* | 1 |
| *Lomatia sp.* | 9 |
| *Sphaerophoria scripta* | 32 |
| Syrphidae | 3 |
| **Humbleflies** | **57** |
| **Bombyliidae** | **57** |
| *Anastoechus sp.* | 44 |
| *Bombilus major* | 13 |
| **Small beetles** | **756** |
| **Anthicidae** | **10** |
| Anthicidae | 10 |
| **Chrysomelidae** | **2** |
| *Cassida sp.* | 1 |
| *Phaedon sp.* | 1 |
| **Curculionidae** | **1** |
| Curculionidae | 1 |
| **Nitidulidae** | **743** |
| *Brassicogethes sp.* | 743 |
| **Small flies** | **135** |
| **Bibionidae** | **4** |
| *Dilophus sp.* | 4 |
| **Empididae** | **2** |
| *Empis sp.* | 2 |
| **Stratiomyidae** | **6** |
| *Nemotelus sp.* | 6 |
| **Ulidiidae** | **123** |
| Ulidiidae | 123 |
| **Solitary bees** | **280** |
| **Andrenidae** | **136** |
| *Andrena argentata* | 4 |
| *Andrena humilis* | 76 |
| *Andrena sp.* | 56 |
| **Apidae** | **4** |
| *Eucera sp.* | 4 |
| **Halictidae** | **126** |
| *Lasioglossum immunitum* | 4 |
| *Lasioglossum malachurum* | 111 |
| *Lasioglossum sp.* | 11 |
| **Megachilidae** | **14** |
| *Osmia ligurica* | 14 |
| **Wasp** | **8** |
| **Braconidae** | **8** |
| Braconidae | 8 |
| **Total** | **1,794** |

## 11 Information transfer efficiencies for the taxonomic groups of Caracoles Ranch (2020)

**Table A11.1:** Efficiencies *ε_ij_* used to compute the weight of intralinks in the case study of Caracoles Ranch.

| Taxonomic group of the pollinator<br>node involve in the intralink | $\varepsilon_{ij}$ |
| --- | --- |
| Small beetles | 0.50 |
| Big beetles | 0.50 |
| Butterflies | 0.75 |
| Flies | 0.75 |
| Flower beetles | 0.50 |
| House flies | 0.75 |
| Hoverflies | 0.75 |
| Humbleflies | 0.75 |
| Small flies | 0.75 |
| Solitary bees | 1.00 |
| Wasps | 0.75 |

## 12 Characterization of community structure

As can be seen in Fig. A12.1, the main visitors by functional groups were small beetles (756), followed by pollenivorous flower beetles (395), solitary bees (280) and small flies (135). The remaining visits are distributed between house flies (82), hoverflies (62), humbleflies (57), butterflies (13), wasps (8), big beetles (5) and flies(1). Regarding plants, during 2020, only ten focal species received insect visits in Caracoles, namely: *L. maroccanus* (1,337), *C. fuscatum* (268), *P. paludosa* (111), *C. tenuiflorum* (23), *M. sulcatus* (15), *B. macrocarpa* (14), *C. mixtum* (14), *S. laciniata* (7), *S. asper* (3), and *S. rubra* (2).

Regarding insect nodes per module, the average amount of them is 2.12 with CI (1, 6), and the amount of different species and morphospecies per module is 1.57 with CI (1, 7.92). Overall, there are 64 modules with a single insect species (or morphospecies), 24 with two different species, 4 modules with three, 2 modules with four, 1 with 5, and 2 modules with 8. Additional details on modules features can be found in table A12.1.

**Fig. A12.1:**
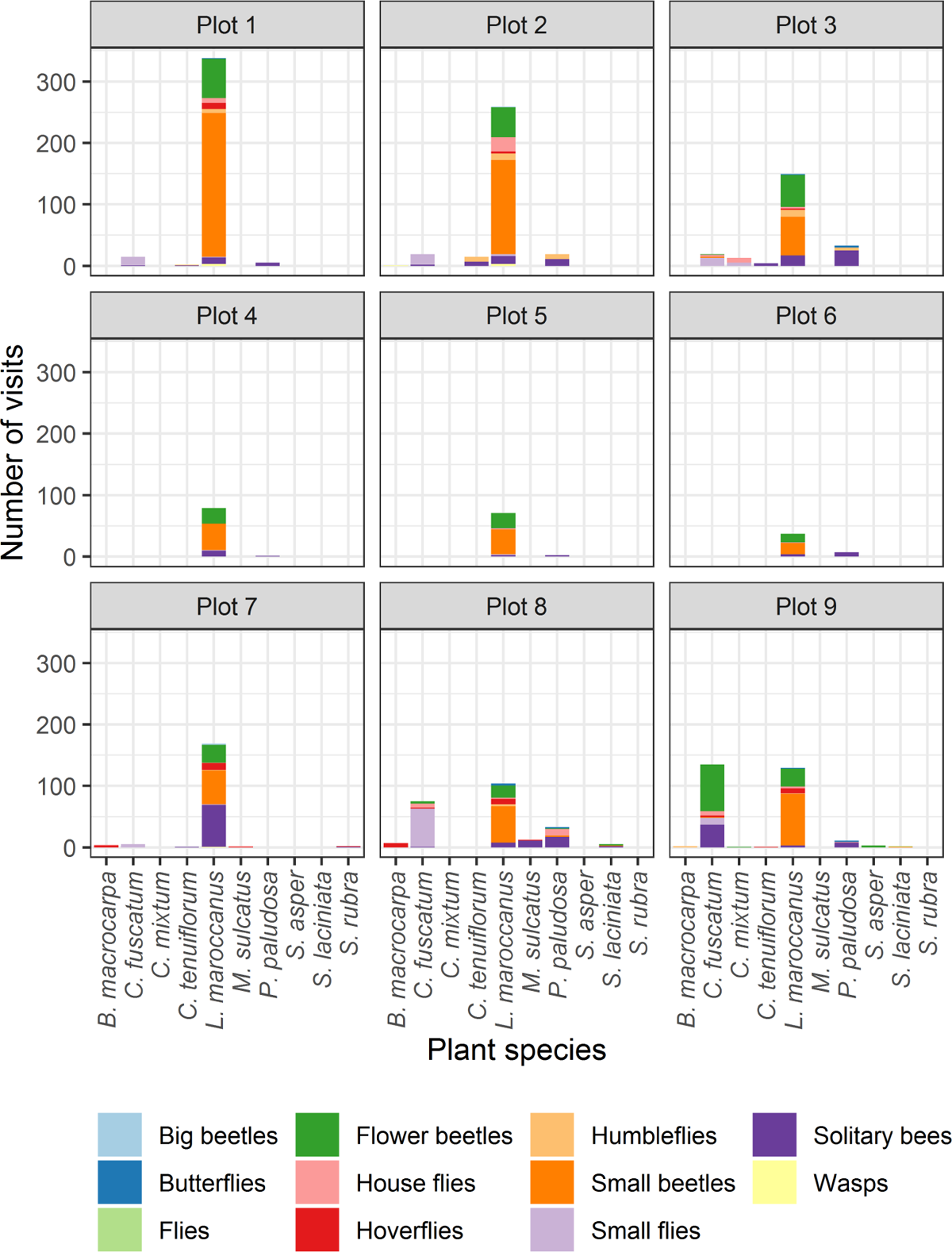
Number of insect visits per functional group, plant species and plot.

**Fig. A12.2:**
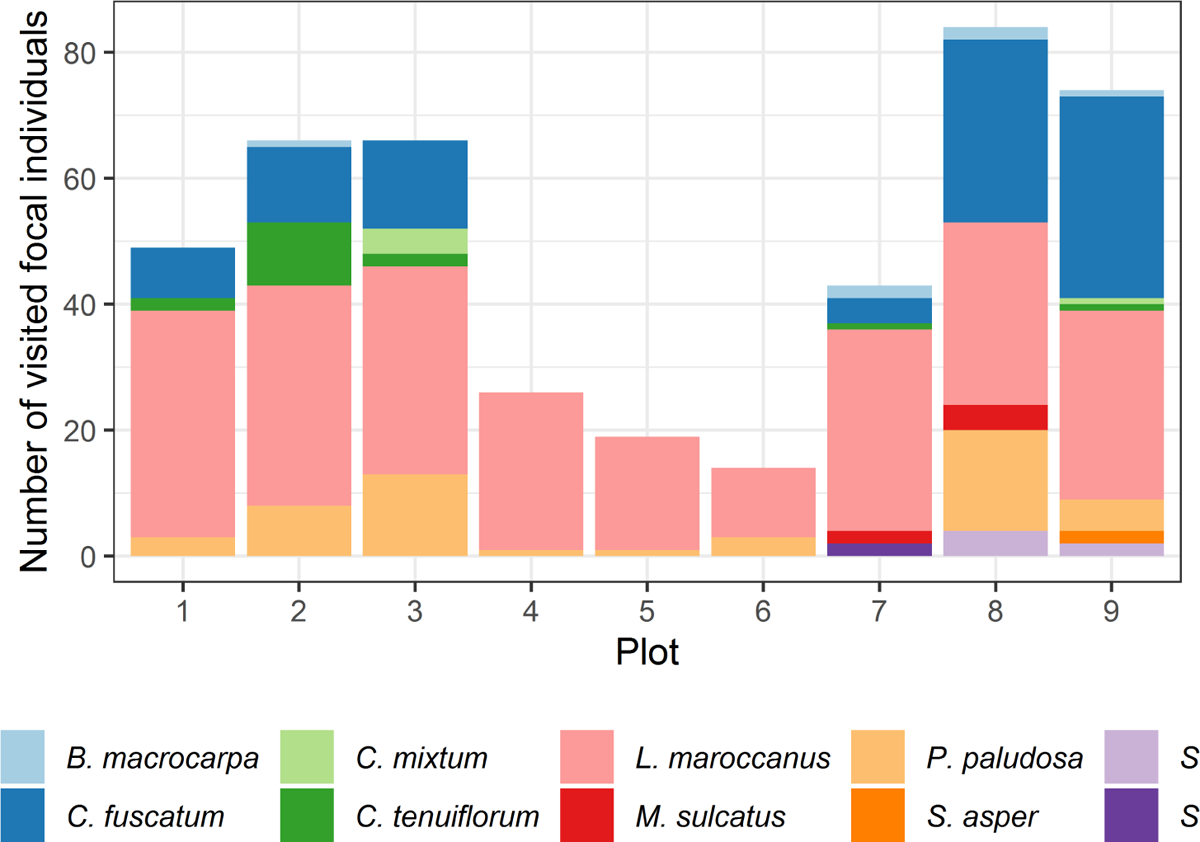
Number of individuals with visits per plant species.

**Fig. A12.3:**
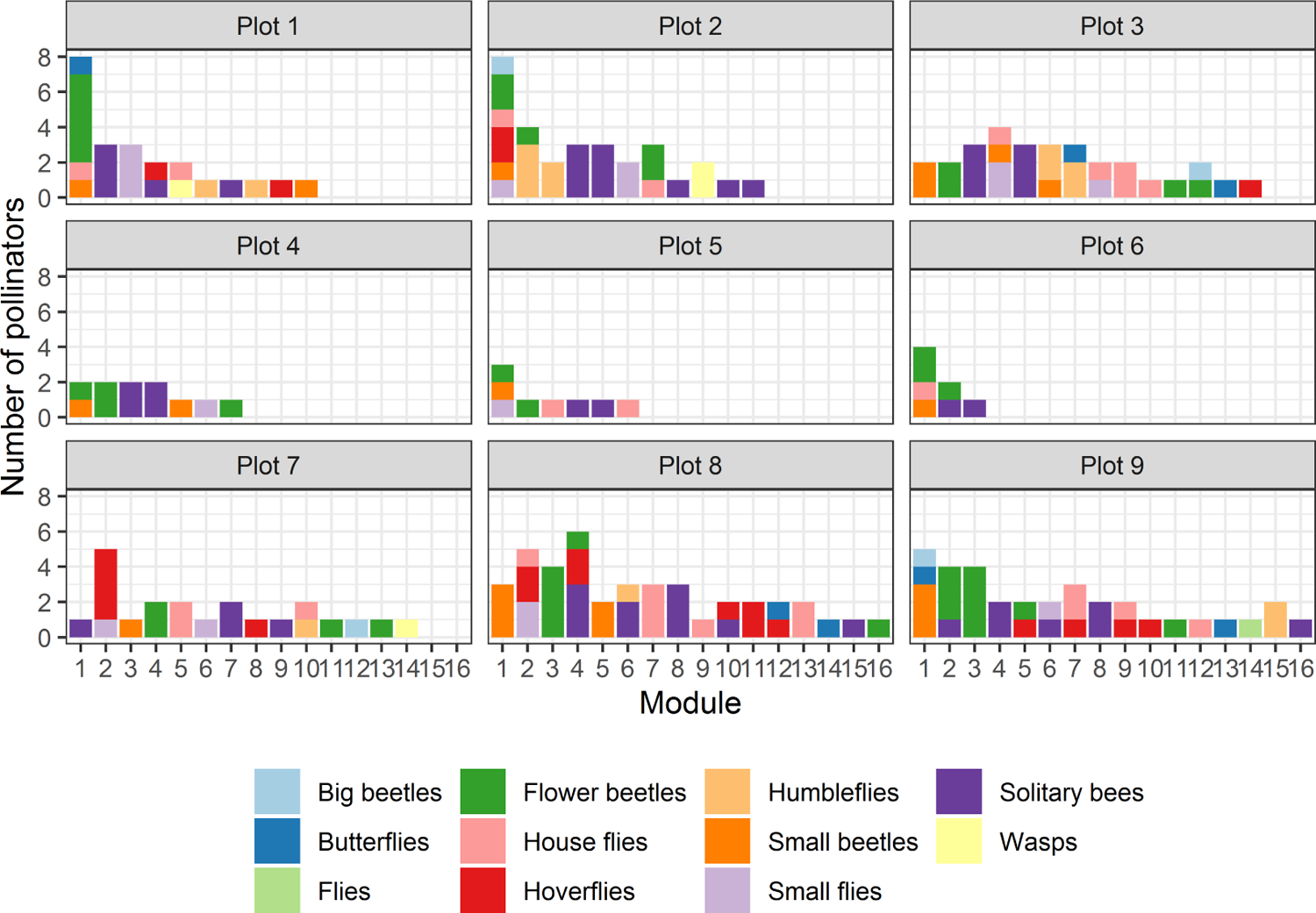
Number of insect visitors per module, taxonomic group and plot.

**Fig. A12.4:**
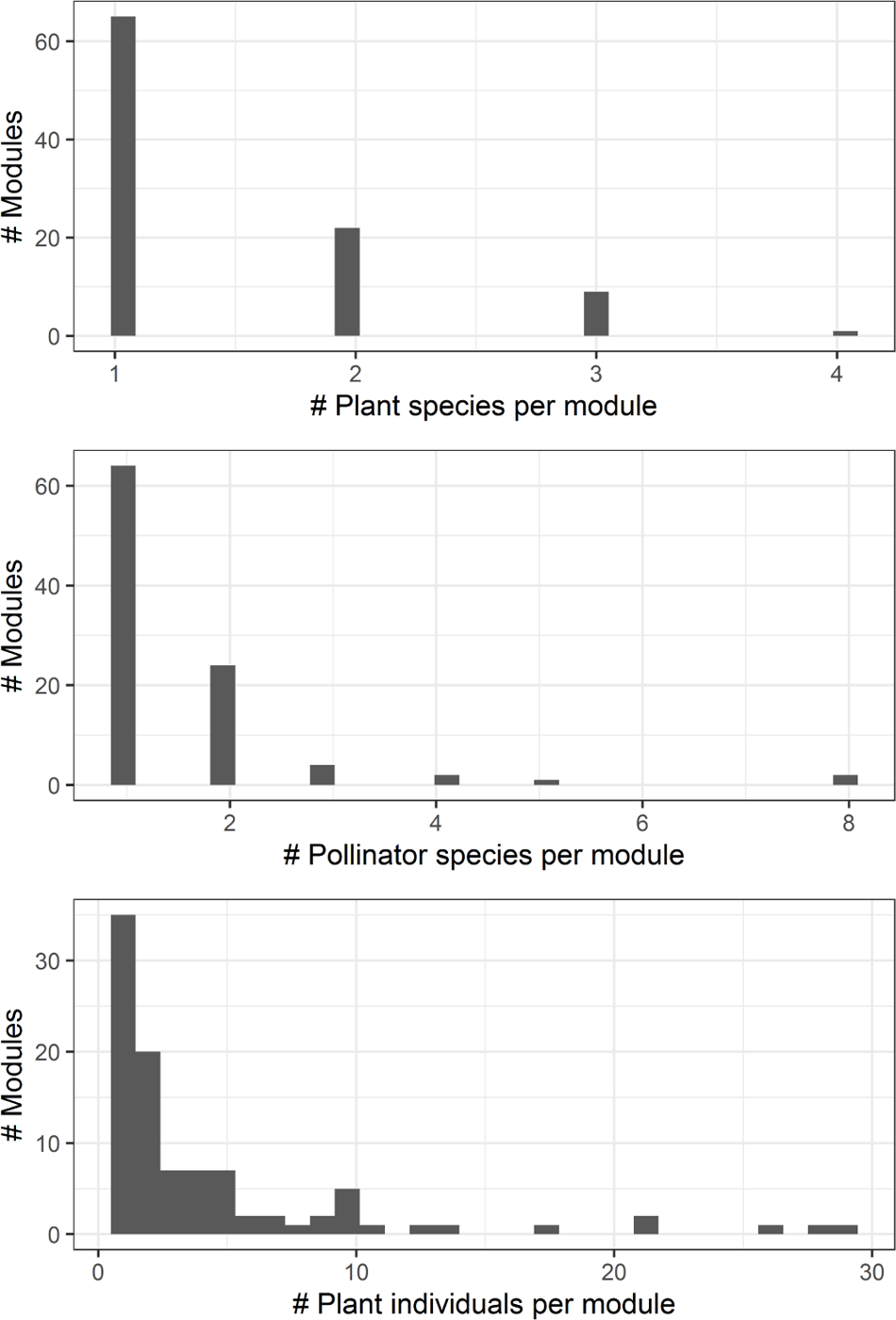
Distribution of number of plant species, insect visitors’ species, and plant individuals per module.

**Fig. A12.5:**
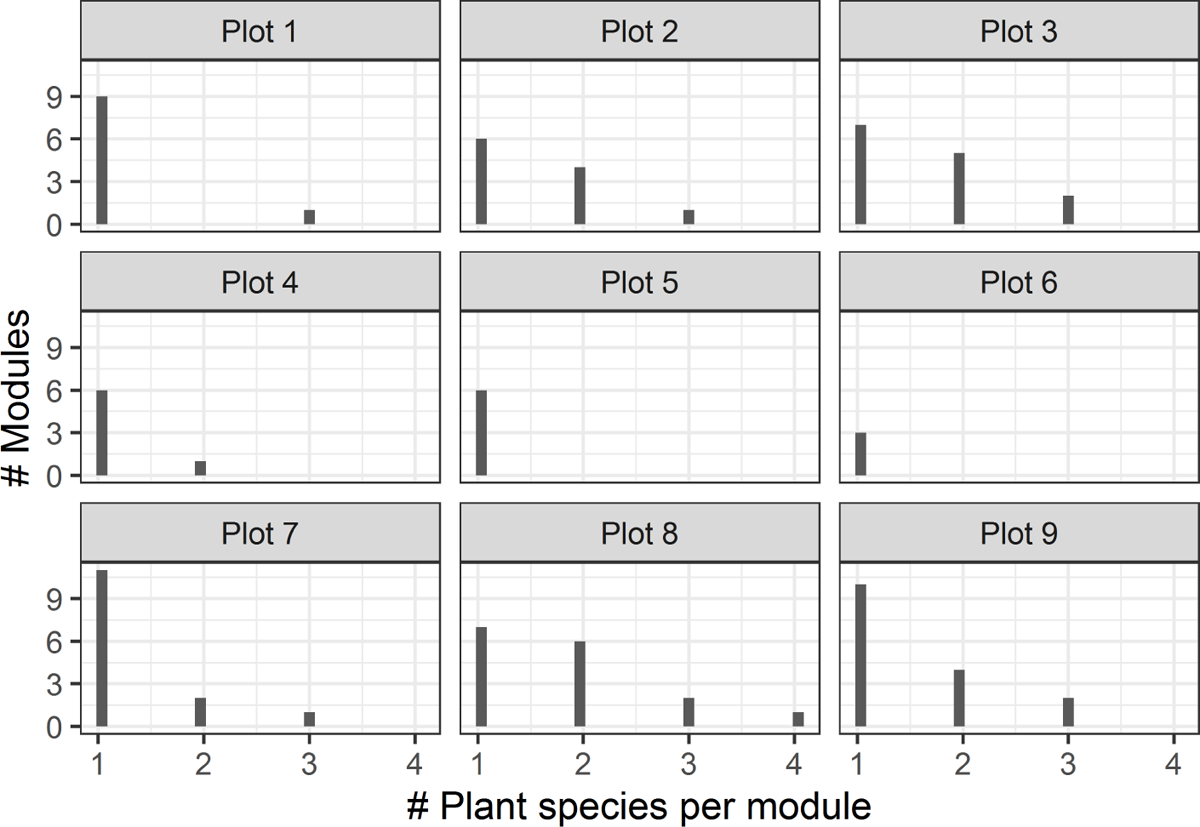
Distribution of number of plant species per module and plot.

**Fig. A12.6:**
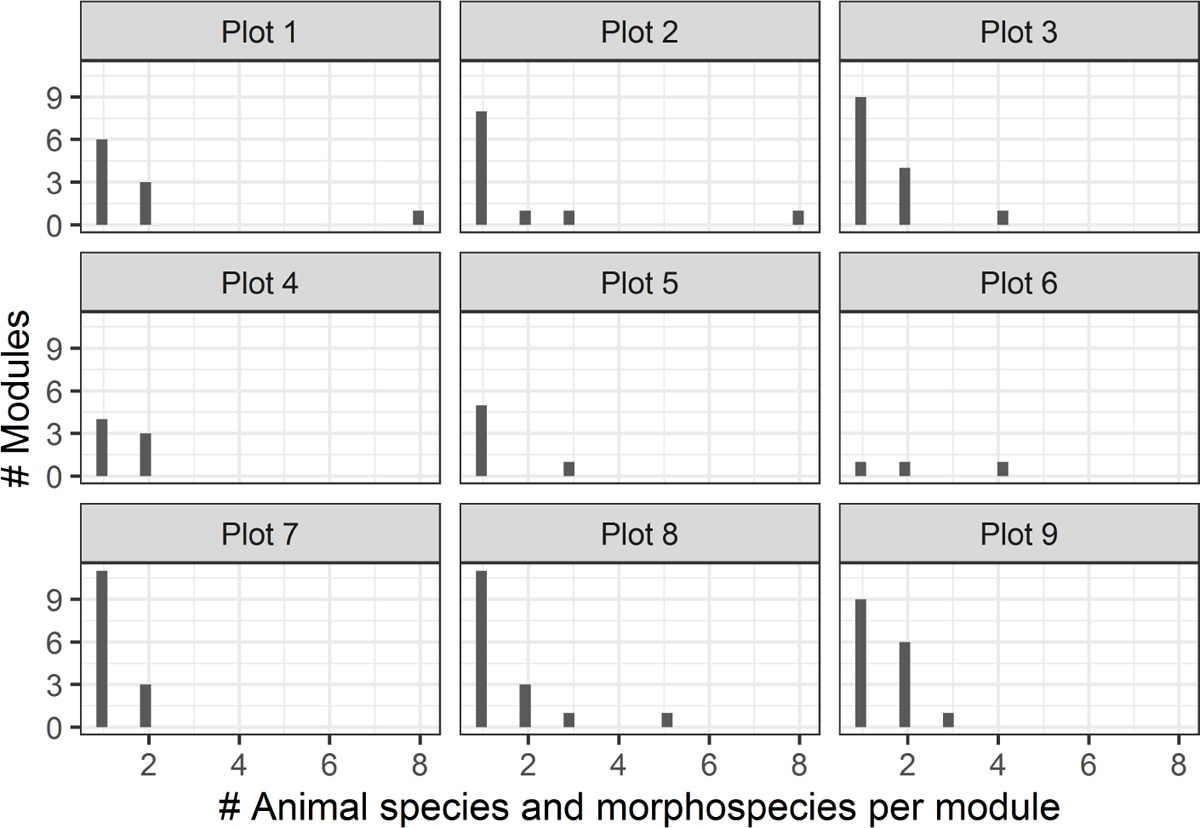
Distribution of insect visitors species per module and plot.

**Fig. A12.7:**
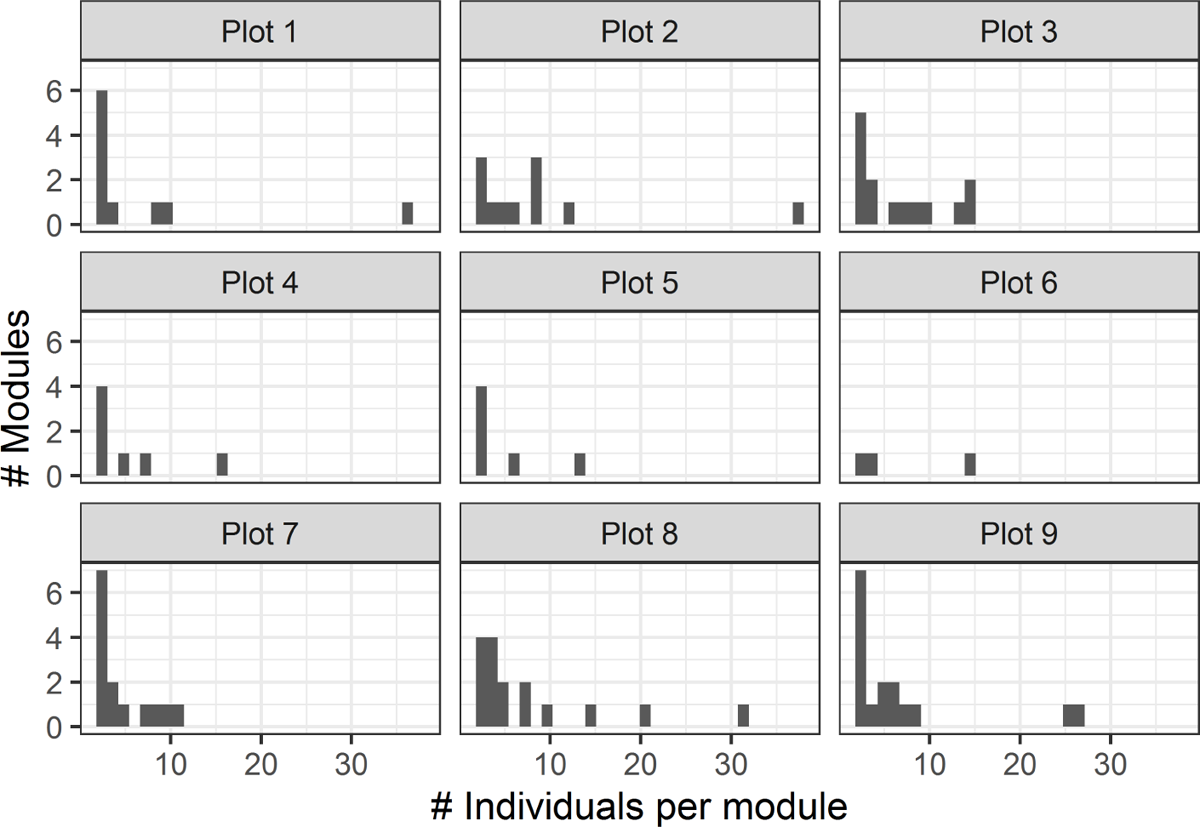
Distribution of number of plant individuals per module and plot.

**Table A12.1:** Characterization of the modules in each plot. Values in parenthesis show the results with 95% confidence intervals.

| Plot | Plant individuals<br>with visitors | Plant<br>visitors | Number of<br>modules | Plant species<br>per module | Visitors per<br>module | Plant individuals<br>per module |
| --- | --- | --- | --- | --- | --- | --- |
| 1 | 49 | 23 | 10 | 1.20<br>( 1, 3 ) | 2.00<br>( 1, 8 ) | 4.90<br>( 1, 28 ) |
| 2 | 66 | 30 | 11 | 1.55<br>( 1, 3 ) | 1.91<br>( 1, 8 ) | 6.00<br>( 1, 29 ) |
| 3 | 66 | 30 | 14 | 1.64<br>( 1, 3 ) | 1.50<br>( 1, 4 ) | 4.71<br>( 1, 13 ) |
| 4 | 26 | 11 | 7 | 1.14<br>( 1, 2 ) | 1.43<br>( 1, 2 ) | 3.71<br>( 1, 14 ) |
| 5 | 19 | 8 | 6 | 1<br>1 | 1.33<br>( 1, 3 ) | 3.17<br>( 1, 10 ) |
| 6 | 14 | 7 | 3 | 1<br>1 | 2.33<br>( 1, 4 ) | 4.67<br>( 1, 10 ) |
| 7 | 43 | 22 | 14 | 1.29<br>( 1, 3 ) | 1.21<br>( 1, 2 ) | 3.07<br>( 1, 10 ) |
| 8 | 84 | 41 | 16 | 1.81<br>( 1, 4 ) | 1.56<br>( 1, 5 ) | 5.25<br>( 1, 26 ) |
| 9 | 74 | 34 | 16 | 1.50<br>( 1, 3 ) | 1.50<br>( 1, 3 ) | 4.62<br>( 1, 21 ) |

## 13 Graph representation of the multilayers analyzed

**Fig. A13.1:**
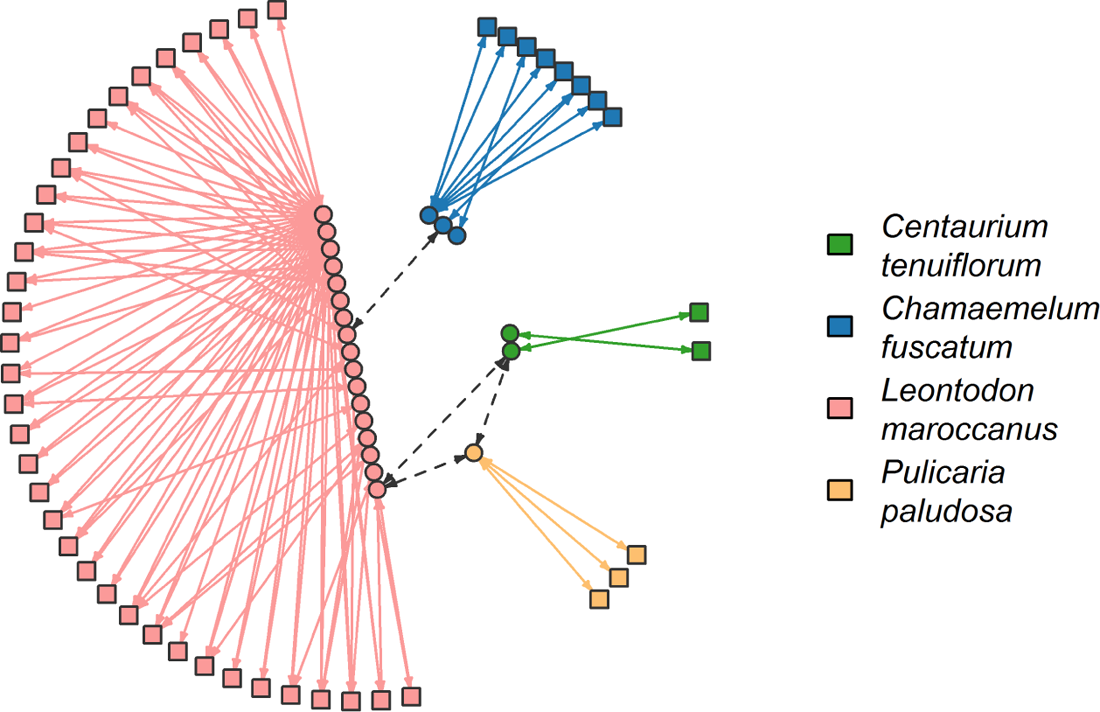
Resulting multilayer for plot 1. Symbols and lines correspond to those in Fig. 3

**Fig. A13.2:**
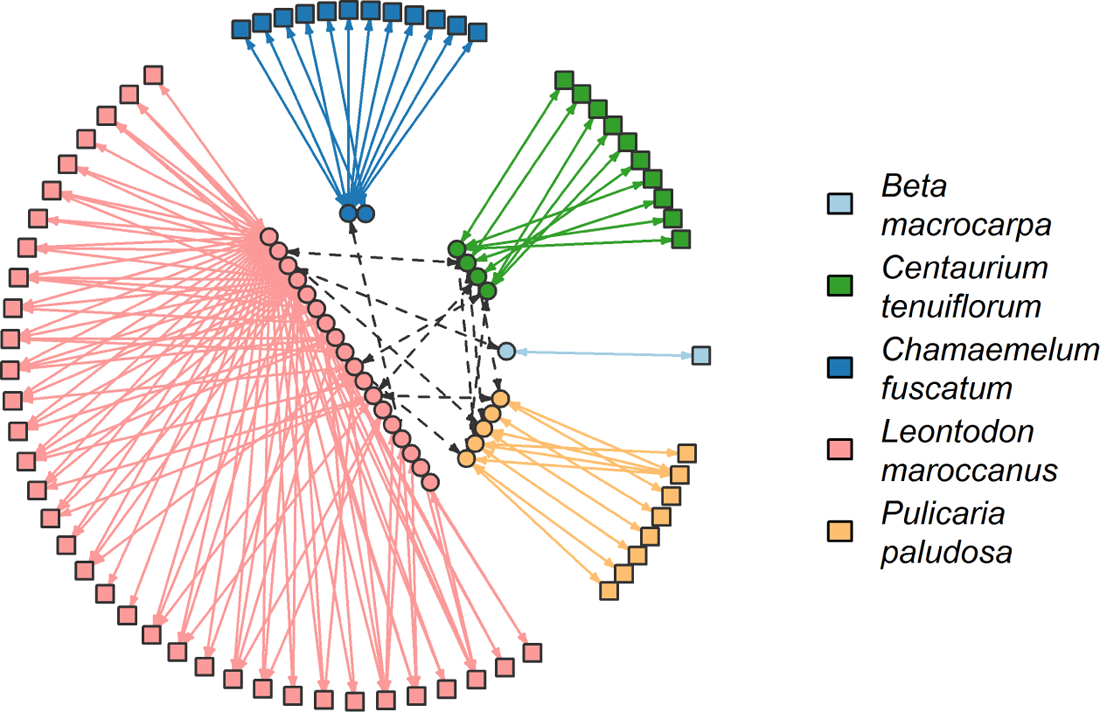
Resulting multilayer for plot 2. Symbols and lines correspond to those in Fig. 3

**Fig. A13.3:**
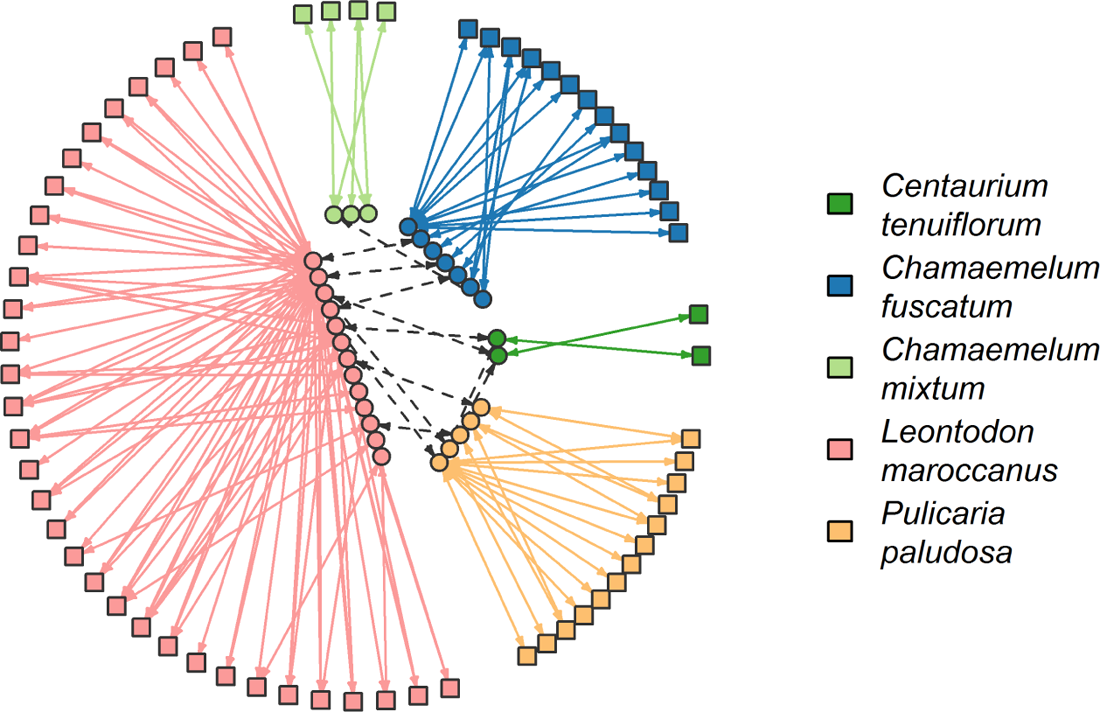
Resulting multilayer for plot 3. Symbols and lines correspond to those in Fig. 3

**Fig. A13.4:**
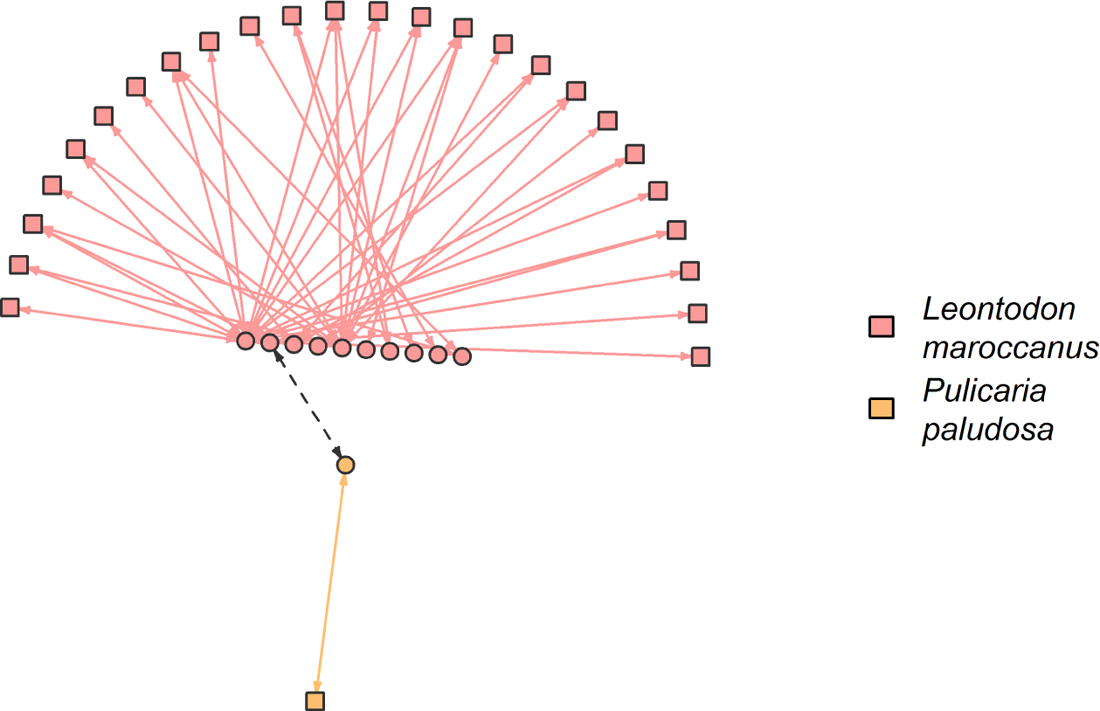
Resulting multilayer for plot 4. Symbols and lines correspond to those in Fig. 3

**Fig. A13.5:**
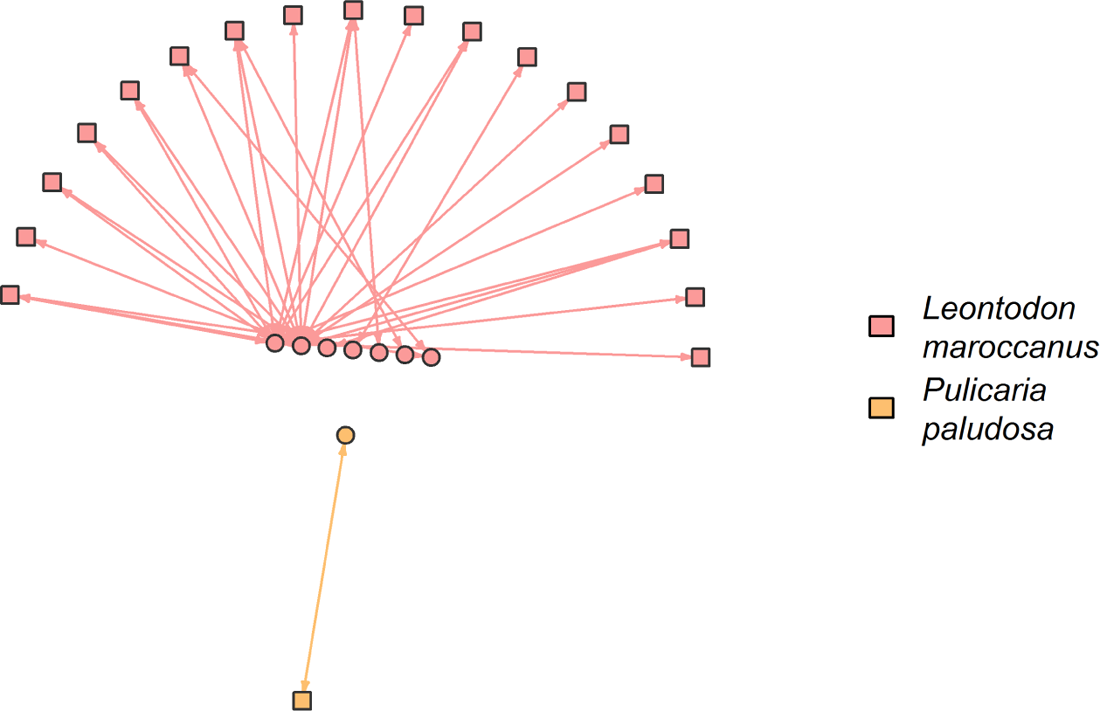
Resulting multilayer for plot 1. Symbols and lines correspond to those in Fig. 3

**Fig. A13.6:**
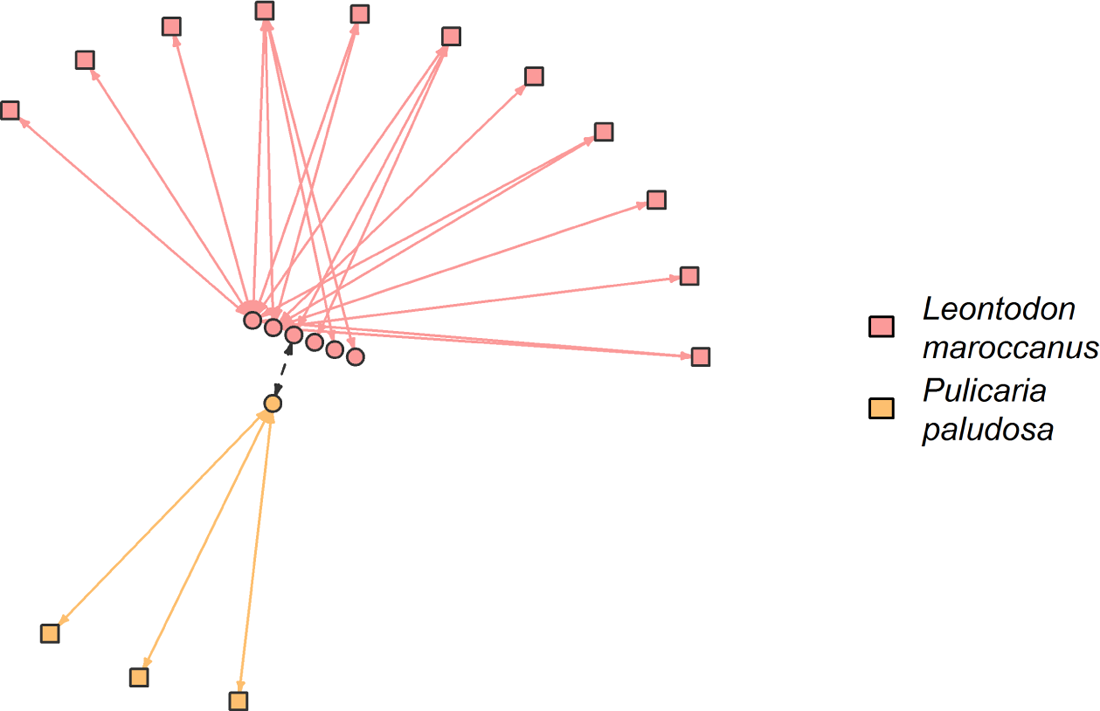
Resulting multilayer for plot 1. Symbols and lines correspond to those in Fig. 3

**Fig. A13.7:**
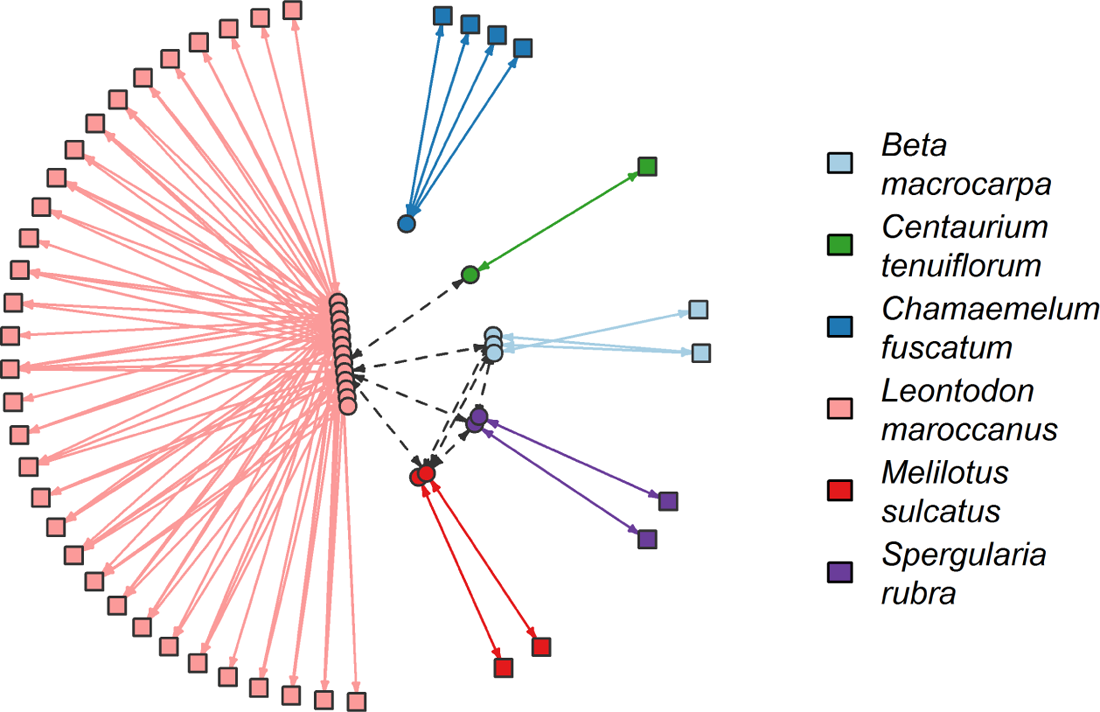
Resulting multilayer for plot 1. Symbols and lines correspond to those in Fig. 3

**Fig. A13.8:**
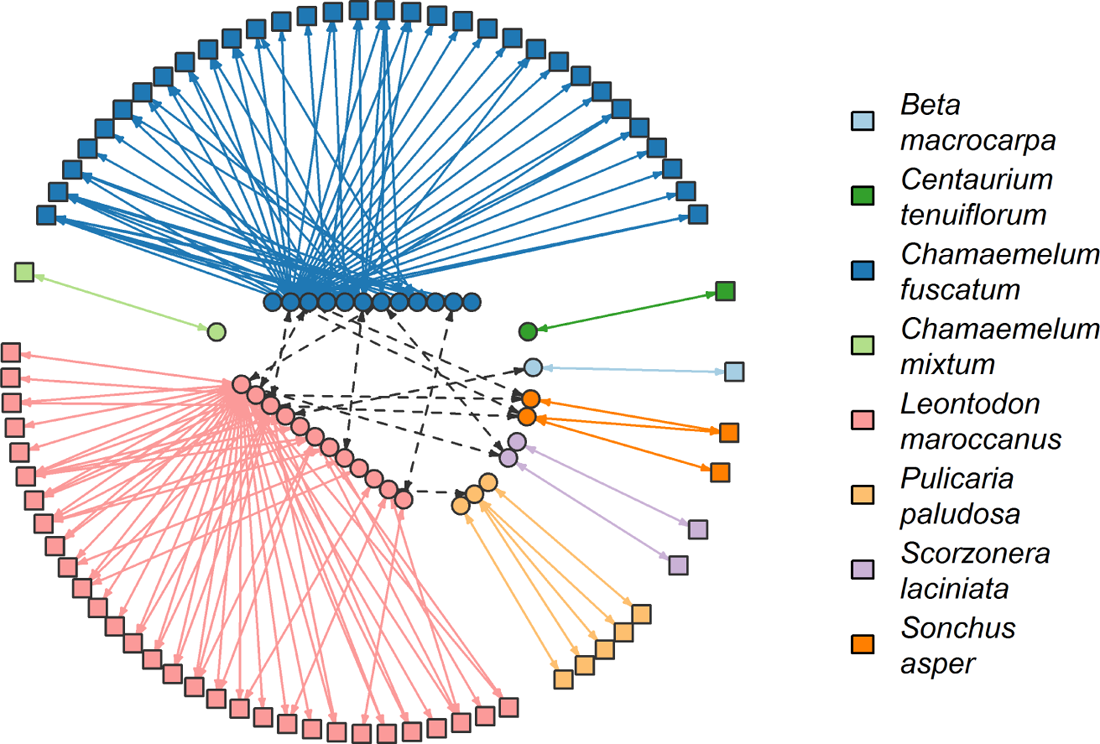
Resulting multilayer for plot 9. Symbols and lines correspond to those in Fig. 3

## 14 Specialization roles of plant individuals and floral visitors

Plant and pollinator nodes within a module may exert a different impact on the information flow that they trap. For instance, depending on their centrality metrics, nodes may hoard the pollen influx and/or outflux. Besides, in the case of insect species, those that connect several plant species and modules can be responsible for distributing high loads of heterospecific pollen. To assess the roles of plant and pollinator nodes in each multilayer, we classified them in the following categories: (i) peripherals, i.e., vertices linked to very few nodes that usually are within their module; (ii) module hubs, i.e., highly connected vertices linked to many nodes within their own module; (iii) connectors, i.e., nodes that link several modules; and (iv) network hubs, i.e., nodes acting as both connectors and module hubs (Olesen *et al*., 2007). To do so, we calculated their weighted standard among module connectivity and within module degree, denoted by c and z, respectively (Dormann & Strauss, 2013; Olesen *et al*., 2007). In unweighted and undirected networks, the former represents the level to which a given node is linked to other modules, and the latter is its standardized number of links to other species in the same module (Guimerà & Amaral, 2005). Since our multilayers are weighted and directed, we must distinguish between the links that arrive at a node (or in-links) and those that depart from it (or out-links). For that reason, we adapted the previous definitions and computed two values of *c* (namely: in- and out-*c*) and two of *z* (in- and out-*z*) by using in- and out-strength of each node, which correspond to the sum of the weights of its in-links and the one for its out-links, respectively. From the in-links (out-links) perspective, we assigned the following roles to nodes: peripherals [low in-(out-)*c* and low in-(out-)*z*], module hubs [low in-(out-)*c* and high in-(out-)*z*], connectors [high in-(out-)*c* and low in-(out-)*z*], and network hubs [high in-(out-)*c* and high in-(out-)*z*].

To define the thresholds for in/out-*c*- and in/out-*z*-values, we followed Watts *et al*. (2016) and used the 95% quantiles of such metrics obtained from a null model for our original multilayer networks. Specifically we generated an ensemble of 100 null multilayers for each plot, respectively, and we calculated in/out-*c*- and in/out-*z*-values for each randomized multilayer. Our null model preserves the total number of observed interactions per pollinator and assumes that the weight of interlinks are those of the observed multilayers. We estimated the thresholds for in/out-*c* and in/out-*z* over all the plots (i.e., over the results for 900 null multilayers), and their values are 1.00/1.00 and 1.93/1.62, respectively.

By using the above thresholds, overall, around 30 - 40% of the plant and insect nodes were peripherals (with all their in- and out-links outside their own module), 40 - 60% were connectors among modules, 2.5 - 10% of insect species acted as module hubs, and 2.5 - 10% of insect species were network hubs. Since insect species channel all the links between conspecific and heterospecific plant nodes (see Fig. 1(e)), the observed absence of plant hubs is expected. In addition, our results also confirmed that most network and module hubs belong to the main insect visitors of the most visited plant species, namely: flower beetles, humbleflies, small beetles, bees. Specific details are as follows. From the in-link (or pollen influx to nodes) perspective, we found that 37.87% of all plant nodes and 29.61% of all floral visitors were peripherals (with c greater than 0, i.e., they all have in-links outside their own module), 58.50% of plant nodes and 50.00% of insect visitors were connectors among modules, 9.71% of insect visitors were module hubs, whereas 2.91% of insects were network hubs (Figs. A14.1 and A14.2). Influx roles for the remaining plant nodes (3.63%) and insect visitors (7.77%) can not be computed because the standard deviation of their respective within-layer centralities is equal to zero, and consequently their within module instrength (i.e. in-z), which is defined in multiples of the within-layer centralities’ standard deviations Dormann & Strauss (2013); Olesen *et al*. (2007)(Olesen et al. 2007, Dormann et al. 2009), can not be properly assessed. Influx network hubs were *Lasioglossum malachurum* (bee) in *P. paludosa* (plot 8), Ulidiidae (small flies) in *C. fuscatum* (plots 1, 2 and 3), *Psilothrix viridicoerulea* (flower beetles) in *L. maroccanus* layers (plot 6), and *Bombylius sp.* (humbleflies) in *C. tenuiflorum* (plot 2), whereas module hubs were *Anastoechus sp.* (humbleflies) in *L. maroccanus* (plot 3), *Andrena humilis* (solitary bees) in *L. maroccanus* (plot 7), *Andrena sp.* (solitary bees) in *C. fuscatum* (plot 9), *Brassicogethes sp.* (small beetles) in *L. maroccanus* (all plots), *Lasioglossum malachurum* (bee) in *P. paludosa* (plot 3), *Psilothrix viridicoerulea* (flower beetles) in *C. fuscatum* (plot 9) and *L. maroccanus* (plots 3, 4, 5, 7, and 8), and Ulidiidae (small flies) in *C. fuscatum* (plot 8). Results for the out-links (or pollen outflux from nodes) show that 37.87% of all plant nodes and 34.47% of all floral visitors were peripherals (with c greater than 0), 51.25% of plant nodes and 37.86% of insect visitors were connectors among modules, 2.91% of insect visitors were module hubs, whereas only 0.45% of plant nodes and 10.68% of insects were network hubs (see Figs. A14.3 and A14.4). Outflux roles for 10.88% of plant nodes and 14.08% of insect visitors are omitted. Outflux network hubs were *Anastoechus sp.* (humbleflies) in layer *C. tenuiflorum* (plot 2), *Andrena humilis* (solitary bees) in *S. laciniata* (plot 8), *Bombilus major* (humbleflies) in *P. paludosa* (plot 3), *Brassicogethes sp.* (Small beetles) in *C. fuscatum* (plots 3 and 9), *P. paludosa* (plot 8) and *S. laciniata* (in plot 8 and 9, respectively); *Empis sp.* (small flies) in *C. fuscatum* (plot 1); *Lasioglossum malachurum* (bees) in *C. tenuiflorum* (2 and 3), *M. sulcatus* (plot 8), and *P. paludosa* (1, 4, 8, and 9); Mordellidae (flower beetles) in *S. asper* (plot 9); *Musca sp.* (house flies) in *P. paludosa* (plot 8); *Psilothrix viridicoerulea* (flower beetles) in *C. fuscatum* (plot 3) and *P. paludosa* layers (plot 8); *Sphaerophoria scripta* (hoverflies) in *M. sulcatus* (plot 7); and Ulidiidae (Small flies) in *C. fuscatum* (plot 2), whereas module hubs are *Anastoechus sp.* (humbleflies) in *P. paludosa* (plot 3); *Bombylius major* (humbleflies) in *P. paludosa* (plot 2); *Brassicogethes sp.* (small beetles) in *L. maroccanus* (plot 9); *Lasioglossum malachurum* (solitary bees) in *P. paludosa* (plot 3); and *Psilothrix viridicoerulea* (flower beetles) in *C. fuscatum* and *S. asper* (both in plot 9).

**Fig. A14.1:**
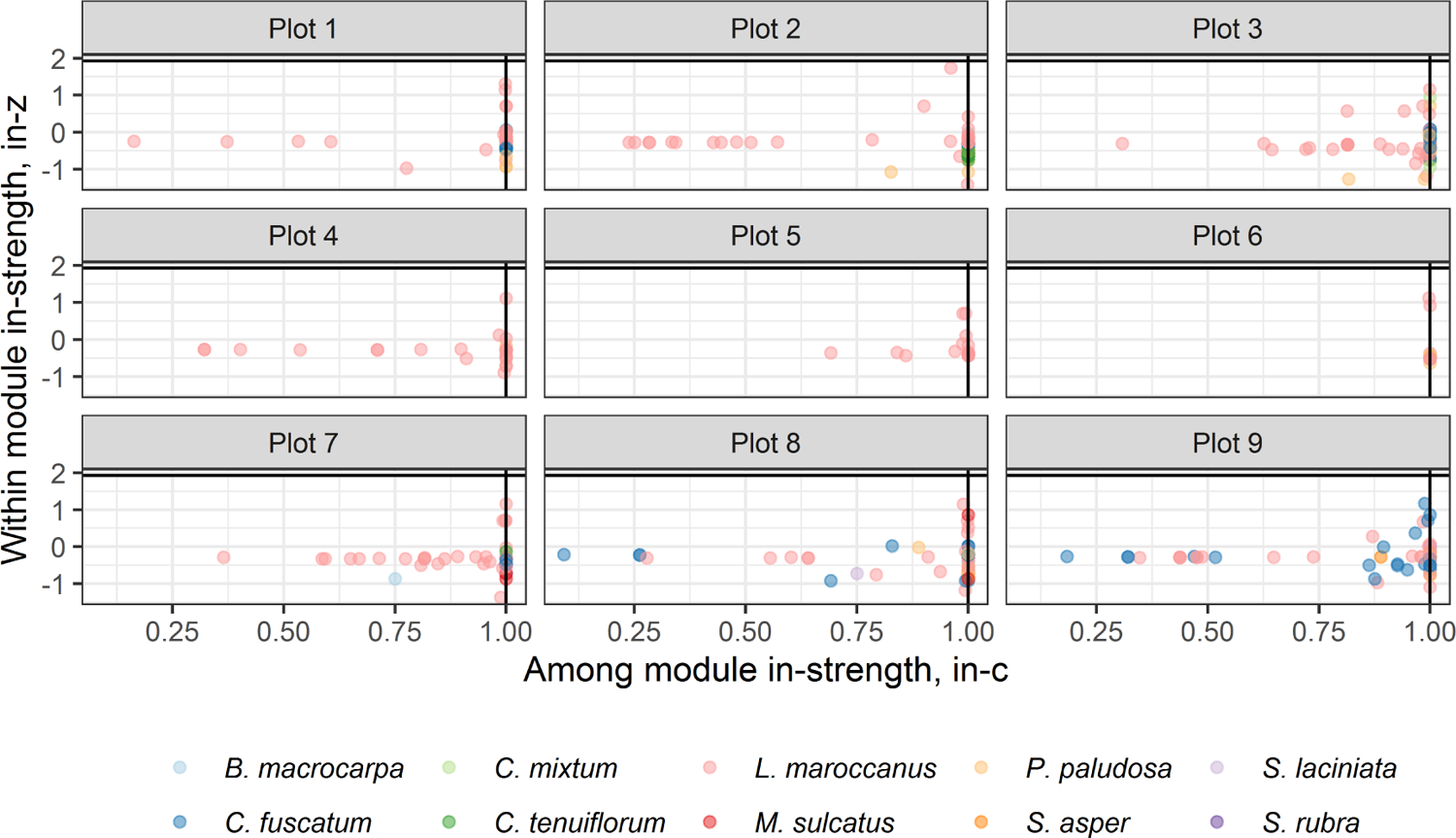
Roles performed by plant nodes in the plot multilayers according to within module in-strength (in-z) and among module in-strength (in-c). Lines at in-z = 1.93 and in-c = 1.00 define species roles.

**Fig. A14.2:**
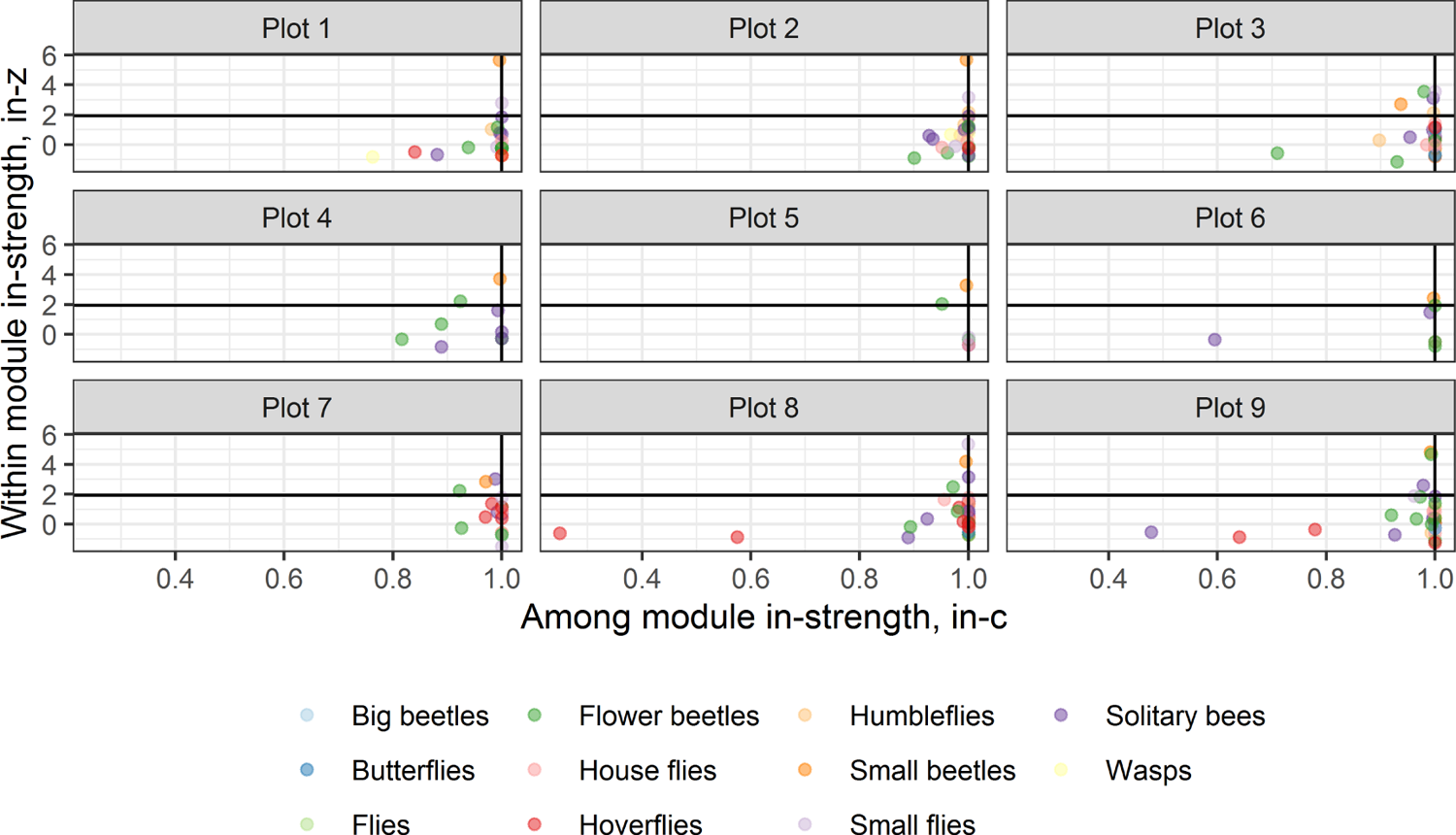
Roles performed by floral visitors in the plot multilayers according to within module in-strength (in-z) and among module in-strength (in-c). Lines at in-z = 1.93 and in-c = 1.00 define species roles.

**Fig. A14.3:**
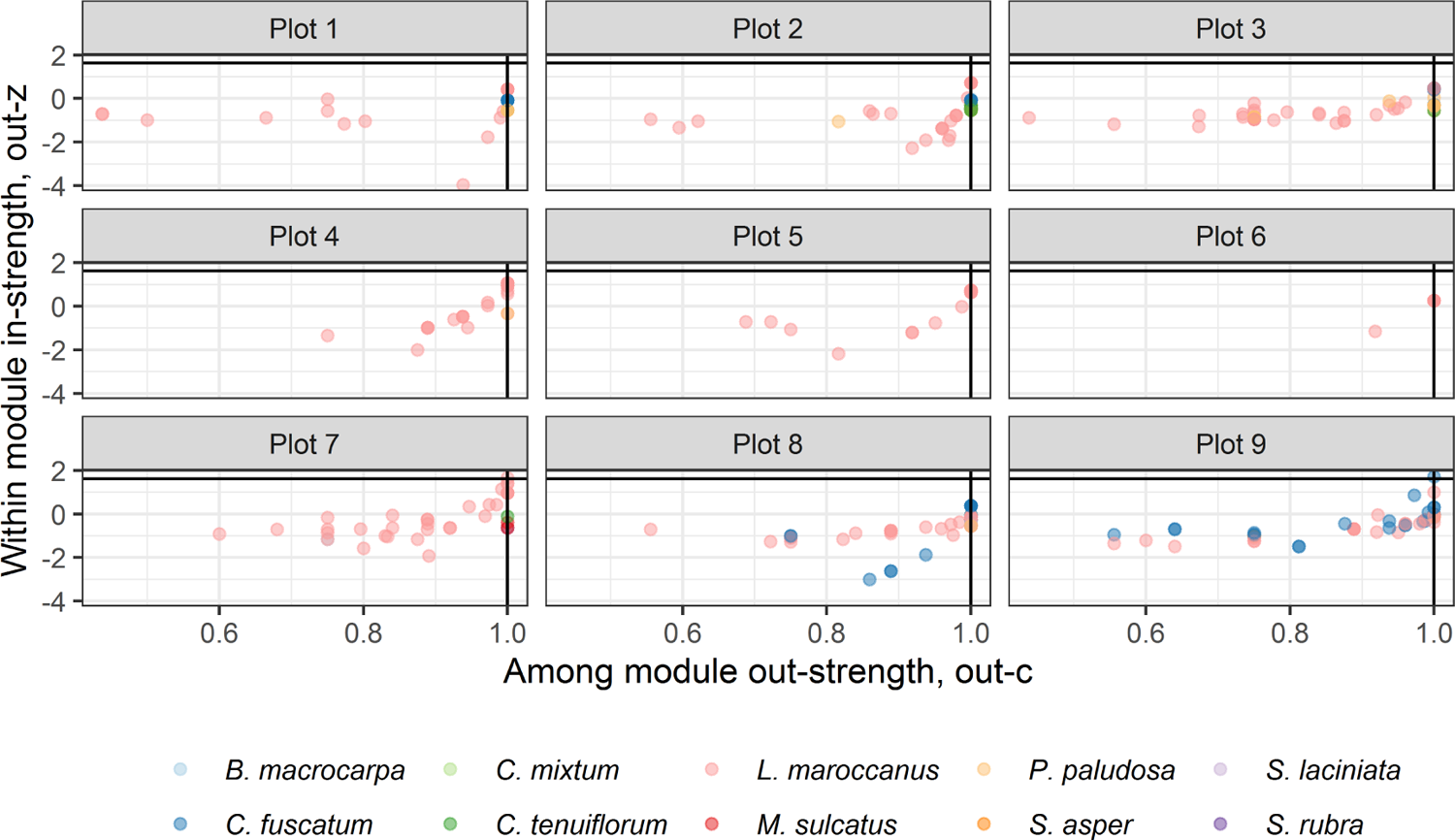
Roles performed by plant nodes in the plot multilayers according to within module out-strength (out-z) and among module out-strength (out-c). Lines at out-z = 1.62 and out-c = 1.00 define species roles.

**Fig. A14.4:**
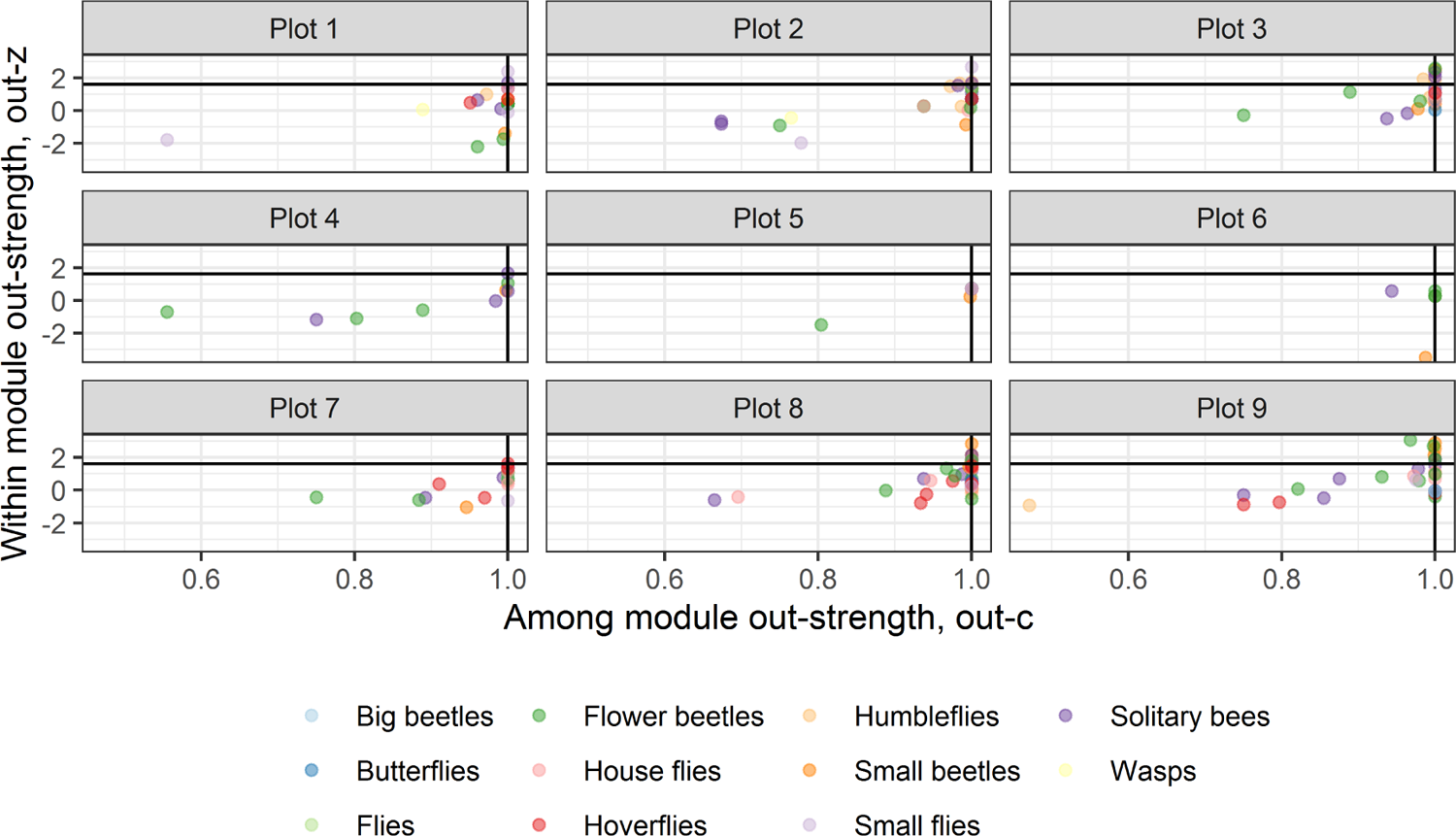
Roles performed by floral visitors in the plot multilayers according to within module out-strength (out-z) and among module out-strength (out-c). Lines at out-z = 1.62 and out-c = 1.00 define species roles.

## 15 Testing significance of modularity in Caracoles Ranch (2020)

In this appendix we test the significance of modularity in our empirical multilayers by using the null models explained above in appendix 3. In particular, we derive a null distribution of map equation and number of modules for each null mode. To do so, we generated an ensemble of 500 random multilayer for each plot and null model (500 *×* 9 *×* 2 = 9, 000 random multilayers in total).

For each randomized multilayer, we have calculated the map equation (*L_Simulated_*) and number of modules (*m_Simulated_*). Then, we estimated the confidence interval for each null distribution and tested whether the observed values of our metrics belong to such intervals (non-significant result) or not (significant result). Our results are summarized in tables A15.1 and A15.2, and Figs. A15.1, A15.2, A15.3 and A15.4. As can be seen, in each plot *L_Observed_*is significantly smaller than the results for randomized multilayer with a confidence level of 95%. Consequently, observed network flows tend to be more constrained within modules than in any of the random multilayers. On the other hand, we found no significant differences between *m_Observed_* and the number of modules of randomized multilayers with a confidence level of 95%.

**Fig. A15.1:**
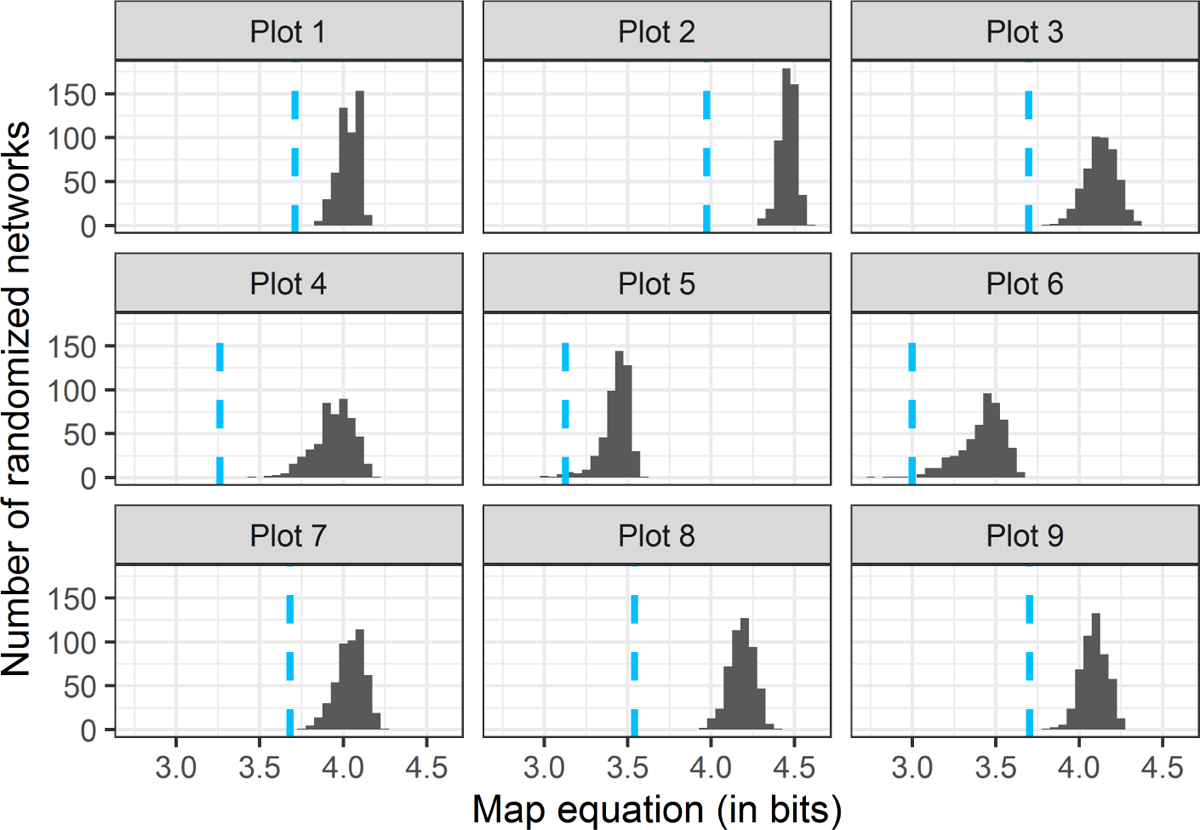
Null distribution of map equation (in bits) obtained from 500 simulations for each plot, when the null model 1 in appendix 3 is considered. Each panel represents the results for a plot. The respective observed values are marked with vertical blue dashed lines.

**Fig. A15.2:**
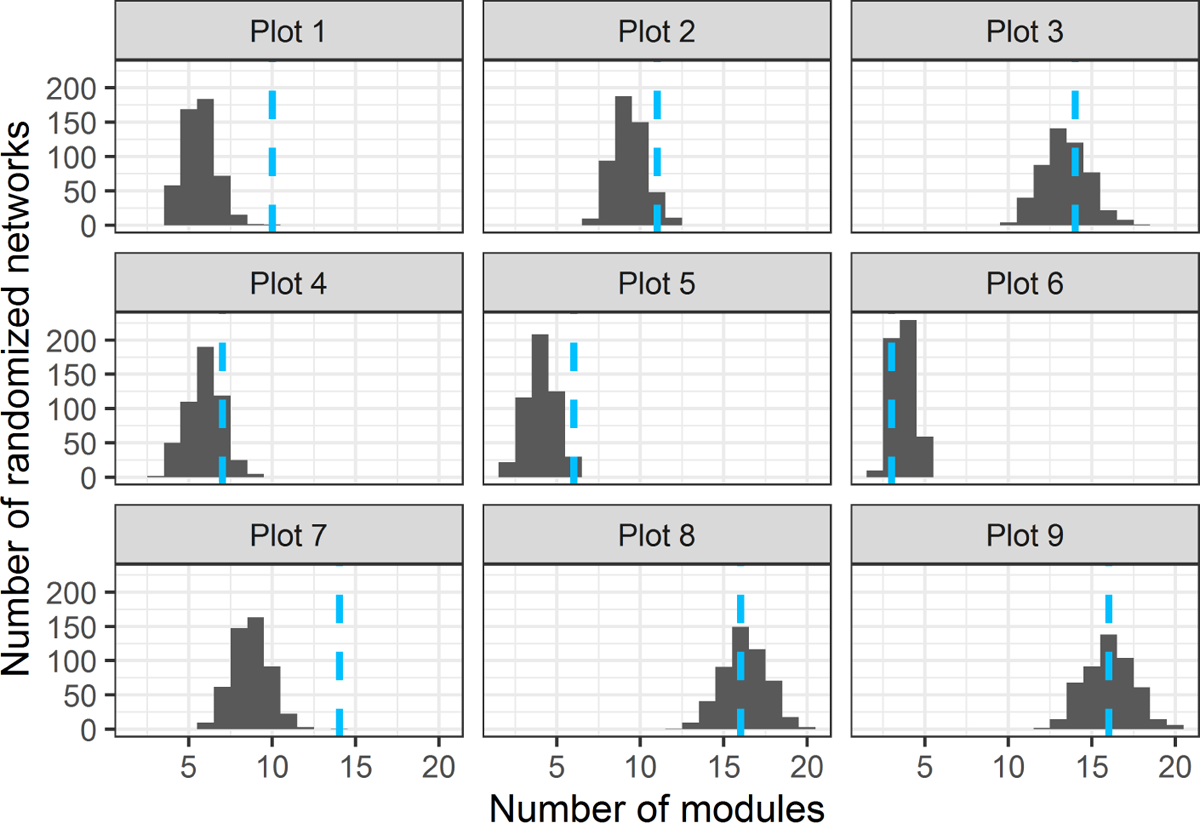
Null distribution of the number of modules obtained from 500 simulations for each plot, when the null model 1 in appendix 3 is considered. Each panel represents the results of a plot. The respective observed values are marked with vertical blue dashed lines.

**Fig. A15.3:**
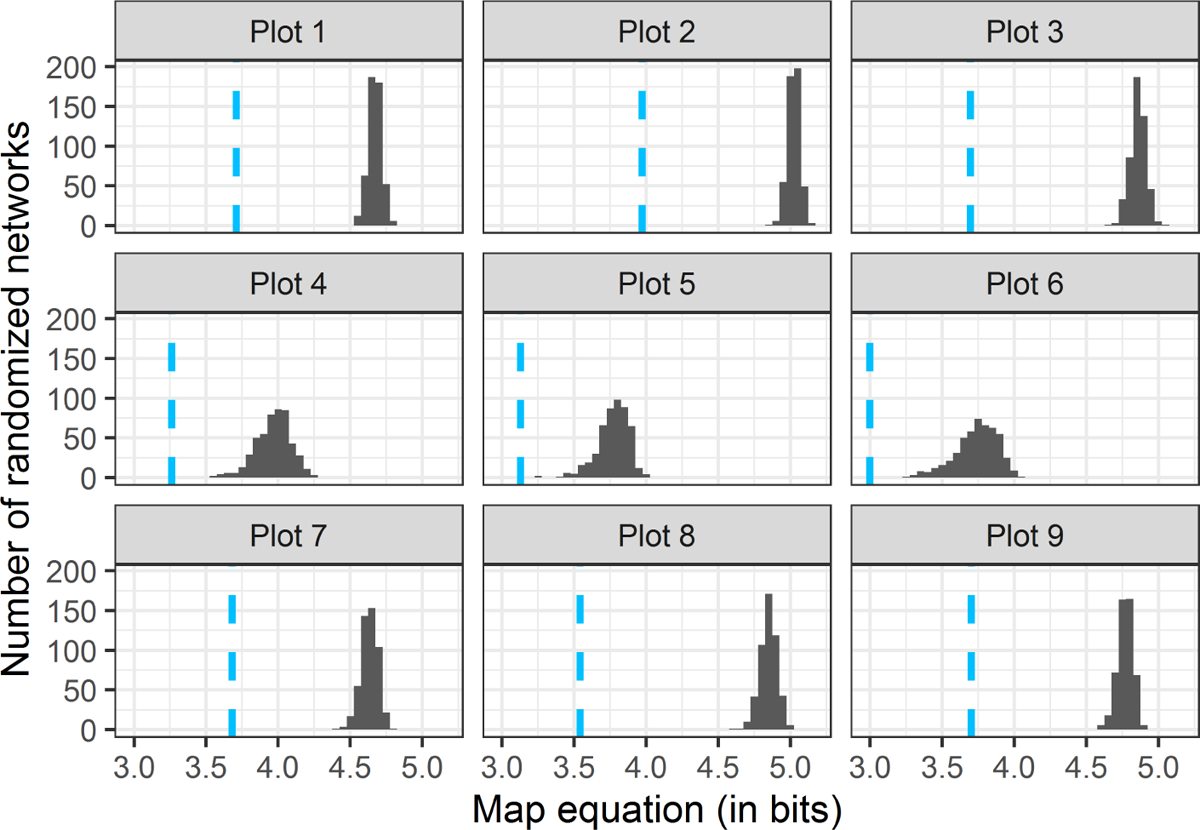
Null distribution of map equation (in bits) obtained from 500 simulations for each plot by using Null model 2 in appendix 3. Each panel represents the results of a plot. The respective observed values are marked with vertical blue dashed lines.

**Fig. A15.4:**
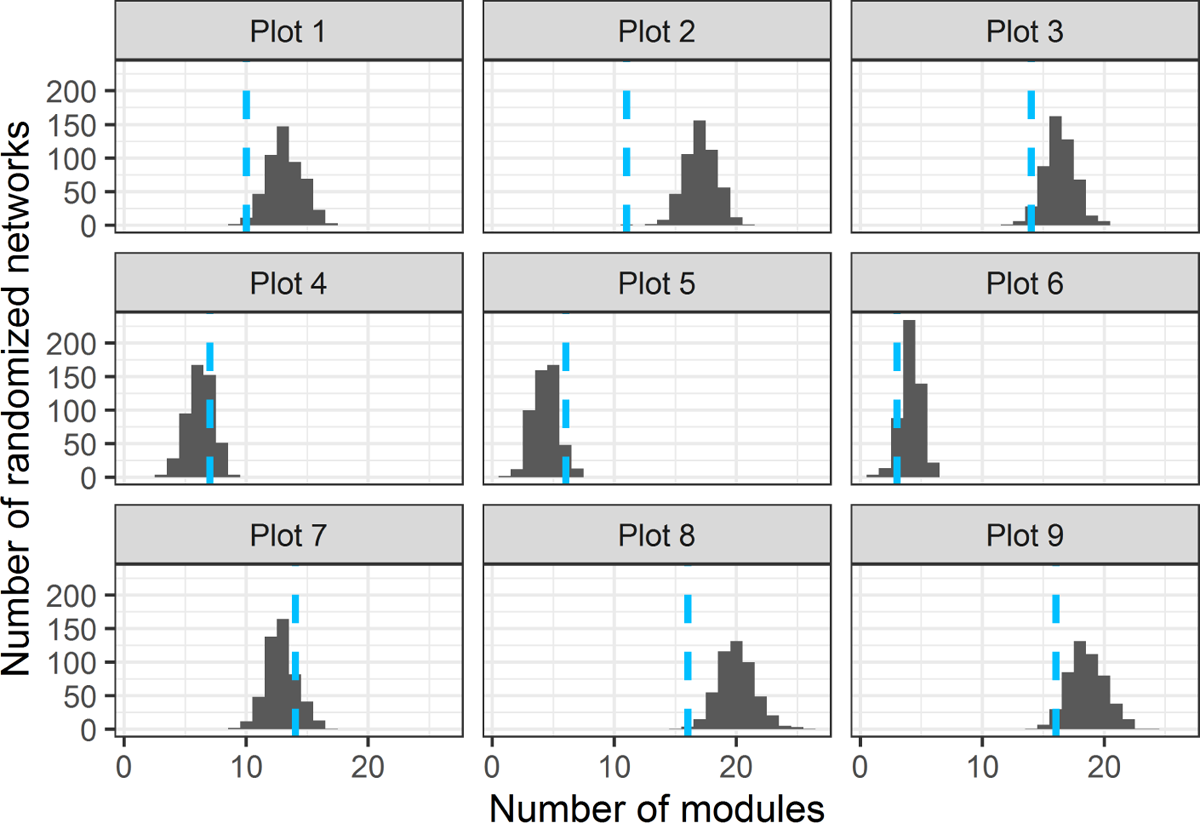
Null distribution of the number of modules obtained from 500 simulations for each plot by using Null model 2 in appendix 3. Each panel represents the results of a plot. The respective observed values are marked with vertical blue dashed lines.

**Table A15.1:** Observed map equation (*L_Observed_*) and number of modules (*m_Observed_*) for each plot, and the corresponding confidence intervals (C.I.) for the simulated map equation (*L_Simulated_*) and simulated number of modules (*m_Simulated_*), when the null model 1 in appendix 3 is considered. Null distributions for *L_Simulated_* and *m_Simulated_* are shown in Figs.A15.1 and A15.2. Bold values denote significant results.

NULL MODEL 1
| Plot | $L_{Observed}$ | C. I. for $L_{Simulated}$ | $m_{Observed}$ | C. I. for $m_{Simulated}$ |
| --- | --- | --- | --- | --- |
| 1 | **3.709** | ( **3.908**, **4.126** ) | **10** | ( 4, 8 ) |
| 2 | **3.971** | ( **4.367**, **4.553** ) | 11 | ( 8, 11 ) |
| 3 | **3.696** | ( **3.943**, **4.297** ) | 14 | (11, 16 ) |
| 4 | **3.259** | ( **3.678**, **4.141** ) | 7 | ( 4, 8 ) |
| 5 | **3.126** | ( **3.175**, **3.538** ) | 6 | ( 2, 6 ) |
| 6 | **2.997** | ( **3.093**, **3.611** ) | 3 | ( 3, 5 ) |
| 7 | **3.680** | ( **3.851**, **4.189** ) | **14** | ( 7, 11 ) |
| 8 | **3.539** | ( **4.019**, **4.315** ) | 16 | (14, 19 ) |
| 9 | **3.701** | ( **3.940**, **4.230** ) | 16 | (13, 19 ) |

**Table A15.2:**
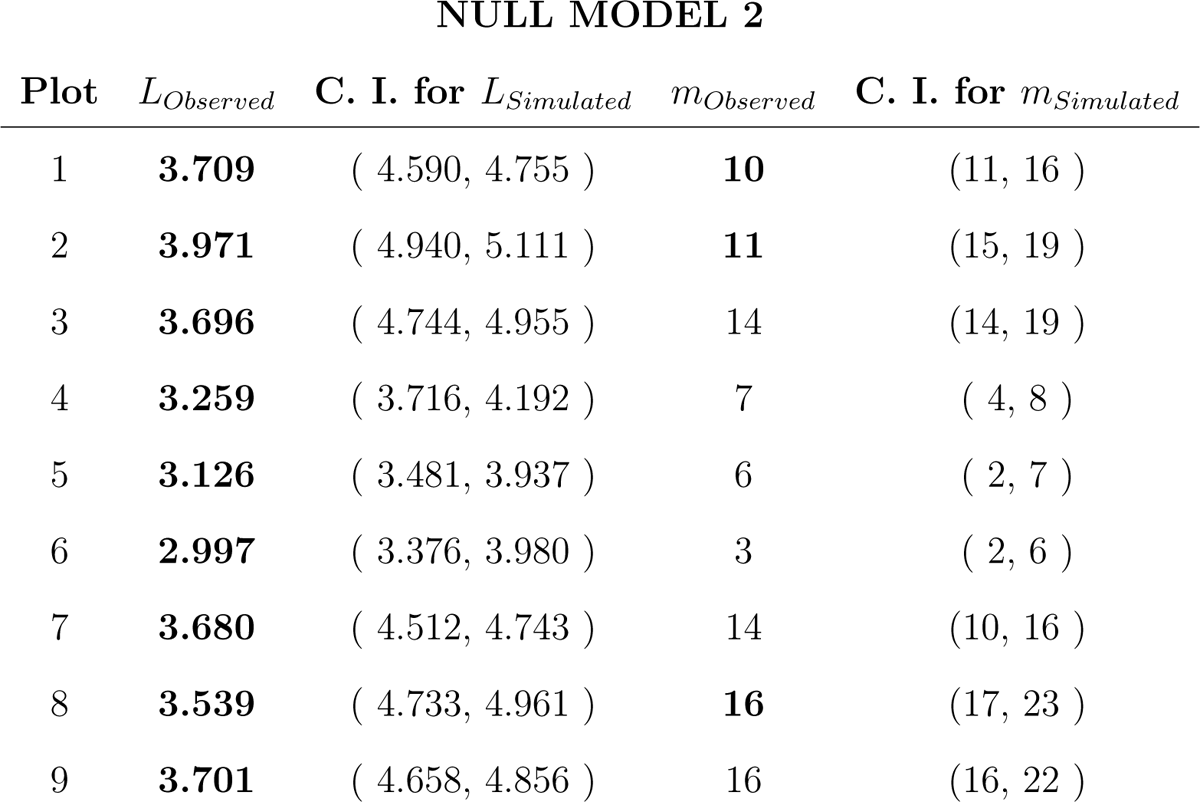
Observed map equation (*L_Observed_*) and number of modules (*m_Observed_*) for each plot, and the corresponding confidence intervals (C.I.) for the simulated map equation (*L_Simulated_*) and simulated number of modules (*m_Simulated_*), when the null model 2 in appendix 3 is considered. Null distributions for *L_Simulated_* and *m_Simulated_* are shown in Figs. A15.3 and A15.4. Bold values denote significant results.

NULL MODEL 2
| Plot | $L_{Observed}$ | C. I. for $L_{Simulated}$ | $m_{Observed}$ | C. I. for $m_{Simulated}$ |
| --- | --- | --- | --- | --- |
| 1 | **3.709** | ( 4.590, 4.755 ) | **10** | (11, 16 ) |
| 2 | **3.971** | ( 4.940, 5.111 ) | **11** | (15, 19 ) |
| 3 | **3.696** | ( 4.744, 4.955 ) | 14 | (14, 19 ) |
| 4 | **3.259** | ( 3.716, 4.192 ) | 7 | ( 4, 8 ) |
| 5 | **3.126** | ( 3.481, 3.937 ) | 6 | ( 2, 7 ) |
| 6 | **2.997** | ( 3.376, 3.980 ) | 3 | ( 2, 6 ) |
| 7 | **3.680** | ( 4.512, 4.743 ) | 14 | (10, 16 ) |
| 8 | **3.539** | ( 4.733, 4.961 ) | **16** | (17, 23 ) |
| 9 | **3.701** | ( 4.658, 4.856 ) | 16 | (16, 22 ) |

## 16 Testing significance of subgraphs in Caracoles Ranch (2020)

In this appendix we test the significance of subgraphs in our empirical multilayers by using the null model 1 explained above, in appendix 3. In particular, we generated 500 random multilayer for each plot (4,500 random multilayers in total). Our findings are summarized in Figs. A16.1 and A16.2. According to such figures, 46.71% (82.09%) of plant individuals have a total amount of homospecific (heterospecific) motifs that is significantly different from the values obtained for randomized systems, with a confidence level of 95%. In the case of homospecific motifs, it is possible to see that, as expected, plant species with few (many) plant individuals tend to exhibit a higher (lower) proportion of significant results, specifically a significantly larger (smaller) number of homospecific motifs. Finally, it is worth mentioning that 97.27% of individual plants with significant heterospecific motifs exhibit larger values of such motifs than the randomized arrangements.

**Fig. A16.1:**
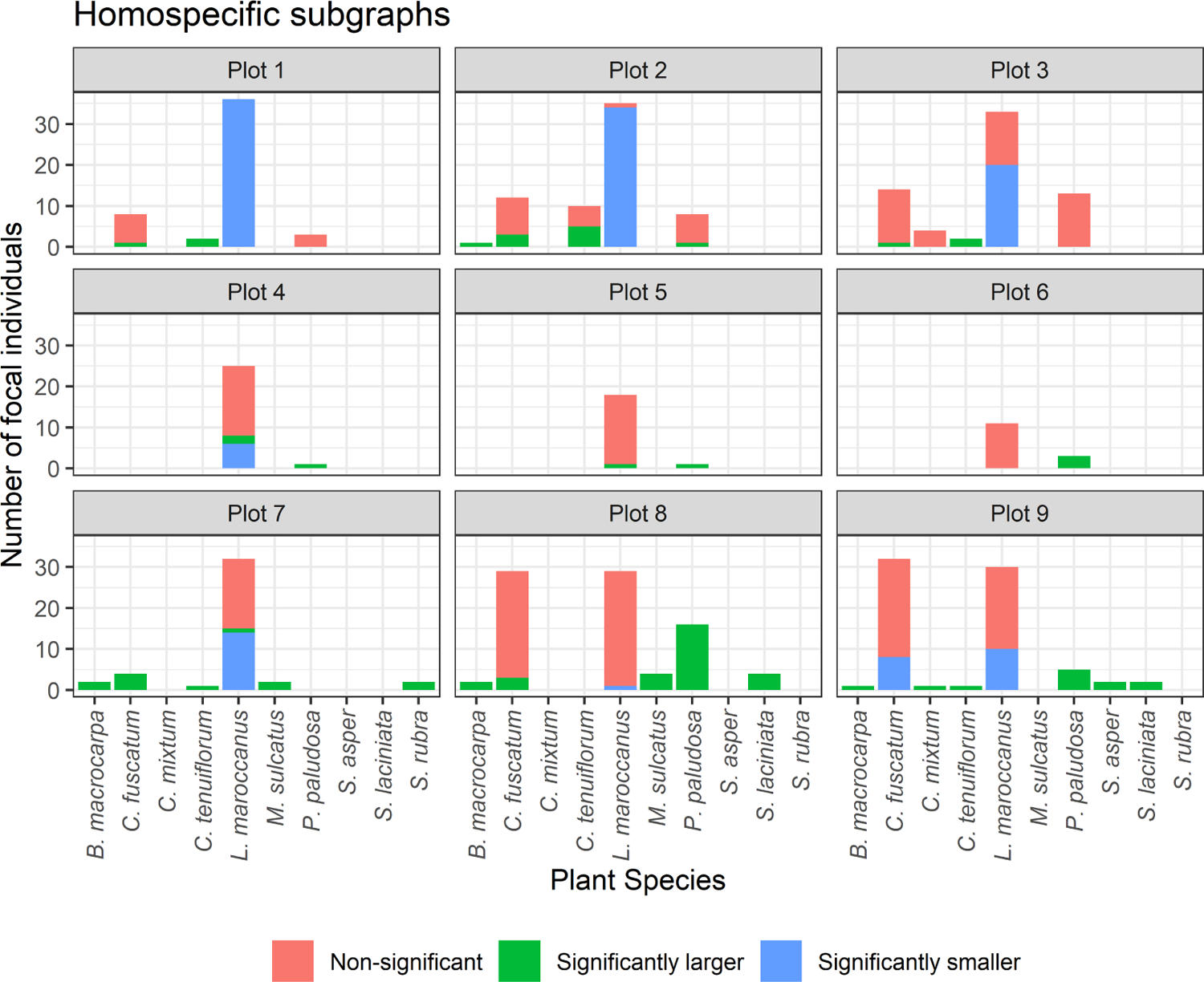
Number of plant individuals with significant and non-significant homospecific motifs per plot and plant species, when the null model 1 in appendix 3 is considered.

**Fig. A16.2:**
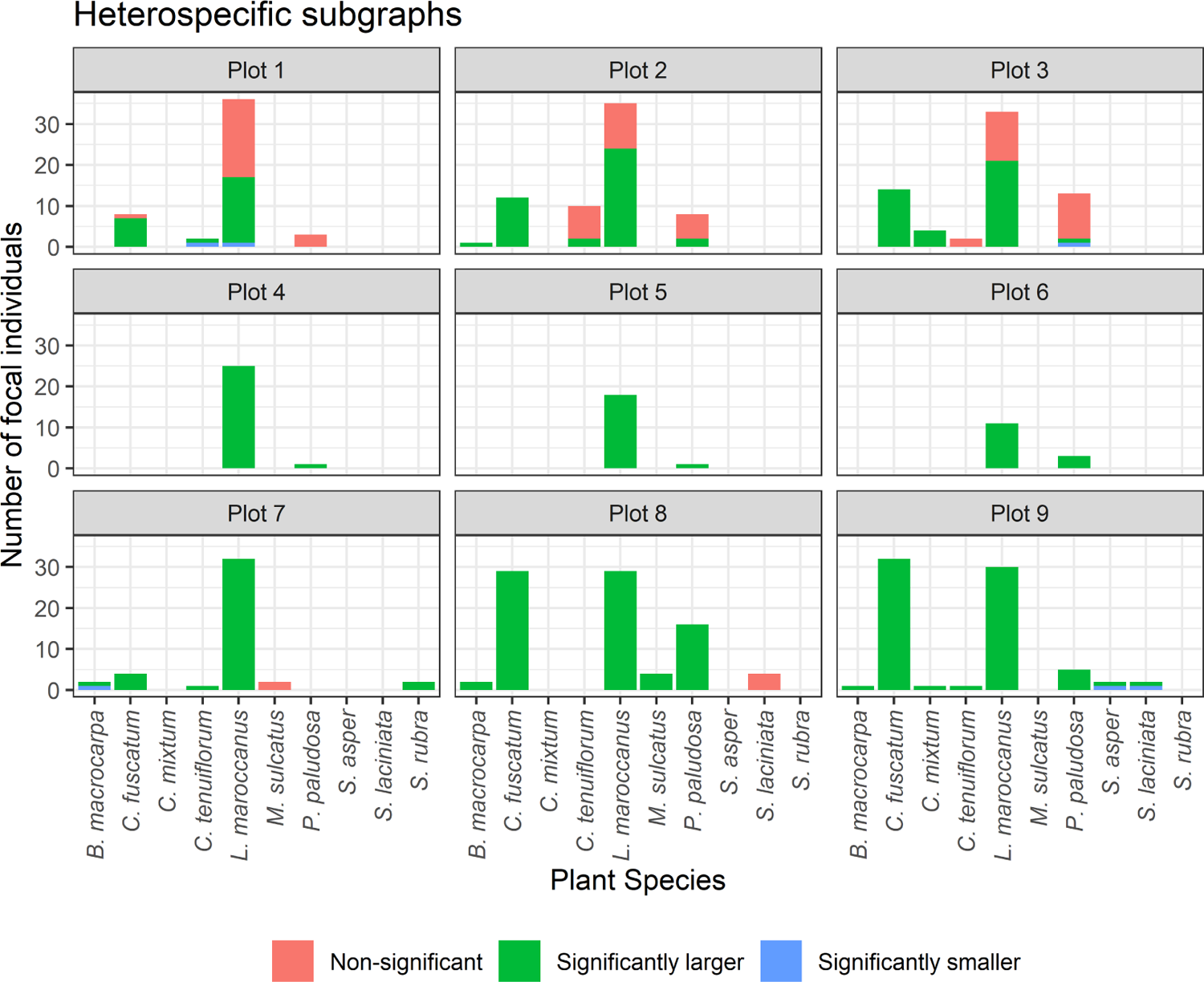
Number of plant individuals with significant and non-significant heterospecific motifs per plot and plant species, when the null model 1 in appendix 3 is considered.

## 17 GLMMs results for seed production and their results

The model for each plant species can be fully specified as follows:

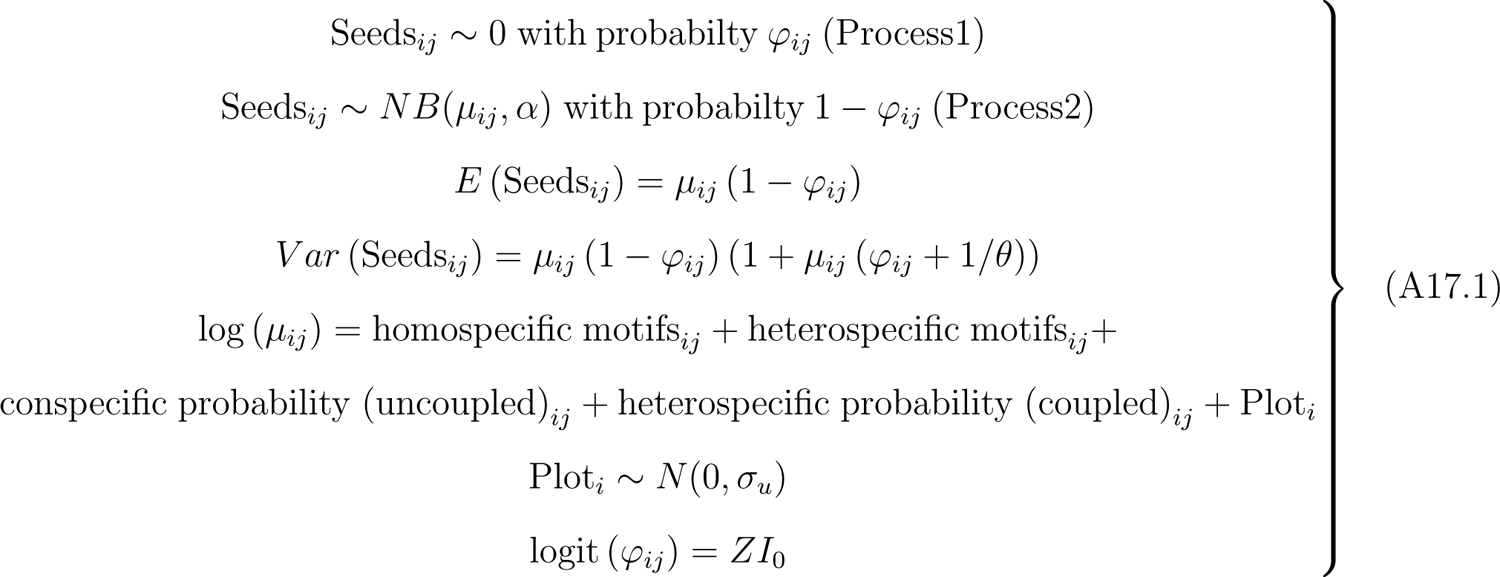

where Seeds*_ij_* is the seed production per fruit of the *j*th individual in Plot *i* for *i* = 1*, · · ·,* 9; Plot*_i_* is the random intercept, which is assumed to be normally distributed with mean 0 and variance *σ*^2^; and represents the overdispersion parameter of the negative binomial. In addition, notice that, for each observation, there are two possible data generation processes and the result of a Bernoulli trial determines which process is used. For observation *ij*, Process 1 is selected with probability *φ_ij_* and Process 2 with probability 1 *− φ_ij_*, where *φ_ij_* is constant and depends on the constant term *ZI*_0_. Process 1 generates only zero counts, whereas Process 2 generates counts from a negative binomial model.

**Table A17.1:**
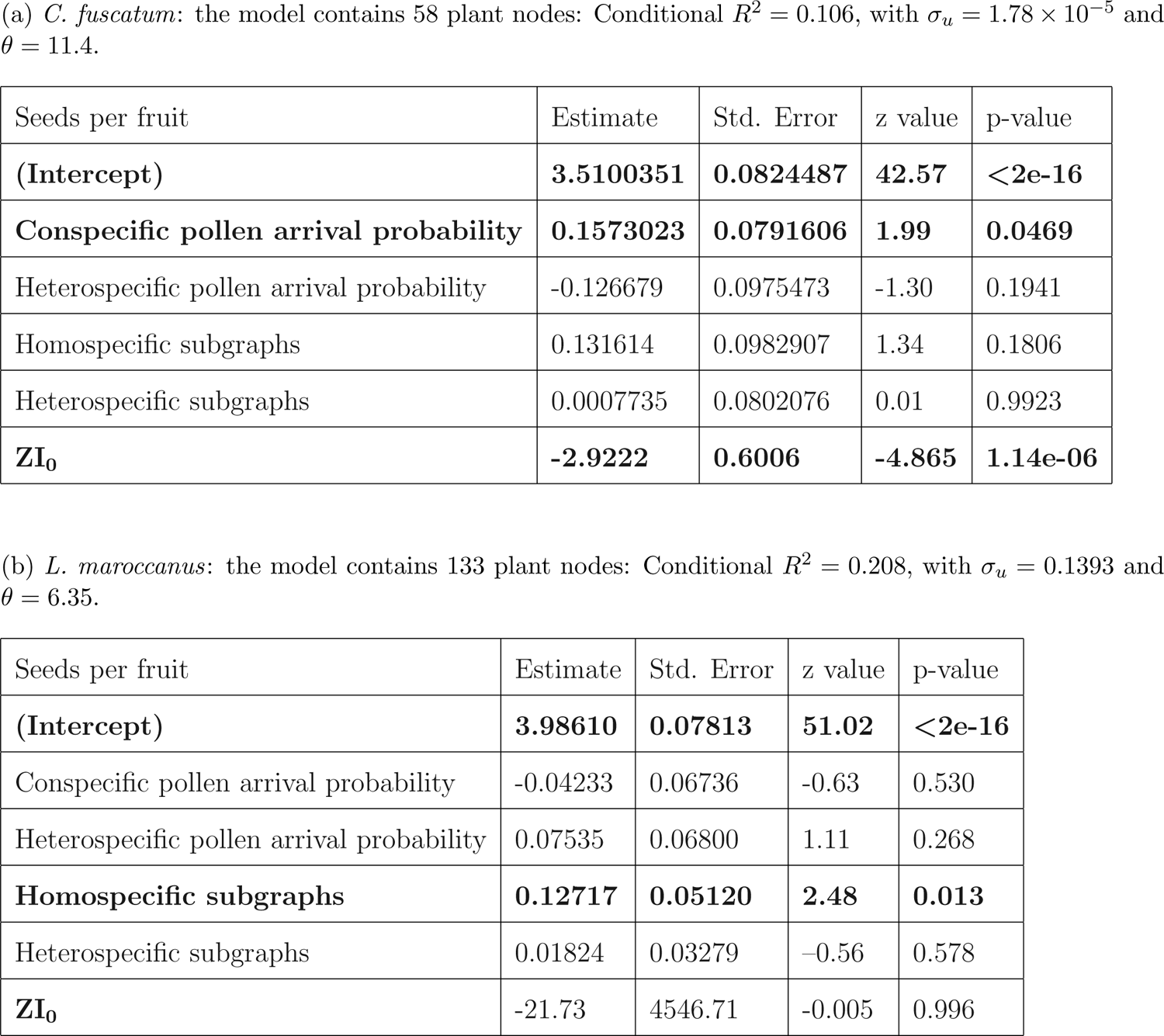
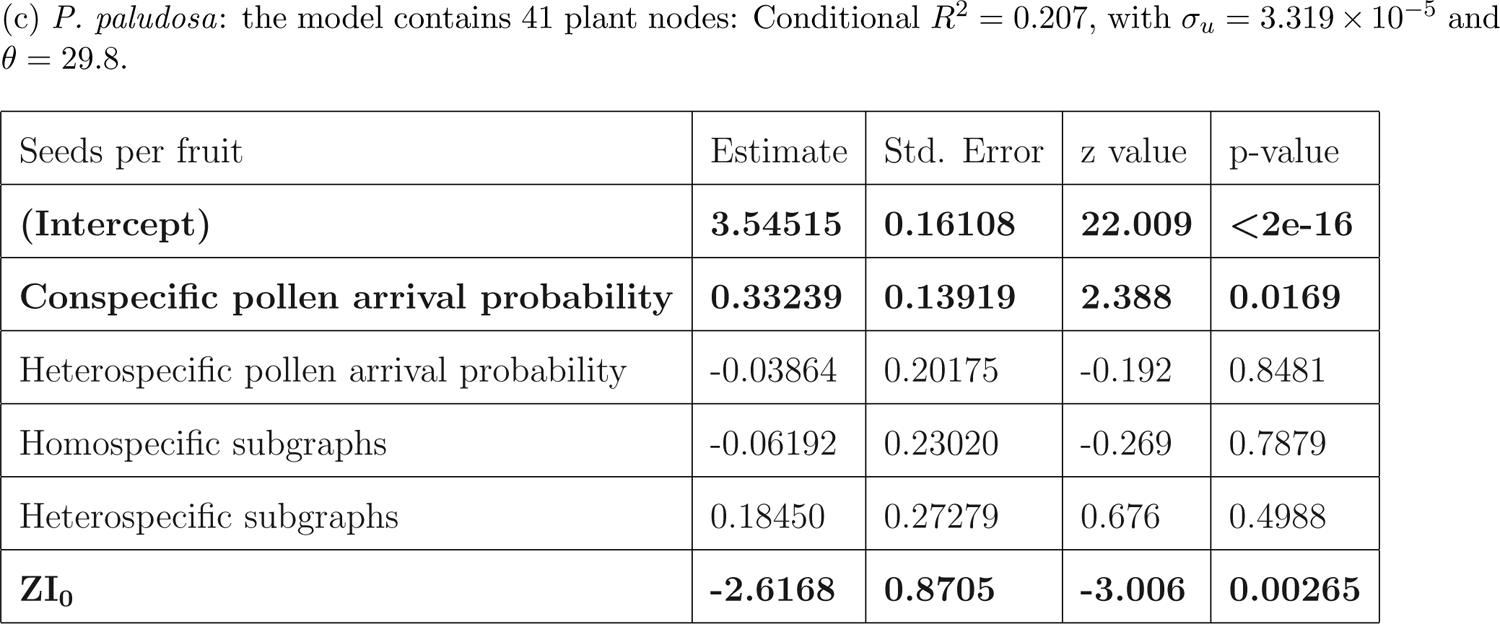
Results for the GLMMs of *C. fuscatum* (a), *L. maroccanus* (b), *P. paludosa* (c) showing the effect on plant reproduction (seeds per fruit). The term *ZI*_0_ describes the zero inflation probability *φ_ij_*, given by logit (*φ_ij_*) = *ZI*_0_ (see Eq. A17.1). Bold characters indicate variables with p-value *<* 0.05.

| Seeds per fruit | Estimate | Std. Error | z value | p-value |
| --- | --- | --- | --- | --- |
| **(Intercept)** | **3.5100351** | **0.0824487** | **42.57** | **<2e-16** |
| **Conspecific pollen arrival probability** | **0.1573023** | **0.0791606** | **1.99** | **0.0469** |
| Heterospecific pollen arrival probability | -0.126679 | 0.0975473 | -1.30 | 0.1941 |
| Homospecific subgraphs | 0.131614 | 0.0982907 | 1.34 | 0.1806 |
| Heterospecific subgraphs | 0.0007735 | 0.0802076 | 0.01 | 0.9923 |
| **ZI<sub>0</sub>** | **-2.9222** | **0.6006** | **-4.865** | **1.14e-06** |

| Seeds per fruit | Estimate | Std. Error | z value | p-value |
| --- | --- | --- | --- | --- |
| **(Intercept)** | **3.98610** | **0.07813** | **51.02** | **<2e-16** |
| Conspecific pollen arrival probability | -0.04233 | 0.06736 | -0.63 | 0.530 |
| Heterospecific pollen arrival probability | 0.07535 | 0.06800 | 1.11 | 0.268 |
| **Homospecific subgraphs** | **0.12717** | **0.05120** | **2.48** | **0.013** |
| Heterospecific subgraphs | 0.01824 | 0.03279 | -0.56 | 0.578 |
| **ZI<sub>0</sub>** | -21.73 | 4546.71 | -0.005 | 0.996 |
| **(Intercept)** | **3.54515** | **0.16108** | **22.009** | **<2e-16** |
| **Conspecific pollen arrival probability** | **0.33239** | **0.13919** | **2.388** | **0.0169** |
| Heterospecific pollen arrival probability | -0.03864 | 0.20175 | -0.192 | 0.8481 |
| Homospecific subgraphs | -0.06192 | 0.23020 | -0.269 | 0.7879 |
| Heterospecific subgraphs | 0.18450 | 0.27279 | 0.676 | 0.4988 |
| **ZI<sub>0</sub>** | **-2.6168** | **0.8705** | **-3.006** | **0.00265** |

## 18 Alternative GLMMs with the covariate “visits”

**Table A18.1:**
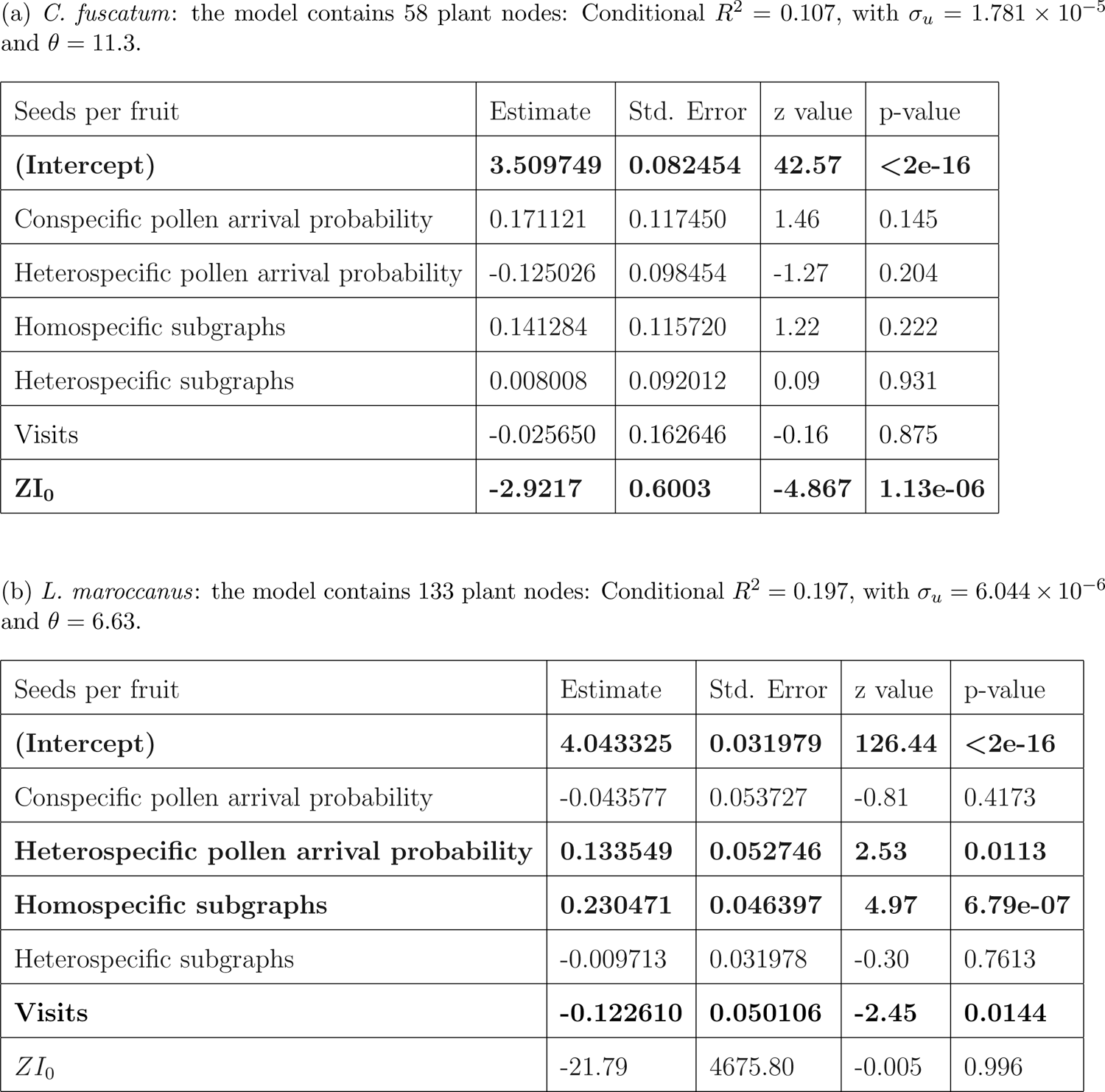
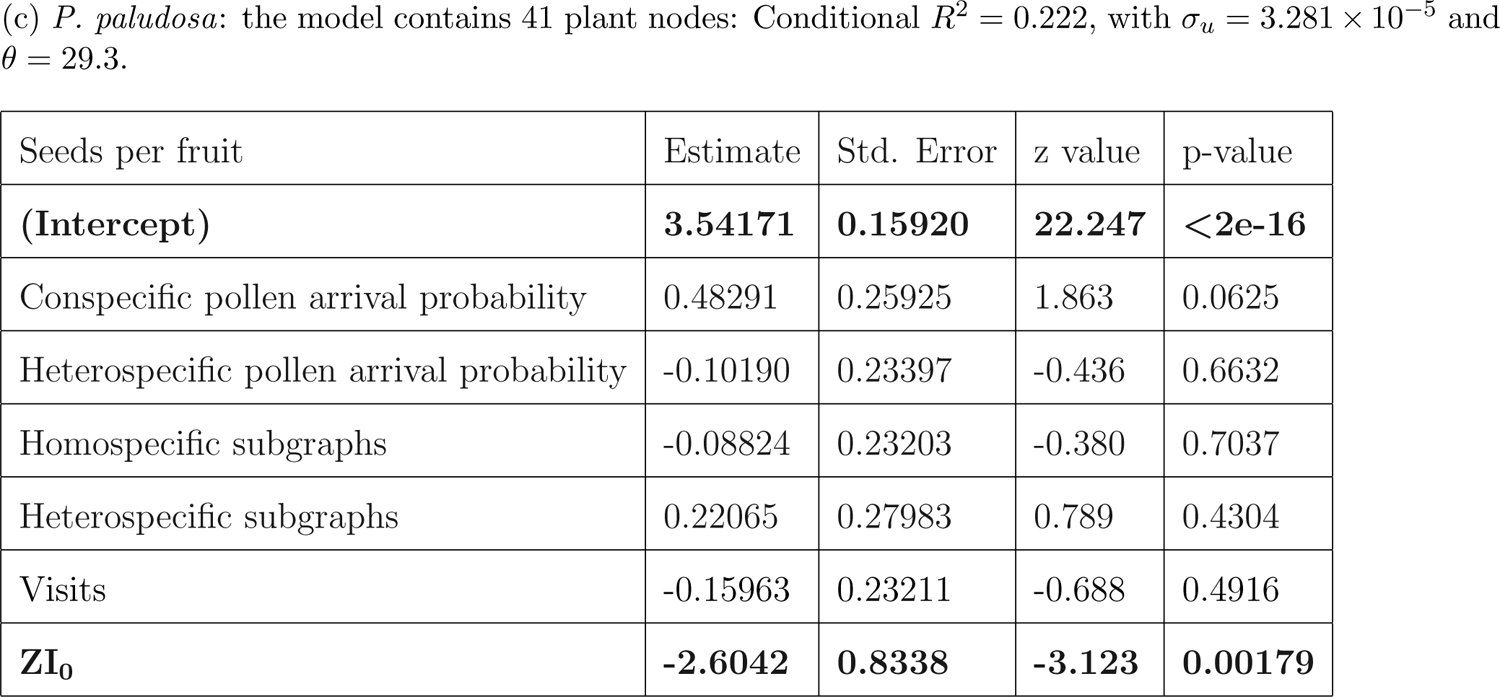
Results for the GLMMs of *C. fuscatum* (a), *L. maroccanus* (b), *P. paludosa* (c) showing the effect on plant reproduction (seeds per fruit). The term *ZI*_0_ describes the zero inflation probability *φ_ij_*, given by logit (*φ_ij_*) = *ZI*_0_ (see Eq. A17.1). Bold characters indicate variables with p-value *<* 0.05.

| Seeds per fruit | Estimate | Std. Error | z value | p-value |
| --- | --- | --- | --- | --- |
| **(Intercept)** | **3.509749** | **0.082454** | **42.57** | **<2e-16** |
| Conspecific pollen arrival probability | 0.171121 | 0.117450 | 1.46 | 0.145 |
| Heterospecific pollen arrival probability | -0.125026 | 0.098454 | -1.27 | 0.204 |
| Homospecific subgraphs | 0.141284 | 0.115720 | 1.22 | 0.222 |
| Heterospecific subgraphs | 0.008008 | 0.092012 | 0.09 | 0.931 |
| Visits | -0.025650 | 0.162646 | -0.16 | 0.875 |
| **$ZI_0$** | **-2.9217** | **0.6003** | **-4.867** | **1.13e-06** |

| Seeds per fruit | Estimate | Std. Error | z value | p-value |
| --- | --- | --- | --- | --- |
| **(Intercept)** | **4.043325** | **0.031979** | **126.44** | **<2e-16** |
| Conspecific pollen arrival probability | -0.043577 | 0.053727 | -0.81 | 0.4173 |
| **Heterospecific pollen arrival probability** | **0.133549** | **0.052746** | **2.53** | **0.0113** |
| **Homospecific subgraphs** | **0.230471** | **0.046397** | **4.97** | **6.79e-07** |
| Heterospecific subgraphs | -0.009713 | 0.031978 | -0.30 | 0.7613 |
| **Visits** | **-0.122610** | **0.050106** | **-2.45** | **0.0144** |
| $ZI_0$ | -21.79 | 4675.80 | -0.005 | 0.996 |
| **(Intercept)** | **3.54171** | **0.15920** | **22.247** | **<2e-16** |
| Conspecific pollen arrival probability | 0.48291 | 0.25925 | 1.863 | 0.0625 |
| Heterospecific pollen arrival probability | -0.10190 | 0.23397 | -0.436 | 0.6632 |
| Homospecific subgraphs | -0.08824 | 0.23203 | -0.380 | 0.7037 |
| Heterospecific subgraphs | 0.22065 | 0.27983 | 0.789 | 0.4304 |
| Visits | -0.15963 | 0.23211 | -0.688 | 0.4916 |
| **ZI<sub>0</sub>** | **-2.6042** | **0.8338** | **-3.123** | **0.00179** |

**Fig. A18.1:**
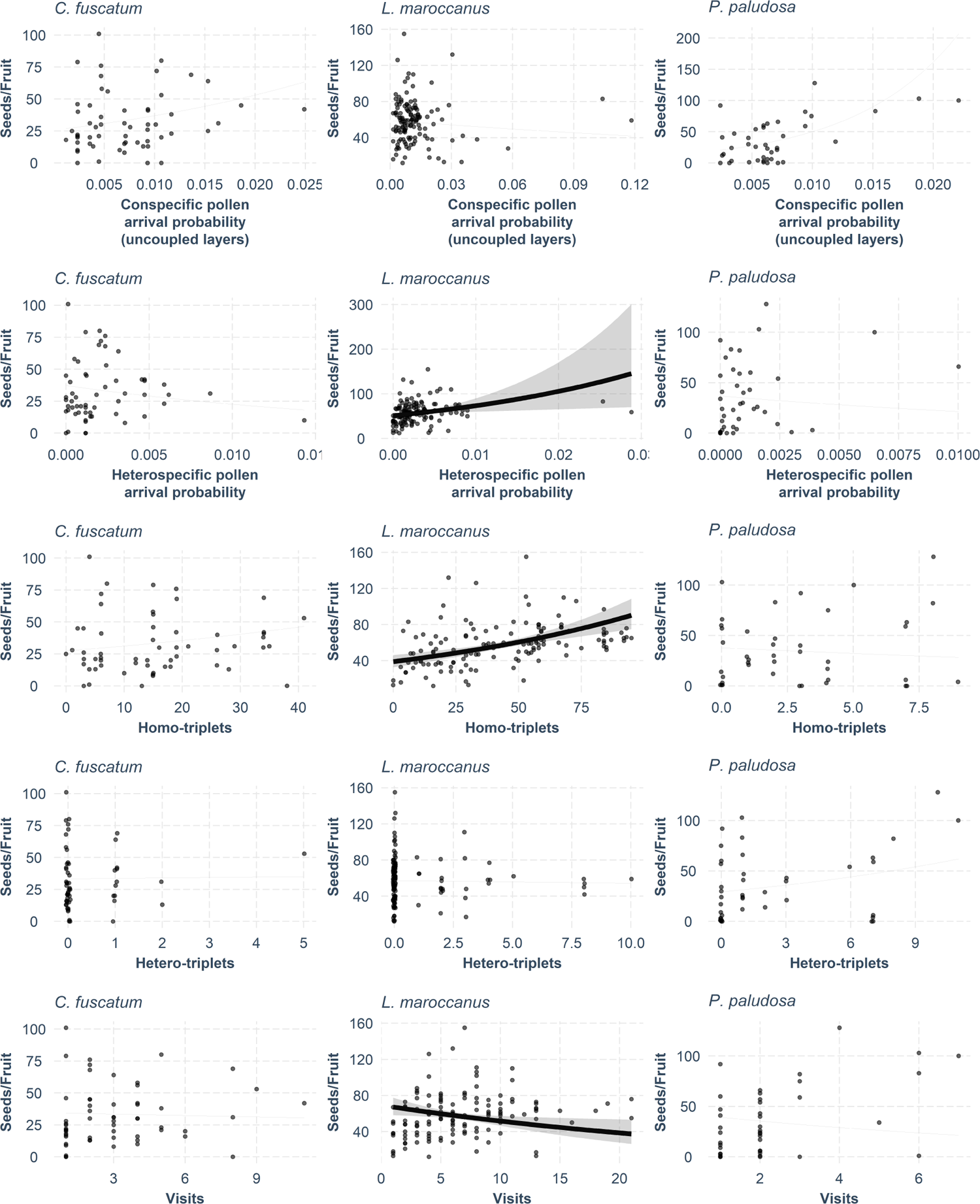
Results for plant’s explanatory variables in the seed production GLMMs (R-package jtools v2.1.4 (Long, 2020)): Conspecific pollen arrival probability (first row), heterospecific pollen arrival probability (second row), homospecific subgraphs (third row), and heterospecific subgraphs (fourth row) and visits (fifth row). In panels with significant explanatory variables (p-value *<* 0.05), we represent (i) the expected values of explanatory variables (black line), (ii) the confidence interval for the expected values (gray band), and (iii) dependence of seed production on the explanatory variable (dark gray dots); otherwise, only the latter is shown.

